# Dynamic recruitment of PV interneurons in the somatosensory cortex induced by experience-dependent plasticity

**DOI:** 10.1101/2025.06.27.662081

**Authors:** Baruc Campos, Xinruo Yang, Iris Marmouset-de la Taille, Mario Dipoppa, William Zeiger

## Abstract

Adaptive circuit plasticity plays crucial roles in the brain during development, learning, sensory experience, and after injury. During chronic whisker trimming, a well-studied paradigm for inducing experience-dependent plasticity, whisker representations in the somatosensory barrel cortex (S1BF) undergo remapping, with expansion of maps for spared whiskers and contraction of maps for trimmed whiskers. At the cellular level, excitatory pyramidal (Pyr) cells in Layer 2/3 shift their whisker tuning, increasing selectivity to spared whiskers and away from deprived whiskers. While these changes are well documented, the circuit mechanisms regulating experience-dependent plasticity remain incompletely characterized. Parvalbumin (PV) interneurons play important roles in regulating the spatial and temporal dynamics of sensory evoked activity in Pyr cells and have been implicated in the regulation of experience-dependent plasticity in other cortical regions. However, there is little evidence as to how the sensory evoked activity of PV cells changes in S1BF during whisker trimming or how those changes might affect cortical remapping. To address these questions, we used longitudinal in vivo two-photon (2P) calcium imaging of PV cells in S1BF before, during, and after inducing experience-dependent plasticity by whisker trimming. At baseline, we found that PV cells have spatially distributed responses to whisker deflections, responding best to the principal whisker of a given barrel and less frequently to surrounding whiskers in a distance-dependent manner. After whisker trimming, there is a substantial recruitment of PV cells responsive to the spared whisker in deprived, but not spared, barrels. Upon whisker regrowth, this recruitment is reversed, but changes in individual PV cell whisker selectivity can persist for weeks. To better understand how changes in PV activity might affect sensory-evoked responses in S1BF, we used chemogenetics to acutely manipulate the activity of PV cells and found that modulating PV cell activity strongly affects sensory-evoked responses in local Pyr and PV cells, as well as Somatostatin (SST) interneurons. Surprisingly, activating PV cell activity strongly suppressed sensory-evoked activity in all three cell types. Computational modeling of the effects of chemogenetic perturbation of PV cells reproduced observed effects on the S1BF microcircuit, underscoring that directly perturbing interneurons in one direction can paradoxically change their activity in the opposite direction due to network effects. Together, our results reveal dynamic changes in the spatial distribution and tuning of PV cells during experience-dependent plasticity and demonstrate that increased PV cell activity can profoundly alter microcircuit dynamics in the adult S1BF.

## Introduction

Cortical circuits have the capacity to adapt and change through a process broadly termed plasticity. Cortical circuit plasticity can occur in many contexts, including development, learning, disease, and in response to sensory experience^1–4^. One of the best studied models for sensory experience-dependent plasticity is the rodent whisker somatosensory system. Mice rely on whiskers as a primary sensory input and the whisker somatosensory area (S1BF) is one of the largest regions of the mouse cortex^5^. The S1BF has a strong somatotopic organization^6^. Thalamocortical inputs from the ventroposteriomedial nucleus (VPM) conveying information from individual whiskers project primarily to Layer 4 (L4) of corresponding columns, or “barrels”^7,8^. From L4, activity propagates vertically to supra- and infra-granular layers and then horizontally to adjacent barrels^9–11^. Within individual barrels, a plurality of excitatory pyramidal (Pyr) neurons respond best to deflection of the barrel’s corresponding principal whisker, but responses are overall very heterogenous, particularly in L2/3, with many Pyr cells responding best to surround whiskers^12,13^. Sensory deprivation via chronic whisker trimming is a classical paradigm for inducing experience-dependent plasticity in the S1BF^14^. Whisker trimming leads to the expansion of the cortical representation of “spared” un-trimmed whiskers and the contraction of the cortical representation of “deprived” trimmed whiskers^15–18^. At the level of individual neurons, excitatory Pyr cells in L2/3 undergo dramatic changes in their whisker tuning, with shifts in selectivity for spared whiskers over trimmed whiskers in some neurons and recruitment of additional neurons becoming newly responsive to the spared whisker^19^. These changes likely arise from both Hebbian and homeostatic synaptic plasticity mechanisms^20,21^, but the precise circuit alterations regulating experience-dependent plasticity remain incompletely characterized.

Evidence suggests that parvalbumin (PV) interneurons may regulate experience-dependent plasticity. PV cells are the most common cortical interneuron class, comprising ∼40% of all cortical interneurons, depending on the cortical region and layer studied^22–25^. They exhibit key electrophysiological properties including short action potential duration, fast afterhyperpolarizations, and high frequency spiking with minimal adaptation^26–28^. PV cells synapse primarily onto peri-somatic regions of excitatory neurons^29–31^. Receiving both strong bottom-up inputs^32,33^ as well as intracortical inputs^34–36^, most PV cells in S1BF have broad receptive fields with responses to stimulation of multiple whiskers^32,33,37,38^. Such connectivity results in PV cells contributing strongly to both feed-forward and feedback inhibition. For example, in L4 PV cells receive strong thalamic input at short latency, positioning them to provide robust and fast feedforward inhibition to excitatory cells in response to whisker deflections^32,33,39^. Functionally, this allows PV cells to sharpen the spatial and temporal precision of activity in excitatory cells^40–45^, leading to sparse firing and contributing to gain modulation^46^. PV cells also generate synchronous activity that regulates cortical oscillations^47–49^.

Several lines of evidence suggest an important role for PV cells during cortical circuit plasticity^50^. In development, maturation of PV cells corresponds with the closure of critical periods in the visual cortex^51,52^. If sensory experience is disrupted, as in the case of monocular deprivation, PV cells undergo dynamic changes in responsivity distinct from those of Pyr cells, first becoming more responsive to the occluded eye before switching and preferentially responding to the spared, open eye^53^. In adult animals, chemogenetic inhibitory modulation of PV cells potentiates experience-dependent plasticity in both primary visual (V1)^54,55^ and primary auditory (A1) cortex^56^. Despite being a site of robust experience-dependent plasticity, the precise role of PV cells in regulating S1BF circuit plasticity remains incompletely understood. Acutely after whisker trimming the intrinsic excitability of PV cells is reduced in deprived barrels, leading to disinhibition of Pyr cells and temporary preservation of Pyr cell firing rates^20,57,58^. However, how PV cell activity and whisker selectivity change over longer time scales and in spared barrels is unknown.

Here, we sought to address these gaps in knowledge using acute and longitudinal two-photon (2P) in vivo imaging of genetically defined PV cells in L2/3 of S1BF before, during, and after chronic whisker trimming. Before trimming, PV cells in S1BF exhibit spatially distributed responses to whisker deflections, with whisker receptive fields similar to those of Pyr cells. During whisker trimming, PV cells undergo barrel-specific changes in whisker selectivity, with strong recruitment of PV cells responsive to the spared whisker in deprived, but not spared, barrels. On the other hand, after whisker regrowth, baseline whisker selectivity of PV interneurons is largely re-established in deprived barrels, but there are long-lasting shifts in responsivity toward the spared whisker in the spared barrel. Acute modulation of PV cells with DREADDs (<u>D</u>esigner <u>R</u>eceptors <u>E</u>xclusively <u>A</u>ctivated by <u>D</u>esigner <u>D</u>rugs) leads to changes in the activity of local Pyr, PV, and somatostatin (SST) cells. In particular, chemogenetic activation of PV cells leads to strong suppression of sensory-evoked activity in local Pyr and SST cells, as well as a paradoxical suppression of sensory-evoked activity in local PV cells themselves. Computational modeling of these DREADD-mediated perturbations broadly reproduced these effects, showing that the response to a PV manipulation depends on the circuit’s baseline state and recurrent dynamics and cannot be inferred from the direct action of the DREADD alone. These results highlight the complexity of responses to changes in PV cell activity that occur in recurrent cortical circuits and underscore the need to carefully interpret results from chemogenetic manipulations in S1BF. Our results provide an important characterization of the spatial distribution of whisker selectivity in PV cells at baseline, how these responses change during and after sensory experience, and demonstrate how PV-mediated inhibition regulates sensory evoked responses across cell types of the S1BF microcircuit.

## Methods

### Materials

DREADD Agonist Compound 21 (C21) was obtained from HelloBio. Recombinant adeno-associated viruses (AAVs), including AAV1-syn-jGCaMP8s-WPRE^59^ (gift from GENIE Project, Addgene viral prep #162374-AAV1), AAV8-Ef1a-Coff/Fon-GCaMP6f^60^ (gift from Karl Deisseroth & INTRSECT 2.0 Project, Addgene viral prep #137124-AAV8), AAV9-hSyn-DIO-mCherry (gift from Bryan Roth, Addgene viral prep #50459-AAV9), AAV9-hSyn-DIO-hM4D(Gi)-mCherry^61^ (gift from Bryan Roth, Addgene viral prep #44362-AAV9), and AAV5-hSyn-DIO-hM3D(Gq)-mCherry^61^ (gift from Bryan Roth, Addgene viral prep #44361-AAV5) were obtained from Addgene.

### Experimental Animals

All experiments followed the U.S. National Institutes of Health guidelines for animal research, under an animal use protocol approved by the University of California Los Angeles Animal Research Committee (ARC). Male and female mice were used, beginning at 7-10 weeks old at the time of cranial window surgery. All animals were housed in a vivarium with a 12 h light/dark cycle. For these experiments we used 68 heterozygous PV-Cre mice (B6.129P2-Pvalbtm1(cre)Arbr/J, Jax line 017320)^62^, 28 double transgenic PV-Cre:Ai162 mice (Ai162(TIT2L-GC6s-ICL-tTA2)-D, Jax line 031562)^63^, and 42 double transgenic PV-Cre:Sst-ires-Flp (B6J.Cg-Ssttm3.1(flpo)Zjh/AreckJ, Jax line 031629) mice. All transgenic lines were maintained on a C57BL/J6 background.

### Cranial Window Surgery

Cranial window implantation was performed according to previously published protocols^64,65^. Mice were deeply anesthetized with 5% isoflurane, with 1.5-2% isoflurane for maintenance. After removal of the scalp and periosteum, an ∼4 mm diameter circular craniotomy, centered over the primary somatosensory cortex (∼3 mm lateral to the midline and ∼2 mm caudal to Bregma) was made using a pneumatic dental drill with a FG ¼ drill bit (Midwest Dental). For mice requiring injection of AAVs, AAVs were injected at a concentration from 1.8 to 2.1 x 10^13 GC/mL and volume of 75 nL directly into 4 sites in the cortex using a glass capillary nanoinjector (Neurostar). The craniotomy was sealed using either a single 5 mm #1 sterile glass coverslip (Harvard Apparatus), or a 4 mm coverslip glued to a 5 mm coverslip using an optical adhesive (Norland Products, #71) which was glued to the skull with cyanoacrylate glue (Krazy Glue) and dental acrylic (OrthoJet, Lang Dental). A stainless steel headbar for head fixation was embedded in dental acrylic. Carprofen (5 mg/kg, i.p., Zoetis) and dexamethasone (0.2 mg/kg, i.p., Vet One) were provided for pain relief and mitigation of edema on the day of surgery and daily for the next 48 h. Mice were allowed to recover from the surgery for at least 3 weeks before the first imaging session.

### Chronic Whisker Trimming Protocol

Imaging was performed in two barrels at baseline in 10 week old mice: 1) The C1 barrel stimulating its PW C1 and the SWs D1, B2 and E3 (**Fig. 1**); and 2) The D1 barrel stimulating its PW D1 and the SW C1 (**Fig. S2**). To ensure whisker stimuli deflected only whiskers of interest, the whiskers directly adjacent to those targeted for stimulation on the right side of the snout were trimmed using fine scissors to a length of ∼10 mm immediately prior to baseline imaging. All whiskers on the right side of the face except the spared D1 whisker were trimmed flush with the vibrissal pad immediately following baseline imaging and re-trimmed every other day to remove any whisker re-growth, for a total of 18 days. To facilitate whisker trimming, mice were anesthetized with isoflurane (5% for induction) and maintained at 1.5-2% isoflurane for ∼ 1 min during trimming. At days 3, 6, 12 and 18, imaging was performed in both the deprived C1 barrel and the spared D1 barrel stimulating the spared D1 whisker. After the imaging session on day 18, whiskers were allowed to regrow for 4 weeks. On day 46, imaging was performed in the C1 and D1 barrels stimulating the same whiskers as at baseline. Sham trimming was performed on the control untrimmed mice (**Fig. S4a-b**). Mice were handled and anesthetized the same as the trimmed mice, but instead of trimming, all the whiskers from the pad were softly deflected with the scissors in a rostral/caudal direction five times. Imaging in this untrimmed group was performed in the C1 and D1 barrels at baseline, day 18, and day 46. A separate cohort of mice were used to evaluate potential changes in the length of whiskers pre- and post-trimming (**Fig. S4c**). These mice were handled, anesthetized, and whisker trimmed in the same way as cohort of trimmed mice that underwent imaging.

**Figure 1.**
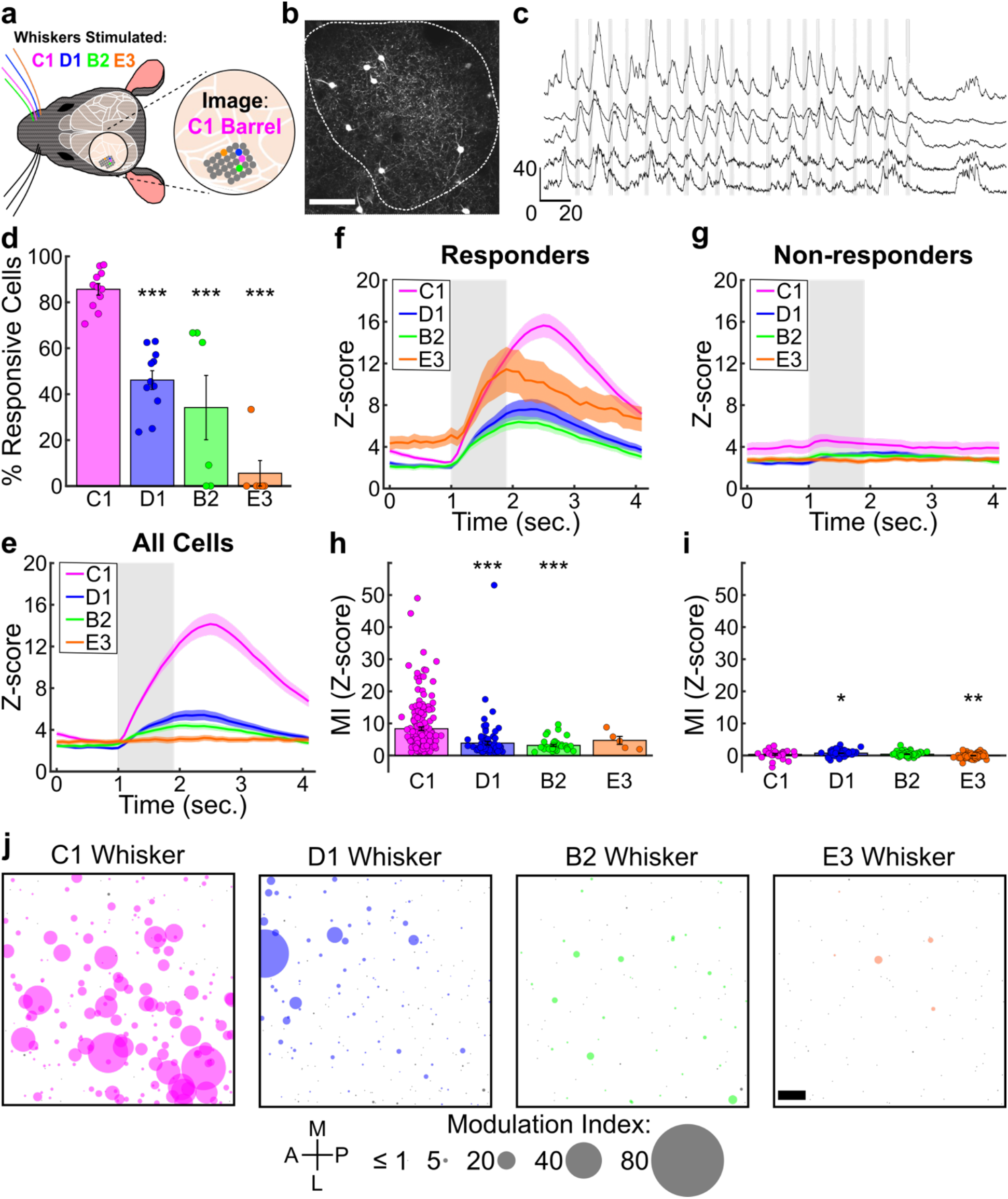
The spatial distribution of whisker-evoked responses of PV cells in the S1BF. **a.** Schematic of the imaging FOV and stimulated whiskers (PW C1, magenta; and SWs D1, blue; B2, green; E3, orange). **b.** Average projection image of a representative FOV of PV cells expressing GCaMP6s. Dotted outline corresponds to the functional IOSI map of the C1 barrel. Scale bar=100 µm. **c.** Z-scored calcium fluorescence traces from PV cells during whisker stimulation. Gray bars indicate epochs of whisker deflections (10 Hz, 1 s). Scale Y-axis, Z-score; X-axis, time (seconds). **d.** Percentage of whisker responsive cells to the indicated whiskers (n=11 mice for PW C1 and SW D1; n=6 mice for SWs B2 and E3). GLME binomial model, fixed effect of whisker, with significance for individual coefficients compared to C1, corrected using Benjamini and Hochberg’s method, indicated over corresponding data points (*, *p*<0.05; **, *p*<0.01; ***, *p*<0.001). **e-g.** Mean evoked calcium traces for all cells (**e**, n=210 cells for C1 and D1, n=100 cells for B2 and E3), responders (**f**, n=183 cells for C1, 101 cells for D1, 39 cells for B2, and 5 cells for E3), and non-responders (**g**, n=27 cells for C1, 109 cells for D1, 61 cells for B2, and 95 cells for E3). **h-i.** Modulation index (MI) calculated for all responders (**h**, n=183 cells for C1, 101 cells for D1, 39 cells for B2, and 5 cells for E3) and non-responders (**i**, n=27 cells for C1, 109 cells for D1, 61 cells for B2, and 95 cells for E3) by whisker stimulated. LME model, fixed effect of whisker. Significance for individual coefficients compared to C1, corrected using Benjamini and Hochberg’s method, are indicated over corresponding data points (*, *p*<0.05; **, *p*<0.01; ***, *p*<0.001). **j.** Spatial distribution of all PV cells plotted according to relative position in the FOV centered on the C1 barrel. Responders are colored according to the indicated whisker, non-responders are colored in gray. Size of the circle corresponds to the MI of the cell for the indicated whisker. Scale bar=50 µm.

**Figure 2.**
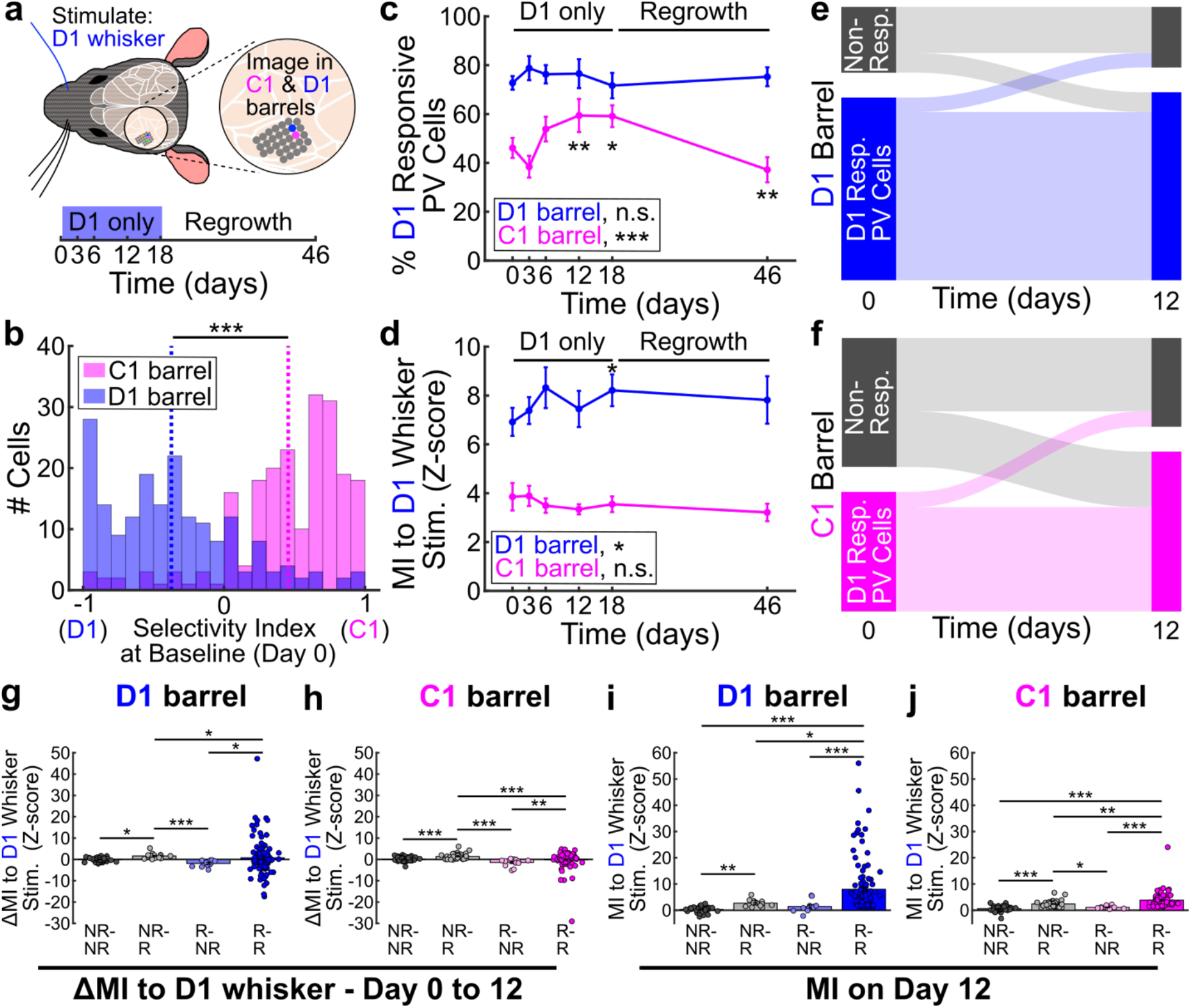
PV cell responses to the spared whisker are increased in the deprived barrel during whisker trimming. **a.** Schematic of the imaging FOV (C1 and D1 barrels), stimulated whisker (D1), and trimming timeline. **b.** Histogram of the selectivity index of cells in the C1 (magenta) or D1 (blue) barrel at baseline (n=189 cells for D1 and 210 cells for C1). LME model, fixed effect of barrel (***, *p*<0.001). **c.** Percentage of SW D1 responsive PV cells in the deprived C1 barrel (magenta) or percentage of PW D1 responsive PV cells in the spared D1 barrel (blue) over time (n=11 mice). GLME binomial model for the spared D1 barrel, fixed effect of timepoint (*p*=0.41). GLME binomial model for the deprived C1 barrel, fixed effect of timepoint (*p*<0.001). Significance for individual coefficients compared to day 0, corrected using Benjamini and Hochberg’s method, are indicated over corresponding data points (*, *p*<0.05; **, *p*<0.01). **d.** MI of SW D1 responsive PV cells in the deprived C1 barrel (magenta) or PW D1 responsive PV cells in the spared D1 barrel (blue) over time (n=139/101, 154/83, 149/116, 143/135, 141/126, and 141/70 at day 0, 3, 6, 12, 18, and 46 for the D1/C1 barrels, respectively). LME for the spared D1 barrel, fixed effect of timepoint (*p*=0.03). LME for the deprived C1 barrel, fixed effect of timepoint (*p*=0.36). Significance for individual coefficients compared to day 0, corrected using Benjamini and Hochberg’s method, are indicated over corresponding data points (*, *p*<0.05). **e-f.** Sankey diagrams showing responsiveness of individual PV cells to the PW D1 in the spared D1 barrel (**e**, n=189 cells) or SW D1 in the deprived C1 barrel (**f**, n=210 cells) between days 0 and 12. **g.** Change in MI of PV cells in the spared D1 barrel to the PW D1 between day 0 and day 12. Cells are grouped by switch category: non-responsive to the PW D1 at day 0 and day 12 (NR-NR, n=35); switched from non-responsive to responsive (NR-R, n=15); switched from responsive to non-responsive (R-NR, n=11); or responsive on both day 0 and day 12 (R-R, n=128). Kruskal-Wallis test with tukey-kramer correction for multiple comparisons (*, *p*<0.05; ***, *p*<0.001). **h.** Change in MI of PV cells in the deprived C1 barrel to the SW D1 between day 0 and day 12, by switch category (n=62, 47, 13, and 88 cells for NR-NR, NR-R, R-NR, and R-R, respectively). Kruskal-Wallis test with tukey-kramer correction for multiple comparisons (**, *p*<0.01; ***, *p*<0.001). **i.** MI of PV cells in the spared D1 barrel to the PW D1 at day 12, by switch category (n=35, 15, 11, and 128 cells for NR-NR, NR-R, R-NR, and R-R, respectively). Kruskal-Wallis test with tukey-kramer correction for multiple comparisons (*, *p*<0.05; **, *p*<0.01; ***, *p*<0.001). **j.** MI of PV cells in the deprived C1 barrel to the SW D1 at day 12, by switch category (n=62, 47, 13, and 88 cells for NR-NR, NR-R, R-NR, and R-R, respectively). Kruskal-Wallis test with tukey-kramer correction for multiple comparisons Kruskal-Wallis test with tukey-kramer correction for multiple comparisons (*, *p*<0.05; **, *p*<0.01; ***, *p*<0.001).

### Intrinsic optical signal imaging (IOSI) Acquisition

IOSI was performed as previously described^18^. Animals were sedated with chlorprothixene (∼3 mg/kg, i.p.), lightly anesthetized with ∼0.5-0.7% isoflurane, and head-fixed. The cortical surface was illuminated with 525 nm light to capture an image of the superficial vasculature. The microscope was then focused 300 μm below the cortical surface and illuminated with 625 nm light to record intrinsic signals. Frames were collected at 10 Hz (100 ms exposure time) using an 8 megapixel CCD camera (Thorlabs, 8051M-USB) during thirty trials of whisker deflections (100 Hz sine wave, 1.5 seconds long). Whisker stimuli were generated in MATLAB, amplified, and delivered using a glass capillary affixed to a piezoelectric bending actuator (Bimitech Python PBA6014-5H200). Cortical whisker maps were generated as previously described^18^. Briefly, stimulus-evoked change in reflectance values (ΔR/R) were calculated and binarized by thresholding for ΔR/R values below a Z-score of -3. Binarized images were then pseudocolored and overlaid onto images of the vasculature. To quantify map area for single whisker-evoked maps, the medfilt2 function in MATLAB was used to apply a median filter with a 3x3 pixel neighborhood size to binarized maps to remove noise and the area of thresholded pixels was calculated. The mean value of all thresholded pixels in the map area was calculated to quantify map intensity.

### Two-photon Calcium Imaging

In vivo calcium imaging was performed using a Thorlabs Bergamo II microscope with fast galvo-resonant scanning mirrors, 14° collection optics with two high-sensitivity GaAsP photomultiplier tubes, and a 16x/0.8 NA objective (Nikon), coupled to an Insight X3 dual output Ti:Sapphire laser (Spectra Physics). For anesthetized imaging, mice were lightly sedated with chlorprothixene (∼3 mg/kg, i.p.) and isoflurane (0.7-0.9%) and kept warm with a heating blanket. For awake imaging, mice were habituated to head-fixation under the microscope in a cylindrical tube prior to imaging. Stimulation of the whiskers (20 stimuli, 1 s duration 10 Hz square wave, with a pseudorandomized 3-6 s interstimulus interval) was delivered to a single whisker by a glass capillary tube affixed to a piezoelectric actuator (Bimitech Python PBA6014-5H200). The glass capillary tube was always positioned ∼7 mm from the follicle, to maintain consistency between mice. The direction of the stimulation was rostral/caudal using square wave stimuli with 0.001 s rise time (amplitude of deflection of 3.35 +/- 0.01mm from peak to peak). We also tested square wave stimuli with 0.01 s rise time (amplitude deflection of 3.07 +/- 0.01mm), or sine wave stimuli with voltage amplitudes of 2, 4, 6, or 8 V (amplitude of deflection of 2.70, 3.50, 4.50 and 5.00 +/- 0.01mm, respectively; **Fig. S1e**). The amplitude of deflection from peak to peak was measured with a digital caliper. Imaging data and whisker stimulation data were synchronized using ThorSync software. Whole field 2x zoom 512 x 512 pixel images (typically 422.16 x 422.16 μm) were acquired with bidirectional scanning at ∼30 Hz, with averaging of 3 frames for a final image acquisition rate of ∼10 Hz. The imaging field-of-view (FOV) was determined according to IOSI maps. For each FOV, ∼150 s of spontaneous activity data was collected prior to imaging whisker-evoked activity. For experiments in which multiple whiskers on an individual mouse were stimulated, imaging was performed serially for each whisker – for example, imaging was performed during stimulation of one whisker, then the whisker affixed to the stimulator was switched and imaging repeated. For experiments involving the use of heterozygous PV-Cre mice injected with AAV1-syn-jGCaMP8s we recorded an average of ∼14 PV cells and ∼123 Pyr cells per FOV. For experiments involving the use of double transgenic PV-Cre:Ai162 mice and PV-Cre:Sst-ires-Flp, we recorded on average ∼19 PV cells and ∼5 SST cells per FOV, respectively.

Motion correction of movies was performed using the motion correction module from *EZcalcium*^66^. Fluorescence traces (ΔF/F) of neuronal calcium transients were extracted using custom-written semi-automated MATLAB routines as previously described^67^ (from PV-Cre:Ai162 and PV-Cre:Sst-ires-Flp double transgenic mice), or using Suite2P^68^ (from PV-Cre mice injected with jGCaMP8s) automated cell segmentation with manual refinement. For Suite2P extracted traces, 0.7 times the neuropil fluorescence signal was subtracted from somatic fluorescence signals. We calculated a modified *Z* score vector from the fluorescence traces for each neuron as previously described^67^. Modified Z-scores for each of the 20 individual whisker stimuli were aligned to the onset of the stimulus and a mean stimulus evoked trace was calculated for each cell. A modulation index (MI) indicating the strength of the whisker-evoked response for each cell was calculated as:

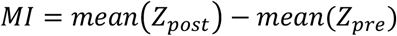

Where Z_post_ is the modified Z-score trace from stimulus onset until 2 seconds later and Z_pre_ is the modified Z-score trace 1 second prior to stimulus onset. For experiments expressing DREADDs in PV cells (**Figs. 4 and 5**), we found the noise of the baseline fluorescence - defined as the standard deviation of the baseline fluorescence (the consecutive 10 seconds with the lowest standard deviation) normalized to the mean of the baseline fluorescence - was strongly modulated by hM3D(Gq) and hM4D(Gi), resulting in Z-scored traces that underestimated the effects of the chemogenetic perturbations. Therefore, we used ΔF/F traces to calculate MI in these experiments. To determine if neurons were responsive to stimulation of a specific whisker we used a probabilistic bootstrapping method to correlate Z-scored calcium traces with epochs of stimulation, shuffling the stimulus trace^67^. Neurons that had significant correlations between whisker stimulation and fluorescence transients (*p* values < 0.01) and had whisker stimulation MI≧1 were considered whisker responsive. We chose this value as we found a small minority of false positive cells were classified as responsive using our bootstrapping correlation method that did not appear to have clear whisker-evoked activity when examining their mean evoked activity traces (**Fig. S1b-d**).

**Figure 3.**
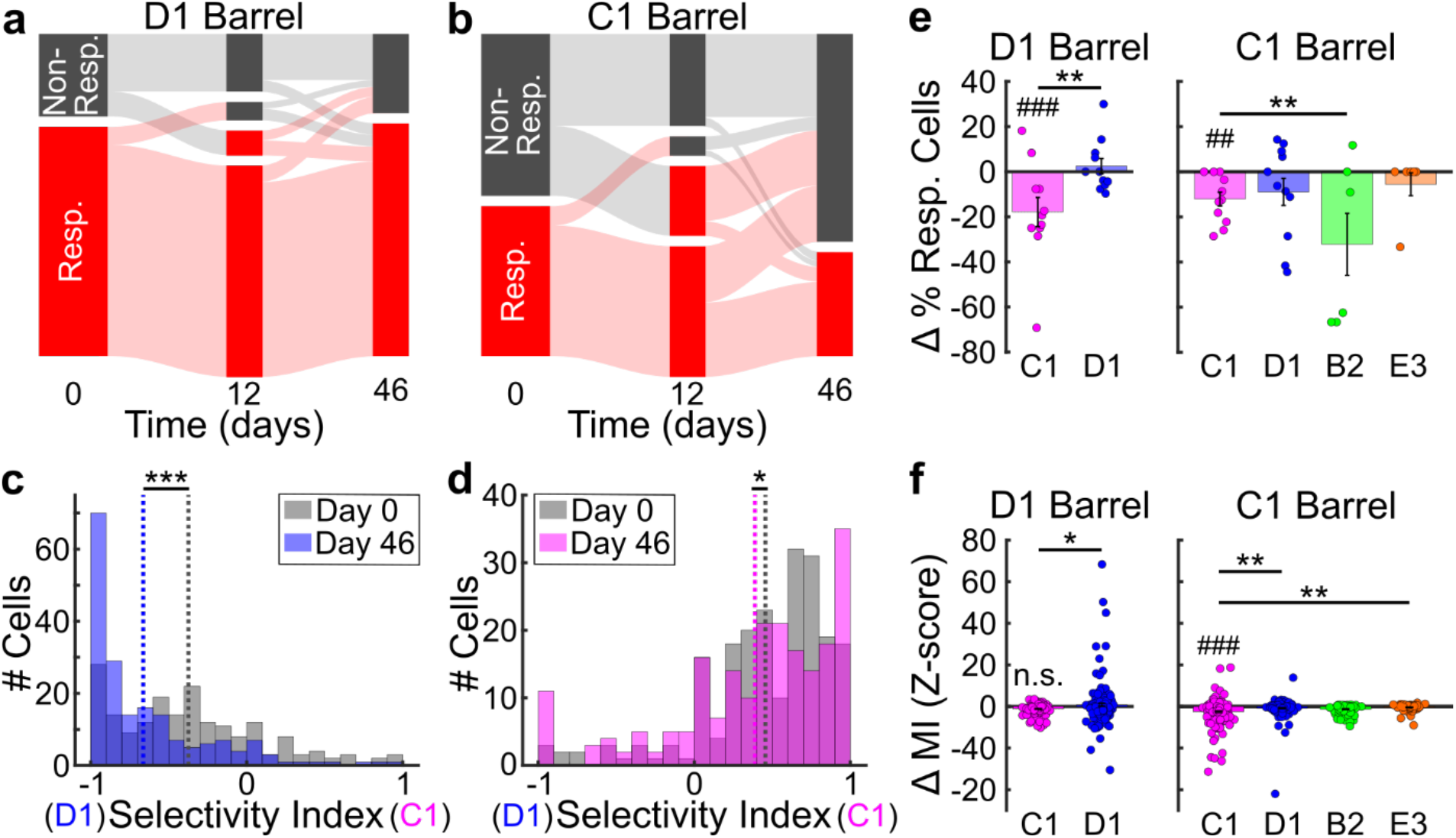
PV cell selectivity shifts toward the spared whisker in the spared barrel persist after whisker regrowth. **a-b**. Sankey diagrams showing responsiveness of individual PV cells to the PW D1 in the spared D1 barrel (**a**) or deprived C1 barrel (**b**) between days 0, 12, and 46. **c-d.** Histogram of the selectivity index of cells in the spared D1 (**c**, n=189 cells) or deprived C1 (**d**, n=210 cells) barrel at day 0 (gray) or day 46 (blue or magenta, respectively). LME model, fixed effect of timepoint (*, *p*<0.05; ***, *p*<0.001). **e.** Change in the percentage of PV cells responsive to the C1 (magenta), D1 (blue), B2 (green), or E3 (orange) whisker in the spared D1 (left) or deprived C1 (right) barrel between day 0 and day 46 (n=11 mice for C1 and D1 whiskers; n=6 mice for B2 and E3 whiskers). D1 barrel, GLME binomial model, ANOVA for fixed effects of whisker (C1 vs. D1 whisker at day 0, *p*<0.001), timepoint (day 0 vs day 46 for C1, ###, *p*<0.001), and whisker*timepoint interaction (**, *p*<0.01). C1 barrel, GLME binomial model, ANOVA for fixed effects of whisker (C1 vs. other whiskers at day 0, *p*<0.001), timepoint (day 0 vs day 46 for C1, ##, *p*<0.01), and whisker*timepoint interaction (**, *p*<0.01), with significance for individual coefficients for whisker*timepoint interactions, corrected using Benjamini and Hochberg’s method, indicated over corresponding data points (**, *p*<0.01). **f.** Change in the MI of PV cells responsive to the C1 (magenta), D1 (blue), B2 (green), or E3 (orange) whisker in the spared D1 (left) or deprived C1 (right) barrel between day 0 and day 46 (D1 barrel: n=189 cells for C1 and D1 whiskers. C1 barrel: n=210 cells for C1 and D1 whiskers; n=100 cells for B2 and E3 whiskers). D1 barrel, GLME binomial model, ANOVA for fixed effects of whisker (C1 vs. D1 whisker at day 0, *p*<0.001), timepoint (day 0 vs day 46 for C1, *p*<0.07), and whisker*timepoint interaction (*, *p*<0.01). C1 barrel, LME model, ANOVA for fixed effects of whisker (C1 vs. other whiskers at day 0, *p*<0.001), timepoint (day 0 vs day 46 for C1, ###, *p*<0.001), and whisker*timepoint interaction (*, *p*<0.05), with significance for individual coefficients for whisker*timepoint interactions, corrected using Benjamini and Hochberg’s method, indicated over corresponding data points (**, *p*<0.01).

**Figure 4.**
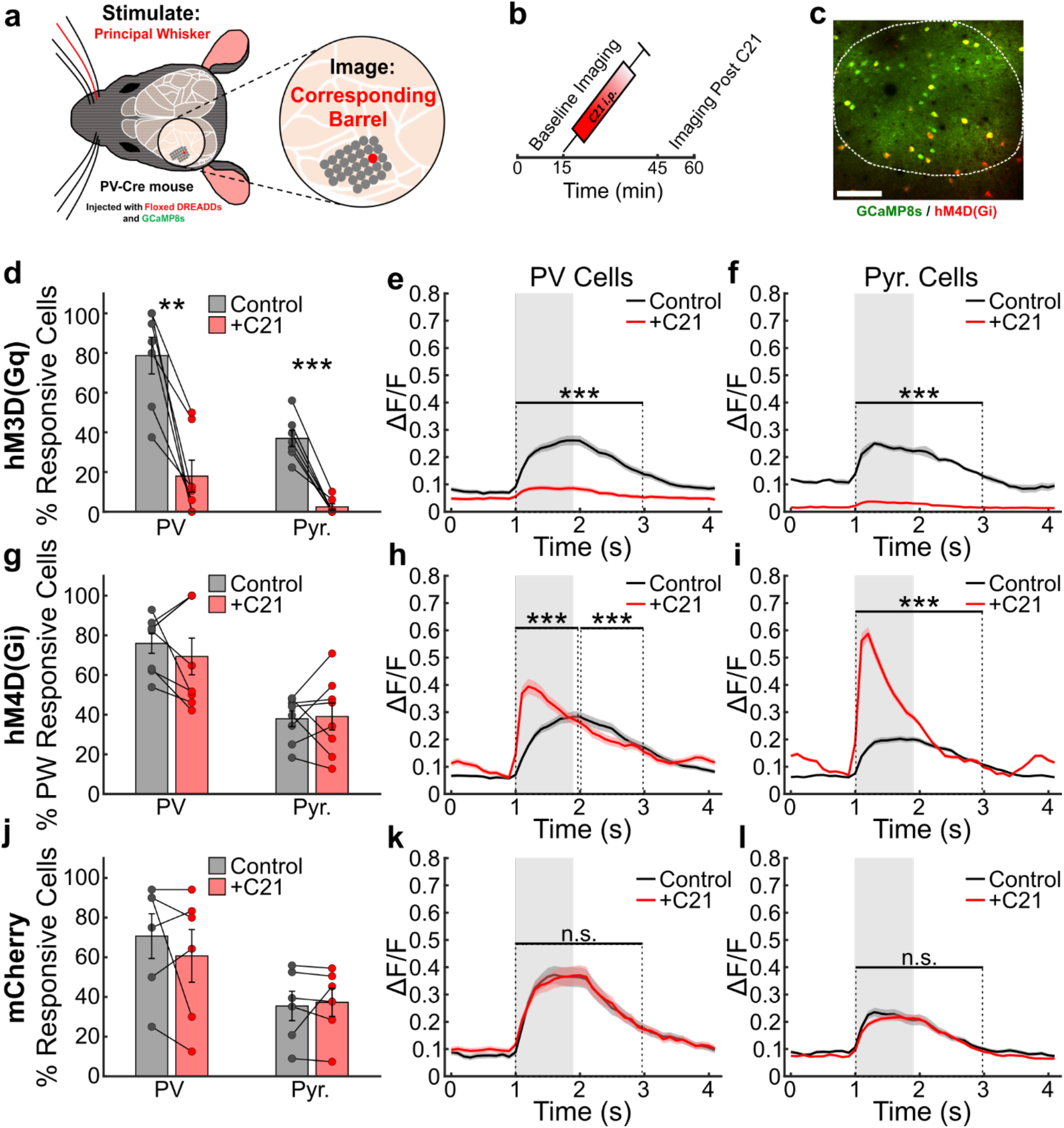
Acute chemogenetic modulation of PV cells affects whisker-evoked activity of local PV and Pyr cells. **a.** Schematic depicting mouse line, AAVs injected, imaging FOV over one barrel and stimulation of the barrel’s PW. **b.** Timeline of the experiment. **c.** 2P representative FOV showing PV cells in yellow expressing both mCherry-tagged hM4D(Gi) DREADDs and jGCaMP8s, PV cells in red expressing only hM4D(Gi), and Pyr cells in green expressing only jGCaMP8s. Dotted outline corresponds to the functional IOSI map of the imaged barrel. Scale bar=100 µm. **d.** Percentage of PV and Pyr cells responsive to PW stimulation in mice (n=7) expressing hM3D(Gq) in PV cells before and after C21 (5 mg/kg, i.p.). Paired t-test comparing mice before and after C21 (PV, **, *p*<0.01; Pyr, ***, *p*<0.001). **e.** Mean evoked ΔF/F calcium trace for all PV cells before and after C21 (n=92 cells). LME model for the MI (Time=1-3 s), fixed effect of condition (***, *p*<0.001). **f.** Mean evoked ΔF/F calcium trace for all Pyr cells before (n=813 cells) and after (n=667 cells) C21. LME model for the MI (Time=1-3 s), fixed effect of condition (***, *p*<0.001). **g.** Percentage of PV and Pyr cells responsive to PW stimulation in mice (n=8) expressing hM4D(Gi) in PV cells before and after C21. Paired t-test comparing mice before and after C21 (PV, *p*=0.41; Pyr, *p*=0.85). **h.** Mean evoked ΔF/F calcium trace for all PV cells before and after C21 (n=132 cells). LME models for the MI (early: Time=1-2 s; and late: Time=2-3 s), fixed effect of condition (***, *p*<0.001). **i.** Mean evoked ΔF/F calcium trace for all Pyr cells before (n=1113 cells) and after (n=979 cells) C21. LME model for the MI (Time=1-3 s), fixed effect of condition (***, *p*=0.001). **j.** Percentage of PV and Pyr cells responsive to PW stimulation in mice (n=6) expressing mCherry in PV cells before and after C21. Paired t-test comparing mice before and after C21, (PV, *p*=0.39; Pyr, *p*=0.71). **k.** Mean evoked ΔF/F calcium trace for all PV cells before and after C21 (n=71 cells). LME model for the MI (Time=1-3 s), fixed effect of condition (*p*=0.21). **l.** Mean evoked ΔF/F calcium trace for all Pyr cells before (n=598 cells) and after (n=617 cells) C21. LME model for the MI (Time=1-3 s), fixed effect of condition (*p*=0.75).

**Figure 5.**
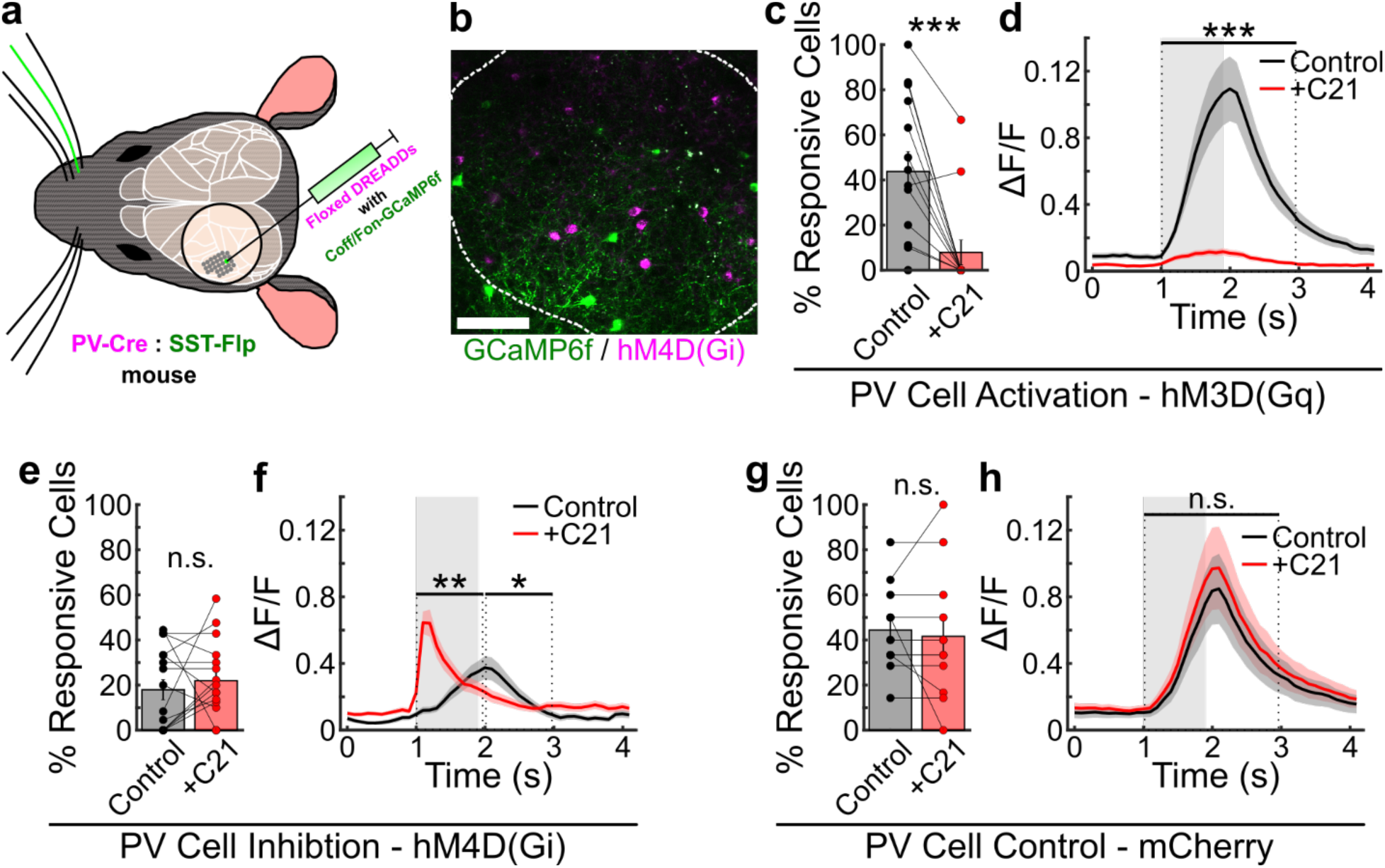
Acute chemogenetic modulation of PV cells affects whisker-evoked activity of local SST cells. **a.** Schematic depicting mouse line, AAVs injected, imaging FOV over one barrel and stimulation of the barrel’s PW. **b.** 2P representative FOV showing SST cells expressing the CreOff/FlpOn-GCamP6f (green) and PV cells expressing hM4D(Gi) DREADDs (magenta). Dotted outline corresponds to the functional IOSI map of the imaged barrel. Scale bar=100 µm. **c.** Percentage of SST cells responsive to PW stimulation in mice (n=14) expressing hM3D(Gq) in PV cells before and after C21 (5 mg/kg, i.p.). Wilcoxon signed rank test comparing mice before and after C21 (***, *p*<0.001). **d.** Mean evoked ΔF/F calcium trace for all SST cells before and after C21 (n=137 cells). LME model for the MI (Time=1-3 s), fixed effect of condition (***, *p*<0.001). **e.** Percentage of SST cells responsive to PW stimulation in mice (n=16) expressing hM4D(Gi) in PV cells before and after C21. Wilcoxon signed rank test comparing mice before and after C21, (*p*=0.33). **f.** Mean evoked ΔF/F calcium trace for all SST cells before and after C21 (n=175 cells). LME model for the MI (early: Time=1-2 s; and late: Time=2-3 s), fixed effect of condition (*, *p*<0.05; **, *p*<0.01). **g.** Percentage of SST cells responsive to PW stimulation in mice (n=12) expressing mCherry in PV cells before and after C21. Paired t-test comparing mice before and after C21 (*p*=0.61). **h.** Mean evoked ΔF/F calcium trace for all SST cells before and after C21 (n=80 cells). LME model for the MI (Time=1-3 s), fixed effect of condition (*p*=0.70).

The selectivity index (SI) of individual cells was calculated using the MI to the C1 and D1 whiskers as follows:

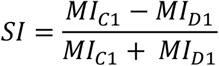

Negative MI values were set to 0 to constrain SI values between -1 and 1, with SI=-1 indicating perfect selectivity for the D1 whisker and SI=1 indicating perfect selectivity for the C1 whisker. For spontaneous activity, amplitude and frequency of calcium transients were detected using the “findpeaks” function in MATLAB with settings of “MinPeakHeight” = 4, “MinPeakDistance” =

0.5 s, and “MinPeakProminence” = 3. We calculated the mean Z-score and area-under-the-curve (AUC) by quantifying the mean and trapezoidal integral of the entire modified Z-scored calcium trace, respectively. For experiments expressing DREADDs in PV cells (**Figs. 4 and 5**), we used the ΔF/F traces to analyze spontaneous activity and set “MinPeakHeight” and “MinPeakProminence” to 4 and 3 times the standard deviation of the baseline fluorescence, respectively. To determine the percentage of PV cells co-modulated by whisker stimuli, for each of the 20 whisker stimuli trials we calculated the MI of each cell, determined the percentage of PV cells within the imaging FOV with MI≧1 on that trial, and then obtained the mean percentage of co-modulated cells across all trials per mouse. To determine cross-correlations between PV cells, we computed the auto-correlation across the entire Z-scored calcium fluorescence trace (either during whisker stimuli trials or for spontaneous activity without whisker stimuli) between individual PV cells and all other PV cells within the imaging FOV using the “xcorr” function in MATLAB. As a control, we also computed the auto-correlation between scrambled traces of PV cells in the FOV in which epochs containing calcium fluorescence transients were shuffled in time.

Principal components analysis to predict changes in responsivity of PV cells over time was performed using the “pca” function in MATLAB. A matrix was created with individual cells as rows and column vectors including the barrel in which cells were located (C1 or D1), MI to C1 and D1 whisker deflection, SI, and the mean Z-score of the spontaneous activity trace for each cell. The same data was used to fit a generalized linear mixed effects (GLME) model with a binomial distribution to predict changes in responsivity of PV cells over time. Receiver operating characteristic analysis of the fitted model was then performed using the “perfcurve” function in MATLAB, with 1000 bootstrap replicas. Tuning curves for PV cells in the C1 barrel were calculated by sorting MI to the C1, D1, B2, and E3 whiskers in descending order, with negative MI set to zero. Responses to each whisker were then normalized to the preferred whisker (the whisker with the largest MI) and the data was fit with a single-term exponential decay function. Decay constants were then used as a measure of tuning width, with larger decay constants indicating more narrow tuning.

### DREADD Agonist Administration

For acute chemogenetic manipulations, mice were imaged under light anesthesia as described above to obtain pre-DREADD ligand C21^69^ activity measures. The imaged barrel varied across mice depending on the specific location in which we found DREADDs and GCaMP were expressed after stereotaxic injections (**Figs. 4 and 5**). We always stimulated the PW of the barrel being imaged for these experiments. Following the pre- C21 imaging, C21 (5 mg/kg, i.p.) was administered and isoflurane anesthesia was turned off. Mice remained on the microscope stage on a heated blanket for 25 minutes. Anesthesia was then resumed as in the pre-C21 imaging session, with ∼1 minute 5% isoflurane induction followed by maintenance at ∼0.7% isoflurane for 4 minutes. Imaging was then repeated exactly as performed in the pre-C21 session. For experiments testing the effects of DREADDs in PV cells on PV and Pyr cell activity, we tested C21 doses of 1 mg/kg, 3 mg/kg and 5 mg/kg in separate imaging sessions (**Fig. S10**). For experiments testing the effects of DREADDs in PV cells on SST cell activity, we tested on a dose of 5 mg/kg.

### Immunohistochemistry

Mice were anesthetized with 5% isoflurane until respirations ceased and transcardially perfused with ice cold phosphate buffer saline PBS (0.1 M), followed by ice cold 4% paraformaldehyde in PBS, and then brains were post-fixed overnight at 4°C. Coronal sections (50 µm) were cut on a vibratome (Precisionary Instruments Inc, #VF-510-0Z). Sections were permeabilized with 0.3% Triton X-100 and blocked with 5% normal goat serum (NGS) for 1 hour at room temperature. Sections were then incubated overnight at 4°C with the primary antibody diluted in PBS with 5% NGS and 0.1% Triton X-100. The following primary antibodies were used: mouse monoclonal anti-parvalbumin (1:250, P3038, Sigma Aldrich) and rabbit polyclonal anti-somatostatin (1:250, ab108456, Abcam). After rinsing 3 times in PBS for 10 min, sections were incubated for 2 hours at room temperature with the corresponding Alexa Fluor-conjugated secondary antibody diluted in PBS (Alexa Fluor 405 for anti-parvalbumin and Alexa Fluor 647 for anti-somatostatin, 1:250, Invitrogen). After rinsing 3 times in PBS for 10 min, sections were mounted onto charged glass slides (Genesee Scientific, #30-185) with mounting medium (Invitrogen, 00-4958-02) and stored in the dark at room temperature. Images were acquired using a Nikon microscope (ECLIPSE Ti2. Objective: 10X and 20X) and manually analyzed with ImageJ (https://imagej.nih.gov/ij/). We counted the total number of cells positive for PV staining, the total number of cells expressing mCherry-tagged DREADDs in PV cells, the colocalization between these markers.

### Mean-field circuit model

We modeled four interacting populations: excitatory pyramidal neurons (Pyr, indexed by E in the equations) and PV, SST and VIP inhibitory neurons. Their non-negative mean firing rates *r_i_*(*t*) followed

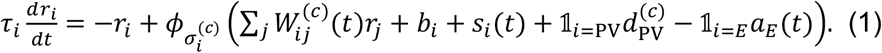

Here *c* denotes Control, hM4D(Gi) or hM3D(Gq). 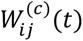 is the effective connection from population *j* to *i*. *b_i_* and *s_i_*(*t*) are background and sensory inputs, 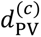 is tonic PV input, and *a_E_*(*t*) is Pyr adaptation. Base weights obeyed Dale’s law^70^. Connectivity, background inputs, time constants, adaptation and short-term plasticity parameters were shared across conditions. Sensory inputs had a common temporal profile, constant during the 1-s stimulus and zero otherwise. Their amplitudes were population-specific and fixed across conditions; VIP received no direct sensory input. Condition dependence entered through tonic PV input, PV output efficacy and transfer-function softness.

The transfer function was a Gaussian-smoothed rectifier^71^,

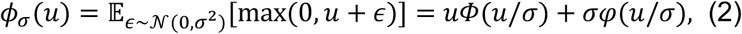

where *Φ* and *φ* are the standard normal cumulative distribution and density. Larger softness *σ* broadens the transition around threshold and raises subthreshold responses. It represents effective input variability unresolved by the population rate^71,72^.

Pyr adaptation^73^ and short-term depression at Pyr→PV and facilitation at Pyr→SST synapses^74,75^ followed

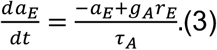

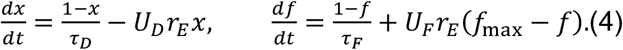

The factors *x*(*t*) and *f*(*t*) multiplied the corresponding connections in *W*^(*c*)^(*t*).

*Representation of hM4D(Gi) and hM3D(Gq) manipulations:* hM3D(Gq) increased tonic PV input^69^ 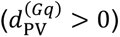, whereas hM4D(Gi) decreased it^76^ 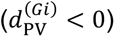 and reduced PV output efficacy, motivated by presynaptic suppression of transmitter release^77,78^. A single factor 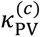 scaled every PV output, including connections deemed negligible (Supplementary Information, Model parameters). It equaled one in Control and hM3D(Gq) and was fitted below one in hM4D(Gi). The shunting label in Fig. 6a refers to this reduction in PV output efficacy rather than to a specific biophysical mechanism. Control tonic input was zero. Softness was parameterized as 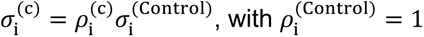. We imposed 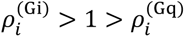 based on the ordering of baseline fluorescence noise in Pyr and PV neurons (**Fig. S9**). Extending this ordering to SST and to unmeasured VIP was a model assumption. These directions were prescribed, and the magnitudes were fitted.

**Figure 6.**
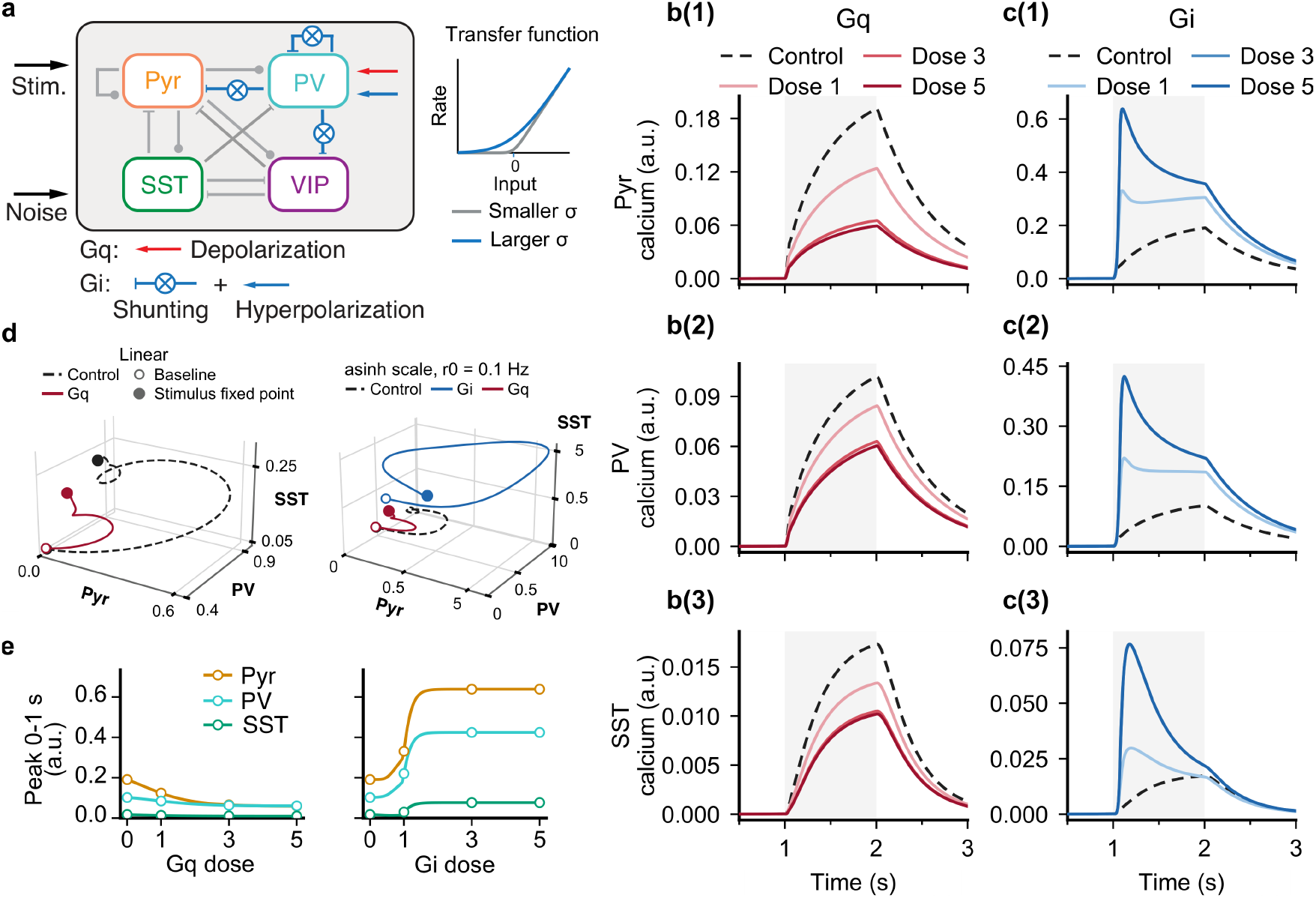
A cell-type-specific circuit model captures the opposing effects of hM3D(Gq) and hM4D(Gi) manipulation on sensory-evoked responses. **a.** Schematic of the four-population model, comprising excitatory pyramidal neurons (Pyr) and inhibitory PV, SST and VIP neurons. hM4D(Gi) decreases tonic PV input and outgoing synaptic efficacy (shunting), whereas hM3D(Gq) increases tonic PV input without changing PV output efficacy. The noise arrow denotes effective input variability, represented in the model by transfer-function softness. The transfer-function inset shows how greater input variability broadens the transition around threshold and increases subthreshold responses. Both manipulations also change transfer-function softness (Methods). Only connections with fitted magnitude ≥0.15 are drawn; all connections were retained in simulations. **b, c.** Baseline-subtracted model-calcium traces for hM3D(Gq) (**b**) and hM4D(Gi) (**c**) in the Pyr (1), PV (2) and SST (3) populations. Dashed black lines denote Control, colored lines denote doses 1, 3 and 5, and gray shading marks the stimulus at 1–2 s (onset at 1 s). Dose-3 and dose-5 traces overlap in c and nearly overlap in b because both fitted dose paths are close to saturation by 3 mg/kg (Fig. S15a,b). **d.** Population-rate trajectories from the prestimulus fixed point to the stimulus-period fixed point, projected into Pyr–PV–SST rate space. The left plot shows Control and hM3D(Gq) dose 5 on linear axes; the right plot shows Control, hM3D(Gq) dose 5 and hM4D(Gi) dose 5 after an asinh(*r*/0.1 Hz) display transformation that compresses the larger hM4D(Gi) range. Tick labels report untransformed rates in Hz. Open and filled circles denote the prestimulus and stimulus-period fixed points, respectively; the Control and hM3D(Gq) prestimulus markers nearly overlap in the linear view. **e.** Peak baseline-subtracted model-calcium response over Time=1-2 s (0–1 s after stimulus onset) across the hM3D(Gq) (left) and hM4D(Gi) (right) dose paths. Dose 0 denotes Control, symbols mark C21 doses of 0, 1, 3 and 5 mg/kg, and lines show the corresponding continuous dose-response curves. Intermediate-dose SST values in **b**, **c** and **e** are model predictions because matched experimental measurements were unavailable. Model-calcium responses are reported in arbitrary units; because they include the fixed observation gain, they are on the scale of the fitted ΔF/F targets (labeled ΔF/F in Figs. S15 and S16).

### Calcium observation and response measures

Absolute rates were filtered as

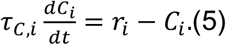

The fixed time constants were 0.60 s for Pyr, PV and VIP and 0.35 s for SST, approximating calcium-fluorescence filtering^59,79^. After subtracting each condition’s prestimulus equilibrium, we applied a fixed population gain: 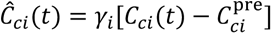. The observation parameters were shared across conditions. Targets were population-mean *ΔF*/*F* traces from all recorded cells, without responsiveness selection; each trial’s prestimulus mean was subtracted before averaging. Peak and AUC denote the maximum and integral over Time=1-2 s (0–1 s after stimulus onset). Traces were simulated for 3 s from stimulus onset, starting from each condition’s prestimulus equilibrium, and sampled at the experimental acquisition times. Shape agreement used Pearson correlation over 0–3 s after stimulus onset.

### Parameter estimation and model selection

We fitted nine traces spanning Pyr, PV and SST under Control and dose-5 hM4D(Gi) and hM3D(Gq), using the hM4D(Gi)-group pre-C21 recordings as the shared Control target. The optimization included 49 circuit parameters; calcium time constants and population observation gains were fixed across conditions and excluded from optimization. Fitted values are listed in Supplementary Tables S1–S4. We optimized with Adam^80^ in double precision, using three stages with increasing emphasis on early responses (Supplementary Information, Parameter search). For window *T*,

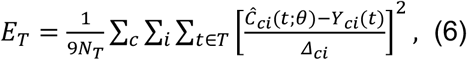

where *Y_ci_* is the measured trace, *θ* the circuit parameters, *N_T_* the number of samples and *Δ_ci_* = max[range_[0,3]_*Y_ci_*, 0.05]. All windows, measured from stimulus onset, used this full-trace normalization. Supplementary Information (Parameter search) summarizes the additional fitting terms, search stages and independent evaluation.

Candidates were re-simulated at 1 ms from independently reconstructed equilibria. Rather than simply taking the candidate with the lowest training loss, we selected the reported model from the 80 final-stage candidates by qualitatively assessing hM4D(Gi) timing and shape, monotonic suppression along the combined hM3D(Gq) path, and compatible prestimulus states within stable, bounded dynamics.

### Dose-dependent model analysis

We calibrated each condition’s dose path using Pyr and PV AUCs at 0, 1, 3 and 5 mg/kg C21. SST supplied only Control and dose-5 measurements, so its intermediate-dose responses are predictions. Shared circuit and observation parameters remained fixed.

A common progress variable ℎ*_c_*(*d*) moved all condition-dependent parameters from Control to the fitted dose-5 endpoint. With expit(*x*) = (1 + *e*^−*x*^)^−1^,

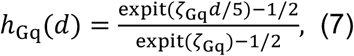

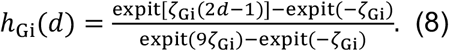

Both satisfy ℎ*_c_*(0) = 0 and ℎ*_c_*(5) = 1. The hM4D(Gi) geometry concentrates progression at lower doses, whereas hM3D(Gq) progresses more gradually. For each condition, only the positive span *ζ_c_* was fitted, using endpoint-normalized model and empirical AUC curves (Supplementary Information, Dose calibration). Parameters varied as

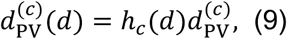

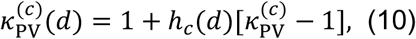

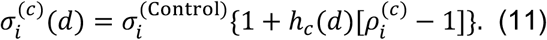

Parameters without a dose argument denote dose-5 values, and hM3D(Gq) PV output efficacy remains one throughout.

### Statistical Analyses

All data are plotted as mean +/- standard error of the mean, unless otherwise stated. Sample sizes were not based on *a priori* power calculations but are consistent with other studies in the field using similar techniques, including our own^67,81^. Statistical analyses were performed in MATLAB. Imaging files were blinded to condition during cell selection. Data were tested for normality using the Lilliefors test and analyzed using parametric or non-parametric statistical tests, as indicated in the figure legends. To account for nested data (measuring multiple cells nested within individual mice) we used mixed effects models^82^. For experiments measuring percentage of whisker-responsive cells across timepoints or conditions (**Figs. 1d, 2c, S1e, S2a, S3b, S4a),** we used GLME models with a binomial distribution, with individual mouse ID modeled as a random effect. For analysis of MI, SI, AUC, peak amplitude, and peak frequency, we used linear mixed effects (LME) models. When individual cells were tracked longitudinally, cells were assigned a unique index and these were nested within individual mice as a random effect. For models with more than one fixed effect, ANOVA was used to test for overall significance of fixed effects. For fixed effects that achieved significance at the level of the ANOVA, *p* values for individual coefficients from the model were corrected for multiple hypothesis testing using the Benjamini & Hochberg procedure^83^. In the figures, results are presented with *p* values for overall fixed effects shown as an inset on respective graphs, with *p* values for individual coefficients depicted adjacent to the respective data points where appropriate. Full statistical results and model specification can be found in the supplementary data.

### Data Availability

Analysis code and links to data tables are available at https://github.com/zeigerlab/Campos-PV-Cells-S1BF-2026.git.

## Results

### Responses of PV cells to principal and surround-whisker stimulation

To understand how the activity of PV cells changes with experience, we first sought to define the spatial distribution of whisker-evoked responses in PV cells. To record neuronal activity from PV cells, we crossed PV-Cre mice with mice expressing the Cre-dependent genetically encoded calcium indicator GCaMP6s^63^. We placed chronic cranial windows over the S1BF and used intrinsic optical signal imaging (IOSI) to localize the cortical C1 and D1 barrel representations (**Fig. S1a**). We then performed 2P in vivo calcium imaging from L2/3 PV cells in fields-of-view (FOV) centered on the C1 barrel while individually deflecting the contralateral principal whisker (PW) C1, and the surround whiskers (SWs) D1, B2, and E3, at 10 Hz for 1 second, delivering 20 trials total for each whisker (**Fig 1a-c**). To identify whisker responsive cells, we used a probabilistic bootstrapping method to correlate calcium transients with epochs of stimulation. We also included a threshold for activity, such that a cell had to have a stimulus evoked modulation index (MI) equal or greater than 1 Z-score to be considered responsive. This threshold removed a small number of PV cells with spurious correlations to stimulation epochs (**Fig. S1b-d**). We tested different waveforms for whisker deflection (square wave or sine wave) and found that faster rise times for square wave deflections and larger amplitude deflections for sine wave deflections led to higher numbers of whisker-responsive PV cells (**Fig. S1e**). We chose a square wave deflection with 1 ms rise time for all further experiments as this led to robust activation of PV cells with clear whisker-evoked calcium transients (**Fig. 1c**). There were no significant differences in the number of whisker-responsive cells or the magnitude of whisker-evoked responses between awake and lightly anesthetized animals using this protocol (**Fig. S1f-g**). For that reason, and to reduce variability associated with behavioral state, we performed all subsequent recordings in lightly anesthetized mice.

We next quantified whisker-evoked activity of PV cells in FOV centered on the C1 barrel. We found that a majority of cells (85.7% +/- 2.5%) responded with time-locked responses to deflections of the PW C1 (**Fig. 1d**). Fewer cells responded to SWs, with the number of whisker responsive neurons decreasing as the distance of the SWs increased from the PW C1 (46.1% +/- 4.1%, 34.1% +/- 14.0%, and 5.6% +/- 5.6% for the SWs D1, B2, and E3, respectively) (**Fig. 1d**). Mean whisker-evoked responses averaged across all trials for all individual PV cells showed that evoked responses were qualitatively substantially higher for PW C1 deflections compared to the other SWs (**Fig. 1e**). Given that significantly more cells were responsive to the PW C1 than the other SWs, this could account for the substantially higher whisker-evoked responses when averaged across all cells. Therefore, we classified PV cells into groups of either “responders” or “non-responders” based on their evoked activity to each whisker to compare whisker-evoked responses across these groups. As expected, clear whisker-evoked activity was seen for cells classified as responders (**Fig. 1f**), whereas mean evoked traces were relatively flat for cells classified as non-responders (**Fig. 1g**). We then calculated the MI to each whisker to quantify the strength of whisker-evoked responses in each PV cell. In responders, whisker-evoked activity was greatest for the PW C1 (MI=8.3 +/- 0.6) and significantly lower for the SWs D1 (MI=3.9 +/- 0.6) and B2 (MI=3.1 +/- 0.3). Whisker-evoked activity for the SW E3 (MI=4.7 +/- 1.2) was also lower, but this result was not significant due to the low number of E3 responsive PV cells in the C1 barrel (**Fig. 1h**). In non-responders, whisker-evoked activity was low across all whiskers, as expected (**Fig. 1i**). Finally, we plotted the position of all responsive cells (relative to the FOV centered on the C1 barrel) imaged across all mice to see if there is spatial clustering of PV cells tuned to specific whiskers. PW C1 responsive PV cells were distributed relatively homogenously across the C1 barrel FOV, whereas the SW D1 responsive PV cells in the C1 barrel FOV tended to cluster in the anteromedial portion of the FOV, closest to the D1 barrel (**Fig. 1j**). SW B2 and E3 responsive PV cells were much rarer and were distributed throughout the C1 barrel FOV (**Fig. 1j**). We repeated the same experiment, imaging this time in the D1 barrel and stimulating the PW D1 and the SW C1, and found similar results. More cells were responsive to the PW D1 in the D1 barrel, whisker-evoked responses in responders were higher for the PW D1 compared to the SW C1, and SW C1 responsive PV cells were clustered in the posterolateral portion of the D1 barrel FOV, closest to the C1 barrel (**Fig. S2**).

### Whisker trimming induces a dynamic recruitment of PV cells responsive to the spared whisker in the deprived barrel

Having defined the spatial distribution of PV cell responses to different whiskers in the adult S1BF, we next sought to determine how PV cell activity changes during experience. We implemented chronic whisker trimming, a well-studied paradigm for inducing experience-dependent plasticity. Macroscopically, chronic whisker trimming leads to expansion of cortical map representations of spared whiskers and contraction of maps for deprived whiskers^15–18^. One day after whisker trimming, intrinsic excitability of PV cells in deprived barrels is reduced^84^. This results in disinhibition of Pyr cells and increased whisker-evoked responses to trimmed whiskers acutely, but over time as trimming is maintained Pyr cell responses to the trimmed whisker become depressed^57^. However, how PV responses change in spared barrels and during more chronic phases of trimming remains unknown. To investigate this, we trimmed all contralateral whiskers except D1 and longitudinally imaged individual PV cells in the C1 (deprived barrel) and D1 (spared barrel) barrels over 18 days of whisker trimming followed by 28 days of regrowth (**Figs. 2a, S3a**). Prior to whisker trimming, the selectivity of individual PV cells in the C1 and D1 barrels were clearly separable, with PV cells in each barrel more selective for the PW of the corresponding barrel (SI in D1=-0.37 +/- 0.03; SI in C1=0.46 +/- 0.03; **Fig. 2b**).

In the spared D1 barrel, the number of PW D1 responsive neurons was high and did not change significantly over 18 days of trimming or after whisker regrowth (Baseline=72.8% +/- 2.8%; Day 18=71.8% +/- 5.2%; Day 46=75.3% +/- 3.9%; **Fig. 2c**). Mean whisker-evoked responses of PW D1 responsive PV cells in the D1 barrel increased slightly after 18 days of whisker trimming and returned to baseline levels after regrowth (MI: Baseline=6.9 +/- 0.6; Day 18=8.2 +/- 0.7; Day 46=7.8 +/- 1.0; **Fig. 2d**). In the deprived C1 barrel, the number of SW D1 responsive cells significantly increased after 12 and 18 days of whisker trimming and was reduced compared to baseline after whisker regrowth (Baseline=46.1% +/- 4.1%; Day 12=59.4% +/- 6.7%; Day 18=59.2% +/- 4.5%; Day 46=37.2% +/- 5.2%; **Fig. 2c**). Mean whisker-evoked responses of SW D1 responsive PV cells in the deprived C1 barrel column did not change significantly over time (MI: Baseline=3.9 +/- 0.6; Day 18=3.5 +/- 0.3; Day 46=3.2 +/- 0.4; **Fig. 2d**). The same overall trends were observed in recruitment of D1 responsive PV cells and MI of D1 responsive PV cells in the spared D1 barrel and the deprived C1 barrel if we removed the MI threshold for classifying cells as responsive (**Fig. S3b-c**). The mean MI across all PV cells (both D1 responsive and D1 non-responsive) increased to PW D1 whisker stimulation in the spared D1 barrel during trimming (**Fig. S3d**), whereas the mean MI across all PV cells to SW D1 whisker stimulation in the deprived C1 barrel showed no change during trimming but decreased after regrowth (**Fig. S3e**).

To further analyze how population activity to whisker stimulation of PV cells changed with trimming, we quantified the mean proportion of positively modulated PV cells during individual whisker stimuli trials (MI≧1) and the mean cross correlation between PV cells in each imaging FOV. The mean proportion of co-modulated cells to D1 whisker stimulation did not change in either the spared D1 barrel or the deprived C1 barrel during whisker trimming but was reduced in the deprived C1 barrel after whisker regrowth (**Fig. S3f**). The cross correlation of activity of PV cells in the spared D1 barrel increased slightly throughout whisker trimming and decreased after whisker regrowth (**Fig. S3g**). In contrast, the cross correlation of activity of PV cells in the deprived C1 barrel was reduced from Day 3 and remained lower than baseline after regrowth (**Fig. S3h**).

In control mice that we imaged without whisker trimming, we did not see any recruitment of D1 responsive PV cells in either the D1 or C1 barrels and there was no change in mean D1 whisker-evoked responses of D1 responsive cells (**Fig. S4b**-**c**). Whisker trimming did not result in a significant change in the length of trimmed whiskers after regrowth (**Fig. S4c**). We also quantified changes in the spontaneous activity of PV cells over time during whisker trimming. In the spared D1 barrel, total spontaneous activity (measured as the area-under-the-curve [AUC] across ∼150 s of activity) of all PV cells changed minimally during trimming but was reduced after whisker regrowth (**Fig. S5a**). More significant changes were seen in the deprived C1 barrel, where spontaneous activity was reduced during whisker trimming and remained below baseline levels after whisker regrowth (**Fig. S5a**). These changes were driven primarily by reductions in the amplitude of calcium transients, as calcium transient peak frequency did not change over time in the deprived C1 barrel (**Fig. S5b-c**). The cross correlation of spontaneous activity of PV cells in the spared D1 barrel showed inconsistent changes during whisker trimming, with small reductions initially after 3 and 6 days of trimming, but a more substantial decrease after whisker regrowth (**Fig. S5d**). Similar trends were observed in the deprived C1 barrel, with reductions in the cross correlation of spontaneous activity of PV cells after 3, 6, and 12 days of trimming as well as after whisker regrowth (**Fig. S5e**).

In addition to population level changes, we also quantified changes in responsivity of individual PV cells longitudinally over time. In the spared D1 barrel, 7.9% of cells that were non-responsive to the PW D1 at baseline became responsive after 12 days of whisker trimming (**Fig. 2e**). This was balanced by 5.8% of cells switching from responsive at baseline to non-responsive after 12 days, resulting in a similar number of responsive PV cells overall (**Fig. 2e**). In the deprived C1 barrel, 6.2% of cells switched from responsive to SW D1 at baseline to non-responsive after 12 days, but 22.4% of cells switched from non-responsive to responsive (**Fig. 2f**), consistent with a recruitment of initially non-responsive cells to the SW D1 after whisker trimming. In both the spared D1 barrel and the deprived C1 barrel, the change in MI to the PW D1 after 12 days of whisker trimming was positive (cells became more responsive) and higher for cells switching from non-responsive to responsive (NR-R) compared to cells that remained non-responsive (NR-NR; **Fig. 2g-h**). In contrast, the change in MI for cells switching from responsive to non-responsive (R-NR) was negative (cells became less responsive) and lower compared to cells that remained responsive (R-R; **Fig. 2g-h**). Similarly, in both the spared D1 barrel and the deprived C1 barrel, the mean MI to D1 whisker stimulation 12 days after whisker trimming was highest for persistently responsive R-R cells and was increased in NR-R cells compared to NR-NR cells (**Fig. 2i-j**). Interestingly, the mean MI to D1 whisker stimulation 12 days after whisker trimming in NR-R cells was lower compared to R-R cells, despite both populations of cells being classified as D1 responsive at this timepoint (**Fig. 2i-j**). This suggests sensory evoked responses are weaker in PV cells newly recruited to become responsive to the spared D1 whisker.

We next sought to determine if baseline neuronal activity could predict which cells might change their responsivity to the spared D1 whisker after whisker trimming. At baseline in the spared D1 barrel, persistently D1 responsive R-R cells had the highest MI to both PW D1 and SW C1 stimulation (**Fig. S6a-b**), significantly higher than both NR-NR and NR-R cells (**Fig. S6a-b**). There was no difference in mean MI to either the PW D1 or SW C1 between NR-NR and NR-R cells in the spared D1 Barrel (**Fig. S6a-b**). Selectivity indices of cells in the spared D1 barrel were biased to the PW D1 across all classes of cells, as expected, with no significant differences among classes (**Fig. S6c**). In the deprived C1 barrel, trends in MI to SW D1 stimulation were similar across classes to the trends seen in the spared D1 barrel, with persistently D1 responsive R-R cells having higher response magnitudes than NR-NR and NR-R cells, and no difference between NR-NR and NR-R cells (**Fig. S6d**). The MI of NR-R cells in the deprived C1 barrel to the PW C1 were higher than NR-NR cells, the only case where baseline responses differed between NR-NR and NR-R cells (**Fig. S6e**). Selectivity of all classes of cells in the deprived C1 barrel were biased to the PW C1, as expected (**Fig. S6f**). Persistently D1 responsive R-R cells in the deprived C1 barrel had lower SI for the PW C1 at baseline compared to D1 whisker non-responders (NR-NR and NR-R cells, **Fig. S6f**), consistent with many of these cells having strong responses to the SW D1 even before trimming.

These data suggested that individual activity metrics were poor predictors of which cells might change responsivity to the spared D1 whisker over time, especially for baseline non-responders to the D1 whisker (NR-NR and NR-R cells). To see if multiple baseline metrics better predict which non-responders switch classes over time, we performed a principal components analysis of non-responder cells (NR-NR and NR-R cells) using the barrel location, MI to C1 and D1 whisker deflection, SI, and the mean Z-score of the spontaneous activity trace for each cell. NR-NR and NR-R cells were poorly separable by PCA (**Fig. S6g**). The first principal component explained 89% of the variance and was heavily weighted toward response magnitude to C1 whisker stimulation at baseline (coefficient = 0.998). We next tried fitting a generalized linear mixed effects (GLME) model to these data to classify cells into NR-NR or NR-R. Performance of this model as measured by receiver-operating characteristic curve was modest (AUC = 0.77, confidence bounds 0.69-0.83) (**Fig. S6h**). In the model, only the coefficient for the MI to D1 whisker stimulation was significant (*p*<0.001). These analyses show that response magnitude of cells to whisker stimulation at baseline has some predictive value, but that a substantial portion of the variability in which cells will switch from spared D1 whisker non-responders to responders with whisker trimming remains unexplained by our studied measures of baseline activity.

### Persistent changes in PV cell selectivity after whisker regrowth

We next examined changes in PW D1 responsivity in the spared D1 barrel, and SW D1 responsivity in the deprived C1 barrel after whisker regrowth. Individual cells exhibited complex trajectories over time, with all possible changes in responsivity represented across baseline, day 12, and day 46 timepoints (**Fig. 3a-b**). In the spared D1 barrel, changes in responsivity were generally balanced, such that the overall proportion of PW D1 responsive and non-responsive cells was maintained at each timepoint (**Figs. 2c, 3a**). In the deprived C1 barrel, the increase in SW D1 responsive cells seen after 12 days of whisker trimming reversed at day 46 after regrowth, with fewer SW D1 responsive cells compared to baseline (**Figs. 2c, 3b**). In addition to the binary outcome of whether a cell was responsive to the D1 whisker or not, we also compared the selectivity of cells for the C1 and D1 whiskers between baseline and day 46. In the spared D1 barrel, although we did not observe any change in the percentage of PW D1 responsive cells over time, selectivity of cells was even more biased to the spared PW D1 after trimming and regrowth compared to baseline (SI=-0.66 +/- 0.03 at day 46 vs -0.37 +/- 0.03 at baseline, **Fig. 3c**). In the deprived C1 barrel, despite significant shifts in the population of SW D1 responsive cells during whisker trimming, cell selectivity changed only slightly between baseline and day 46, shifting subtly toward the SW D1 (SI=0.39 +/- 0.04 at day 46 vs 0.46 +/- 0.03 at baseline, **Fig. 3d**). We also calculated a measure of tuning width for cells in the C1 barrel across all four stimulated whiskers (PW C1, and SWs D1, B2, and E3), with evoked response magnitudes normalized to the response of the most preferred whisker (**Fig. S7a-b**). Tuning width of PV cells in the C1 barrel shifted slightly between baseline and day 46, becoming narrower (more selective), for the cell’s preferred whisker (**Fig. S7c**).

To better understand these persistent changes in selectivity after whisker regrowth, we calculated the change in the percentage of cells responsive to either the C1 or D1 whisker in each barrel between baseline and day 46 for each mouse. In the spared D1 barrel there was minimal change in the percentage of PW D1 responsive neurons (2.5 ± 3.4% increase), but the percentage of SW C1 responsive neurons decreased substantially (-17.9 ± 6.5%; **Fig. 3e, left**). We also compared changes in the magnitude of whisker-evoked responses and found a small but significant increase in evoked response over time to the PW D1 compared to the SW C1 (ΔMI=0.8 ± 0.7 for D1; -1.2 ± 0.2 for C1; **Fig. 3f, left**). In the deprived C1 barrel, the percentage of PW C1 responsive neurons decreased significantly between baseline and day 46 (-12.1 ± 3.1%), but the percentage of neurons responsive to the SWs D1, B2, and E3 also decreased (- 8.9 ± 6.0%, -32.2 ± 13.7%, and -5.6 ± 5.1%, respectively; **Fig. 3e, right**). The magnitude of whisker-evoked responses was slightly reduced for the SWs D1, B2, and E3 (ΔMI=-0.8 ± 0.2, - 1.3 ± 0.2, and -0.4 ± 0.1, respectively) but was most strongly reduced for the PW C1 (ΔMI=-2.5 ± 0.4; **Fig. 3f, right**). These changes suggest that the persistent shift in selectivity toward the PW D1 in the spared D1 barrel is driven by a reduction in SW C1 responsivity, rather than potentiation of PW D1 responsivity. In the deprived C1 barrel, reductions in PW C1 responsivity are partially balanced by reductions in responsivity to other whiskers, leading to relatively preserved cell selectivity for PW C1 after whisker regrowth compared to baseline.

### Chemogenetic modulation of PV cells affects whisker-evoked activity across multiple cell types

To understand how changes in PV cell activity might affect whisker-evoked activity across different components of the S1BF microcircuit, we used chemogenetic DREADDs to manipulate the activity of PV cells (**Fig. 4a-c**). We expressed jGCaMP8s in all neurons using an AAV with a pan-neuronal synapsin promoter and recorded activity of PV cells (expressing mCherry-tagged DREADDs) and presumptive Pyr cells (mCherry negative) before and after injection of the DREADD ligand C21^69^ (5 mg/kg). DREADDs were expressed in ∼81% of local PV cells (**Fig. S8**). We quantified both spontaneous activity and activity evoked by deflection of the imaged barrel’s PW. In animals expressing the activating DREADD hM3D(Gq) in PV cells, C21 led to a strong reduction in the number of PW responsive PV and Pyr cells (**Fig. 4d**). Activation of PV cells with C21 also strongly suppressed the noise of the baseline fluorescence, affecting interpretation of Z-scored fluorescence traces (**Fig. S9a**). Therefore, we quantified mean PW evoked responses using ΔF/F values to calculate MI across all cells and found these were strongly reduced for both Pyr cells and PV cells (**Figs. 4e-f, S9b**). The effect of C21 on number of responsive cells was dose dependent (**Fig. S10a**), whereas the reduction in MI was evident even at low doses (1 and 3 mg/kg, **Fig. S10b-c**). Total spontaneous activity, calcium transient frequency, and calcium transient amplitudes were all reduced in Pyr cells after C21, whereas only spontaneous transient amplitude was reduced in PV cells after C21 (**Fig. S11a-c**). While somewhat counterintuitive, these results are in agreement with other studies demonstrating a paradoxical suppression of PV cell activity upon activation^85,86^.

In mice expressing the inhibitory DREADD hM4D(Gi) in PV cells, C21 did not change the number of PW responsive Pyr or PV cells (**Fig. 4g**). Opposite to what we observed with hM3D(Gq), in mice expressing hM4D(Gi) in PV cells C21 strongly increased the noise of the baseline fluorescence for both Pyr and PV cells (**Fig. S9c**). Mean PW evoked responses in PV cells showed complex changes after C21, with a larger mean MI early after onset of PW stimulation (Time=1-2 s), but then faster decay and smaller mean MI late after onset of PW stimulation (Time=2-3 s; **Figs. 4h, S9d-e**). After C21, activity of Pyr cells also peaked early but MI was strongly increased after PW stimulation (Time=1-3 s; **Figs. 4i, S9d**). There were no consistent changes in the number of PW responsive PV or Pyr cells across doses of C21 (**Fig. S10d**). The shift in peak activity and increased MI was apparent even at low doses of C21 in Pyr cells but became more apparent in PV cells as doses increased (**Fig. S10e-f**). Total spontaneous activity, calcium transient frequency, and calcium transient amplitude all significantly increased after C21 (**Fig. S11d-f**). Controls expressing mCherry in PV cells did not show any changes in the number of whisker responsive cells or PW evoked responses in PV or Pyr cells after C21 (**Figs. 4j-l, S10g-i**). Some measures of spontaneous activity showed very slight increases after C21 (**Fig. S11g-i**), possibly reflecting off-target effects of C21 or cumulative effects of anesthesia.

In conjunction with PV cells, SST interneurons provide direct inhibition to the dendrites of Pyr cells, contributing to regulation of Pyr cell excitability^87–89^. In addition, there are reciprocal connections between PV and SST cells^74,90–92^. Given the substantial changes observed in local PV cells and Pyr cells following PV cell modulation, we wondered how the activity of SST cells might change following modulation of PV cell activity. To test this, we injected S1BF of PV-Cre:Sst-ires-Flp double transgenic mice with AAVs to express Cre-dependent DREADDs in PV cells and Flp-dependent GCaMP6f in SST cells (**Fig. 5a**). GCaMP6f and DREADDs were expressed in unique cell populations (**Fig. 5b**), with GCaMP6f-expressing cells co-staining for SST (**Fig. S12**). We then recorded spontaneous and whisker-evoked activity from SST cells. In mice expressing the activating DREADD hM3D(Gq) in PV cells, the number of PW responsive SST cells and the mean PW evoked responses were strongly reduced (**Fig. 5c-d**), similar to the changes observed in Pyr cells (**Fig. 4d-f**). Spontaneous activity of SST cells was also decreased (**Fig. S13a-c**). In mice expressing the inhibitory DREADD hM4D(Gi) in PV cells, we found no change in the number of whisker responsive SST cells after C21 (**Fig. 5e**). Mean PW evoked traces from SST cells showed an earlier peak after C21, with higher mean MI early after PW stimulation and lower mean MI late after PW stimulation (**Fig. 5f**), similar to the results observed in Pyr cells (**Fig. 4g-i**). Spontaneous activity of SST cells was also increased after C21 (**Fig. S13d-f**). Controls expressing mCherry in PV cells did not show any changes in the number of whisker responsive cells or mean PW evoked responses in SST cells after C21 (**Fig. 5g-h**) and only a slight increase in the frequency of spontaneous calcium transients (**Fig. S13g-i**). Together, these data demonstrate that modulation of PV cell activity acutely alters microcircuit dynamics across cell types in the S1BF.

### A recurrent circuit model captures DREADD-dependent sensory response dynamics

To test whether recurrent cortical dynamics could account for the paradoxical suppression of sensory-evoked responses during hM3D(Gq) activation of PV neurons (**Figs. 4e-f, 5d**) and the enhanced early responses during hM4D(Gi) suppression (**Figs. 4h-i, 5f**), we constructed a four-population mean-field model of Pyr, PV, SST and VIP neurons (**Fig. 6a**). We modeled hM3D(Gq) activation as a positive tonic input to PV neurons^69^, and hM4D(Gi) suppression as reduced tonic PV drive^76^ combined with reduced efficacy of PV output^93^, consistent with hM4D(Gi) suppression of synaptic release^77,78^. We additionally represented effective input variability through the softness of the population transfer function, because input fluctuations make population firing increase more gradually around threshold^71,72^. Because baseline fluorescence noise followed the order hM4D(Gi) > Control > hM3D(Gq) (**Fig. S9**), we constrained softness to be greatest for hM4D(Gi), smallest for hM3D(Gq) and intermediate for Control (**Fig. 6a**). All remaining parameters, including recurrent connectivity and sensory input, were shared across conditions, under the assumption that they were not affected by DREADD perturbations. We converted model firing rates to calcium-like signals and baseline-subtracted them within each condition for comparison with the experimental ΔF/F traces (**Figs. 4, 5**).

The model reproduced hM3D(Gq) suppression despite the excitatory action of hM3D(Gq) on PV neurons. Baseline-subtracted calcium responses decreased in Pyr, PV and SST in both peak amplitude and area under the curve (AUC) over Time=1-2 s (0–1 s after stimulus onset; **Fig. 6b**). Increased tonic PV input and reduced effective input variability jointly shifted the prestimulus circuit state and the trajectory evoked by the same sensory input. Increasing tonic PV drive alone reduced the early calcium response (0–200-ms AUC), and suppression persisted across combinations of tonic input and softness (**Supplementary Table S7**). An alternative account of the weak hM3D(Gq) responses is that strong depolarization drives PV neurons into depolarization block. However, complete PV silencing in the fitted hM3D(Gq) circuit did not reproduce the observed suppression; instead, it released large onset transients in the remaining populations and drove rates beyond the model’s validity range (**Supplementary Table S8**). PV-mediated inhibition remained substantial in the fitted hM3D(Gq) state (**Fig. S14a–c**), indicating that a smaller evoked calcium increment need not imply a loss of PV-mediated inhibition.

The same circuit produced a qualitatively different response under hM4D(Gi). The model generated a large early transient in Pyr, PV and SST that decayed during the stimulus period (**Fig. 6c**), and in rate space the trajectory overshot its stimulus-period fixed point before relaxing toward it (**Fig. 6d**). Reduced PV output efficacy permitted a pronounced recurrent excitatory transient that peaked before total recurrent inhibition, consistent with a transient excitation– inhibition imbalance underlying the onset response and the resulting early calcium peaks (**Fig. S14d–f**). hM4D(Gi) therefore did not uniformly disinhibit the circuit throughout the stimulus but changed the time course over which recurrent inhibition was recruited. Two complementary tests supported a role for PV-output attenuation in the early-peak phenotype. First, restoring PV output efficacy to its Control value in the fitted model eliminated the early calcium peaks in all three populations (**Fig. S14j,k**). Second, refitting the model with PV output efficacy held at its Control value, under the same optimization conditions, recovered no candidate within the prescribed rate bound with early peaks in all three populations, and the best aggregate fit, whose error was comparable to that of the fitted model, peaked at stimulus offset in every hM4D(Gi) trace (**Supplementary Table S9**). Thus, within the tested model family and optimization conditions, the early peaks in all three populations depended on PV-output attenuation.

We next asked whether the same circuit could account for the dose dependence of these effects. We calibrated a path for each condition between Control and the fitted 5 mg/kg C21 state using the experimental Pyr and PV dose–response AUCs, varying condition-dependent parameters together while shared circuit parameters remained fixed (**Fig. S10c,f**). The calibrated paths reproduced the opposing trends observed across C21 doses. hM3D(Gq) responses decreased progressively toward the 5 mg/kg dose state, whereas hM4D(Gi) responses rose steeply and approached saturation by 3 mg/kg (**Figs. 6e, S15a–d**). To see how far this behavior would extend, we examined a hypothetical linear increase in the hM3D(Gq) parameters beyond their fitted endpoint. PV firing during the stimulus period initially decreased along this path because the increasing drive suppressed Pyr activity and thereby withdrew the recurrent excitation PV neurons receive^74,94,95^. Once Pyr activity approached zero, no further recurrent excitation remained to withdraw, and PV firing rose again with the added drive, placing the fitted hM3D(Gq) state on the descending branch of a nonmonotonic curve (**Fig. S15e**). This demonstrates that, in a manner similar to the paradoxical effect^96^, tonic input drive to PV could in fact increase its activity, but we predict it would need to be increased beyond what we achieved with the doses used here, and perhaps beyond what would be achievable at any dose of C21.

Finally, to test whether these response patterns persisted beyond the mean-field approximation, we expanded the model into a heterogeneous 1,400-unit recurrent network with sparse connectivity, cell-to-cell variability and dynamic input noise. The network’s population mean responses preserved the patterns of the four-population model across Control, hM4D(Gi) and hM3D(Gq) while individual units varied in response amplitude and time course (**Fig. S16a(1),b(1)**). Without fitting individual neuronal traces, the network reproduced the broad temporal structure and population differences of the experimental responses, including hM3D(Gq) suppression and hM4D(Gi) early peaks (**Fig. S16a,b**). Population responses of the fitted four-population model followed the measured trends in amplitude and time course across conditions and doses, although the simulated hM3D(Gq) suppression was smaller than measured (**Fig. S16c–e**). Together, these simulations show that a single recurrent circuit can account for the direction, time course and much of the magnitude of the sensory-evoked responses observed under both hM3D(Gq) and hM4D(Gi). More broadly, the model indicates that a DREADD’s effect on an evoked response depends on the circuit’s baseline state and transient dynamics, so its direct action on the targeted cell type alone does not predict the response.

## Discussion

PV cells are the most common type of inhibitory interneuron in the S1BF. Until now, the specific response properties of PV cells to whisker stimulation and the spatial distribution of those responses have not been well studied. Our data define for the first time the spatial distribution of whisker-evoked PV cell responses within and across barrels. In a given barrel, deflection of the PW evokes responses in the greatest number of PV cells, with fewer PV cells responding as distance of the whisker increases from the PW of the barrel (**Figs. 1d, S2a**). This pattern is similar to the well characterized “salt-and-pepper”-like distribution of Pyr cell responses in L2/3 of S1BF, where PW selective cells are intermixed with other cells that respond best to surround barrel whiskers^12,13^. The number of PV cells responsive to whisker stimulation were higher than we, and others, have reported for Pyr cells, both for the PW (70-86% for PV cells here [**Figs. 1d, S2a**] vs. 25-52% for Pyr cells) and SWs (up to 37-46% for PV cells 1 row/arc distant here [**Figs. 1d, S2a**] vs. 2-20% for Pyr cells)^12,13,81^. We also found that many PV cells are responsive to multiple whiskers (**Fig. S7**). Although no studies have reported responses of PV cells to multiple whiskers in mice, our results are in agreement with earlier recordings from suspected interneurons in L2/3 of rabbit S1BF, where putative interneurons responded on average to ∼5.5 whiskers^38^. Although PV cells often responded to multiple whiskers, one whisker, typically the PW, evoked substantially larger responses compared to other whiskers. For example, the response magnitude of PV cells for the PW was ∼2-fold greater compared to SWs (mean MI 2.2-fold for C1>D1 in the C1 barrel, or 1.7-fold for D1>C1 in the D1 barrel, **Figs. 1h, S2d**). This was reflected in relatively high selectivity indices for most PV cells for the PW vs an adjacent SW (SI=∼0.37-0.46 for PV cells, **Fig. 2b**)^19^. The high selectivity of these cells may be a result of convergent bottom up inputs from L4, which make up the majority of pre-synaptic inputs onto PV cells^97^. In contrast, smaller differences in SI and magnitude of response evoked by deflection of the PW compared to SW have been reported for Pyr cells^12,13,19,81^. These results, suggesting relatively narrow tuning of PV cells for specific whiskers, stand in contrast to the more broad orientation tuning of PV cells compared to Pyr cells in V1^98–100^. On the other hand, while S1BF PV cells may be narrowly tuned to whisker identity, they may be more broadly tuned to other properties of whisker deflection, such as direction^101,102^.

We next determined how PV cell responsivity changes with experience by implementing a chronic whisker trimming paradigm. Whisker trimming is a well-established method for inducing experience-dependent plasticity in S1BF. During whisker trimming, L2/3 Pyr cells shift their selectivity toward spared whiskers, with potentiation of responses to the spared whisker and reductions in response to the deprived whisker^17,19^. These changes are mediated by 1) strongly reduced responses for the trimmed whisker coupled with more modest reductions in responses for the spared whisker in cells with high whisker responsivity at baseline; and 2) small increases in responses to spared whiskers in cells with low whisker responsivity at baseline^17,19^. In deprived barrels, L4 PV-mediated feed-forward inhibition is acutely reduced for the trimmed whisker, mediating disinhibition and transient, homeostatic firing rate stabilization of L2/3 Pyr cells^57,84^. We hypothesized that a similar reduction in L2/3 PV-mediated inhibition might facilitate tuning changes in Pyr cells toward the spared whisker after trimming. However, we found that after trimming all whiskers except D1, the number of SW D1 responsive PV cells increased significantly in the deprived C1 barrel (**Fig. 2c**). This effect was barrel specific, as the number of PW D1 responsive PV cells did not change in the spared D1 barrel (**Fig. 2c**), although there was a small increase in response magnitude in the D1 barrel after 18 days of trimming (**Fig. 2d**). Although there is significant representational drift in the barrel cortex^13,103^, the increase in SW D1 responsive PV cells is unlikely due to random drift as we did not see any increase in control animals without whisker trimming (**Fig. S4a-b**). We tried to determine what factors predict which PV cells will maintain or switch responsivity after trimming. However, we found that these populations largely overlap in terms of baseline activity metrics using PCA (**Fig. S6g**), and a GLME model incorporating baseline neural activity metrics achieved only modest performance (**Fig. S6h**). Baseline response magnitude to the C1 and D1 whiskers contributed most to the PCA and GLME, respectively, but a significant portion of the variability remains unexplained. Prior studies have found that horizontal intracortical connections from excitatory Pyr cells in spared barrels to deprived barrels increase after whisker trimming^104–106^. This could account for the increase in PV cells responsive to the spared whisker we observed in the deprived barrels, though future work will be required to directly test this.

After whisker regrowth, expanded whisker cortical map representations contract back to pre-trimming sizes^15^. Less is known about changes in whisker-evoked neuronal activity after whisker regrowth, but selectivity of Pyr cells for spared and trimmed whiskers also appears to return to pre-trimming levels^19^. For PV cells, we found changes in neuronal activity that persisted even after 28 days of whisker regrowth. In the deprived C1 barrel, selectivity was largely maintained, with only a small shift in SI toward the SW D1 after regrowth (**Fig. 3d**). However, the number of responsive PV cells and evoked response magnitude were reduced across all trimmed whiskers and the spared D1 whisker compared to baseline (**Fig. 3e-f**). This suggests that during regrowth a reduction in the number of SW D1 responsive PV cells might be a compensatory response to restore neuronal whisker selectivity to the PW C1. On the other hand, in the spared D1 barrel, there was a strong, persistent shift in neuronal selectivity toward the PW D1 (**Fig. 3c**). The percentage of PW D1 responsive neurons and response magnitude to PW D1 stimulation were similar compared to baseline, but the number of SW C1 responsive neurons was strongly reduced (**Fig. 3e**). This suggests the persistent shift in selectivity is driven more strongly by reductions in responses to the trimmed whiskers rather than persistent potentiation of responses to the spared whisker. Reductions in PV activity may be necessary to facilitate cortical map restoration after whisker regrowth, but future manipulation studies will be required to test a potential causal role for PV activity in this process.

The recruitment of PV cells responsive to the spared whisker during whisker trimming could reflect either a passive homeostatic response to changes in the activity of local Pyr cells and/or bottom-up inputs. Alternatively, the recruitment of PV cells could contribute causally to experience-dependent cortical remapping, either by facilitating or limiting this process. In S1BF, brief optogenetic inhibition of PV cells increases whisker-evoked activity of Pyr cells, locally and in surround barrels^40,41^. However, few studies have reported how more prolonged changes in PV cell activity affect sensory evoked responses and experience-dependent plasticity. To test this, we expressed chemogenetic DREADDs in PV cells and recorded the acute effects on the activity of local PV, Pyr, and SST cells. Acute modulation of PV cells led to substantial changes in the number of whisker responsive Pyr cells (**Fig. 4d,g**). As expected, sensory evoked responses of Pyr cells were strongly reduced by activation of PV cells (**Fig. 4d**). In PV and SST cells, the number of whisker responsive cells and the magnitude of sensory evoked responses was also decreased (**Figs. 4d-e, 5c-d**). Inhibition of PV cells did not change the number of whisker responsive PV, Pyr, or SST cells but did significantly alter whisker-evoked responses, leading to earlier and larger peak activity with faster decay (**Figs. 4g-i, 5e-f**).

To better understand these counterintuitive results and their potential circuit mechanisms, we built a recurrent circuit model including Pyr, PV, SST, and VIP cells (**Fig. 6a**). Like ours, previous models of chemogenetic manipulation changed neuronal excitability^107^, sometimes with weakened inhibitory transmission^93^. Unlike ours, however, they had one excitatory and one inhibitory population^93,107^ or represented inhibition with a single parameter^108^, did not simulate sensory responses over time, and were compared with data qualitatively rather than fitted to them. Here, we fitted this circuit to the sensory response time courses of the manipulated PV neurons and of Pyr and SST neurons under both DREADDs. Our model shows why an evoked response cannot be read as a direct report of the manipulation. A DREADD alters the circuit before the stimulus arrives, so the same sensory input evokes a different trajectory of population activity (**Fig. 6d**). Under hM3D(Gq), depolarizing PV neurons reduced evoked responses in all three measured populations, including PV neurons, which still provided substantial inhibition. These reduced responses therefore do not require depolarization block, and complete PV silencing did not reproduce them.

The DREADDs also changed the variability of the calcium signals, which was largest under hM4D(Gi) and smallest under hM3D(Gq) in Pyr and PV neurons (**Fig. S9**). Such changes are easy to overlook when comparing only mean responses. We treated this variability as a circuit property and represented it as input variability, which sets how gradually a population responds near threshold. Under hM3D(Gq), tonic drive alone reduced the baseline-subtracted PV response while raising PV baseline firing (**Supplementary Table S7**). Adding the lower hM3D(Gq) variability returned PV baseline firing to near its Control level and lowered PV firing during the stimulus, so the fitted hM3D(Gq) suppression did not depend on baseline subtraction. This fall in PV firing resembles the paradoxical response of inhibition-stabilized networks^85,96^ but here also required the lower variability. With a stronger hM3D(Gq) manipulation, PV firing would rise again only as Pyr activity approached zero (**Fig. S15e**). Because the hM3D(Gq) effect, like the hM4D(Gi) effect, was already close to saturation at 3 mg/kg C21 (**Fig. 6e**), higher doses may not reach this regime.

Under hM4D(Gi), the model reproduced the early response peaks only when PV synaptic output was weakened along with PV excitability, within the model family and optimization conditions we tested. When we held PV output at its Control value and refit the model, the best fit had a comparable overall error but peaked at stimulus offset in every hM4D(Gi) trace (**Supplementary Table S9**). The early hM4D(Gi) peaks therefore point to a contribution from the presynaptic action of hM4D(Gi)^77,78^. Expanding the mean-field model into a sparse network with cell-to-cell heterogeneity and input noise preserved both the hM3D(Gq) suppression and the hM4D(Gi) early peaks. Although the network was not fitted to single neurons, its single-neuron response maps showed broad trends similar to those of the recordings (**Fig. S16**). The fitted model captures the direction and time course of the hM3D(Gq) effect but underestimates its full magnitude (Fig. S16c,d). This may reflect our minimal representation of hM3D(Gq), which consists only of increased tonic PV drive and reduced input variability. The model omits changes in PV firing pattern^109^ or inhibitory transmission^110^ reported in other circuits, and its linear calcium readout cannot capture a nonlinear, cell-dependent relation between firing and ΔF/F^111,112^; constraining these factors experimentally may close the remaining gap.

Our study has additional limitations. We used currently available genetically encoded calcium indicators, including GCaMP6^79^ and jGCaMP8^59^ to monitor neuronal activity. These indicators, with non-linear responses to changes in firing rates and imaged with framerates used in this study, cannot discriminate individual action potentials in fast-spiking PV cells. However, even without single-spike resolution, our work here, and multiple prior studies, demonstrate these indicators can reliably detect relative changes in neuronal activity in PV cells^113–117^. Investigating the spatial distribution of whisker-evoked responses in PV cells, we chose only select SWs (C1, D1, B2, and E3 for the C1 barrel). However, intracortical connectivity within S1BF is asymmetric - barrels within a row are more strongly interconnected than those across rows^118,119^. Additional experiments systematically comparing PV responses to SW stimuli within row or within arc will be needed to determine if these anatomical connections result in greater functional representation as predicted.

In our chronic whisker trimming experiment, we were only able to stimulate the spared whisker during trimming as we kept deprived whiskers trimmed flushed with the vibrissal pad. Stimulus evoked Pyr cell responses for deprived and spared whiskers are differentially affected by trimming^19,58^. We also observed such differential responses to whisker regrowth, so it is likely PV responses to the deprived whisker would be different compared to the spared whisker, but this remains speculative until experimentally tested. We focused on L2/3 in this study as supragranular layers maintain capacity for cortical plasticity into adulthood^16^. It is likely there may be other layer-specific changes in PV cell activity that contribute to experience-dependent plasticity. For example, following neonatal unilateral whisker trimming and regrowth whisker-evoked responses of L4 PV cells are reduced^120^ and the normal differential thalamocortical innervation between excitatory and PV cells is disrupted^121^. Morphological studies have also shown that whisker trimming in adults affects microglia^122^, myelination^123^, perineuronal nets^124–126^ and dendritic spine density^127,128^, highlighting the potential involvement of different cell types and complex homeostatic mechanisms in the barrel cortex after sensory deprivation. These questions will need to be explored further in future studies examining layer- and cell-type specific mechanisms regulating experience-dependent plasticity of PV cells.

Despite these limitations, our work reveals, for the first time, how PV cell activity changes dynamically during sensory evoked experience-dependent plasticity. PV cells responsive to the spared whisker are recruited over time in a spatially-specific manner, parallelling changes previously observed in Pyr cells. This recruitment can be reversed as sensory experience changes, though alterations in individual PV cell selectivity can persist for weeks after whisker regrowth. Our chemogenetic manipulation experiments demonstrate how changes in PV cell activity influence the activity of other local PV, Pyr, and SST cells. These changes can be difficult to predict, highlighting the reciprocal connectivity of PV and Pyr cells in local microcircuits and underscoring the importance of directly measuring the effects of chemogenetic manipulations on circuit activity. These experiments do not directly probe the causal role of PV cells in regulating experience-dependent changes in cortical sensory maps. However, the increased responsivity of PV cells to the spared whisker and strong suppression of sensory evoked responses in Pyr and SST cells upon activation of PV cells with DREADDs suggest a novel hypothesis – that PV-mediated inhibition could constrain experience-dependent plasticity and cortical sensory remapping. Reducing SST cell activity in deprived barrels with DREADDs has recently been shown to block whisker trimming induced map expansion of the spared whisker^129^. Recruitment of PV cells to the spared whisker during whisker trimming might thus constrain remapping both through direct increases in PV-mediated inhibition of Pyr cells, but also indirectly through inhibition of SST cells. Future experiments will be required to test these possibilities directly.

## Acknowledgements

We thank Ms. Brenda Vasquez, Ms. Callie Liu, Ms. Sravya Gadepalli, and Mr. Noe Cazares Jr. for assistance with mouse breeding and colony management. We thank Drs. Aye Theint Theint, Sammy Alhassen, Nazim Kourdougli, Anand Suresh, Michelle Wu, and Carlos Portera-Cailliau for helpful feedback, constructive criticism, and advice.

## Author Contributions

BC designed and performed research, analyzed data, and revised the manuscript; XY, IM and MD analyzed data and revised the manuscript; XY and MD developed and analyzed the computational model; WZ designed and performed research, analyzed data, and wrote the paper.

## Supplementary Figures

**Figure S1.**
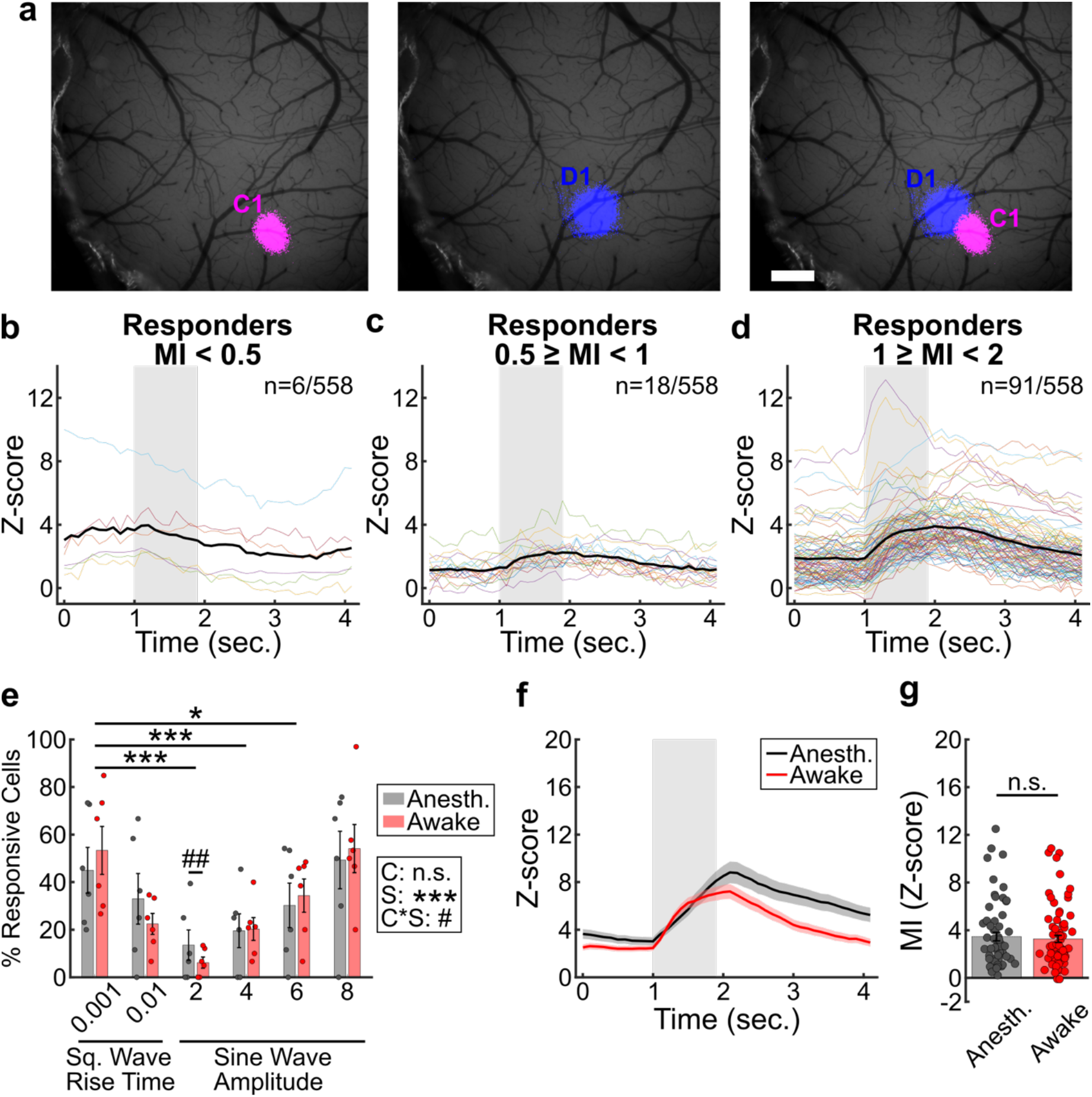
Whisker-evoked responses in PV cells are similar in awake and lightly anesthetized mice. **a.** Representative IOSI images of the C1 whisker cortical map (magenta, left), the D1 whisker cortical map (blue, middle) and the merged image of both the C1 and D1 maps (right). Scale bar=500 µm. **b-d.** Mean evoked calcium traces for PV cells classified as responsive with MI of < 0.5 (**b,** n=6/558 responsive cells), MI between 0.5 and 1 (**c**, n=18/558 responsive cells), or MI between 1 and 2 (**d**, n=91/558). A threshold of MI≧1 was chosen to consider cells as whisker responsive hereafter to minimize false positive classifications. **e.** Percentage of whisker responsive PV cells in the D1 barrel to PW D1 deflection in lightly anesthetized (gray) or awake (red) mice (n=6). Whisker deflection stimuli were delivered at 10 Hz for 1 second for a total of 20 trials using either square (Sq.) wave stimuli with 0.001 or 0.01 s rise times (amplitude of deflection of 3.35 and 3.07 +/- 0.01mm from peak to peak, respectively) or sine wave stimuli with voltage amplitudes of 2, 4, 6, or 8 V (amplitude of deflection of 2.70, 3.50, 4.50 and 5.00 +/- 0.01mm from peak to peak, respectively). GLME binomial model, ANOVA for fixed effects of anesthesia condition (C, *p*=0.09), stimulus type (S, ***, *p*<0.001), and condition*stimulus interaction (C*S, #, *p*<0.05), with significance for individual coefficients for stimulus type and condition*stimulus interactions, corrected using Benjamini and Hochberg’s method, indicated over corresponding data points (S: *, *p*<0.05; ***, *p*<0.001. C*S: ##, *p*<0.01). **f.** Mean evoked calcium trace for responder PV cells to square wave whisker deflections with 0.001 s rise time in lightly anesthetized (gray, n=61 cells) or awake (red, n=75 cells) mice. **g.** MI of responder PV cells to square wave whisker deflections with 0.001 s rise time in lightly anesthetized (gray, n=61 cells) or awake (red, n=75 cells) mice. LME model for the MI, fixed effect of condition (*p*=0.64).

**Figure S2.**
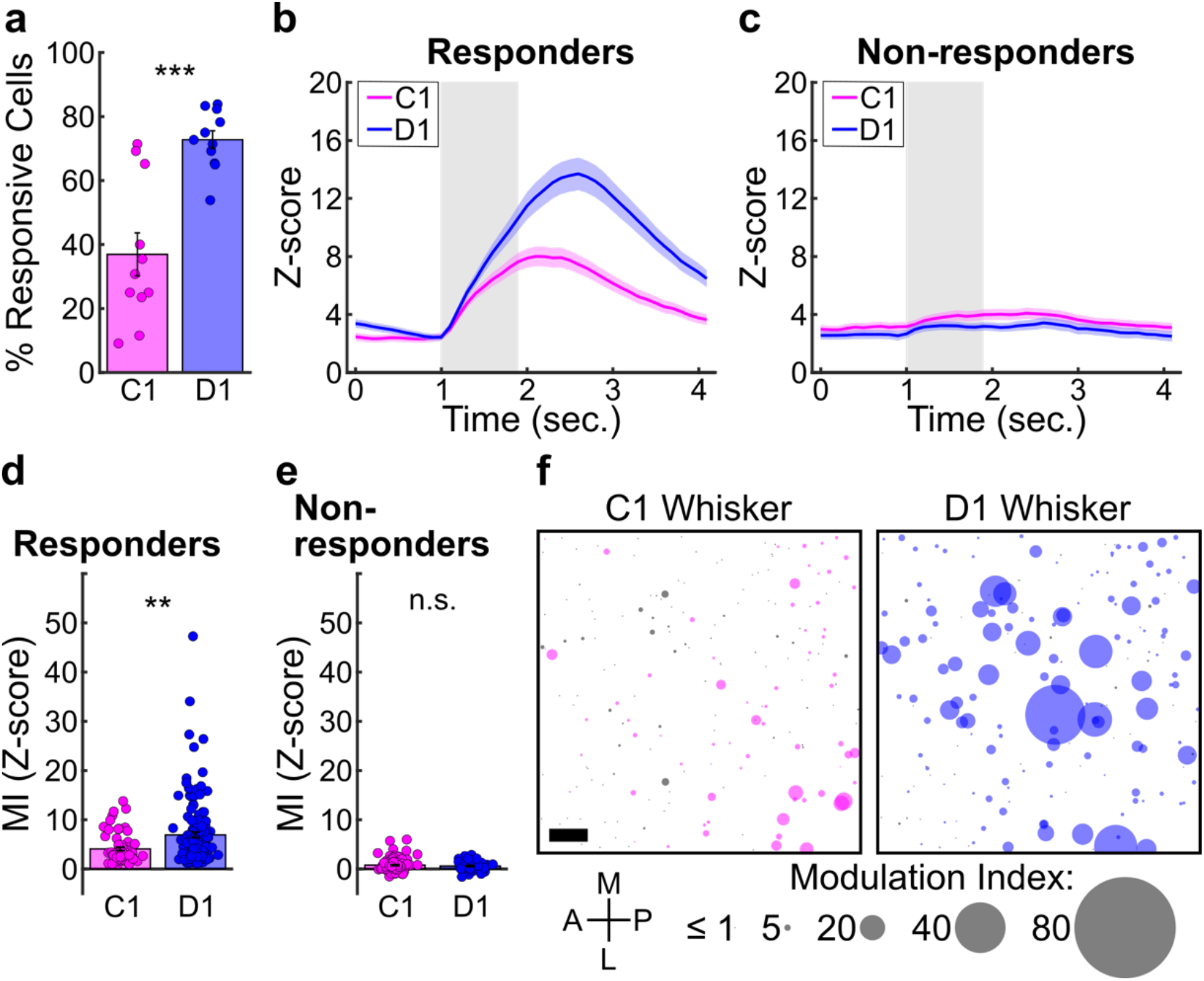
The spatial distribution of whisker-evoked responses of PV cells in the D1 barrel. **a.** Percentage of whisker responsive cells in the D1 barrel to the PW D1 and to the SW C1 (n=11 mice). GLME binomial model, fixed effect of whisker (***, *p*<0.001). **b.** Mean evoked calcium traces for responders (n=67 cells for SW C1, 139 cells for PW D1). **c.** Mean evoked calcium traces for non-responders (n=122 cells for SW C1, 50 cells for PW D1). **d.** MI calculated for responders (n=67 cells for SW C1, 139 cells for PW D1). LME model, fixed effect of whisker (**, *p*<0.01). **e.** MI calculated for non-responders (n=122 cells for SW C1, 50 cells for PW D1). LME model, fixed effect of whisker (*p*=0.47). **f.** Spatial distribution of all PV cells plotted according to relative position in the FOV centered on the D1 barrel. Responders are colored according to the indicated whisker; non-responders are colored in gray. Size of the circle corresponds to the MI of the cell for the indicated whisker. Scale bar=50 µm.

**Figure S3.**
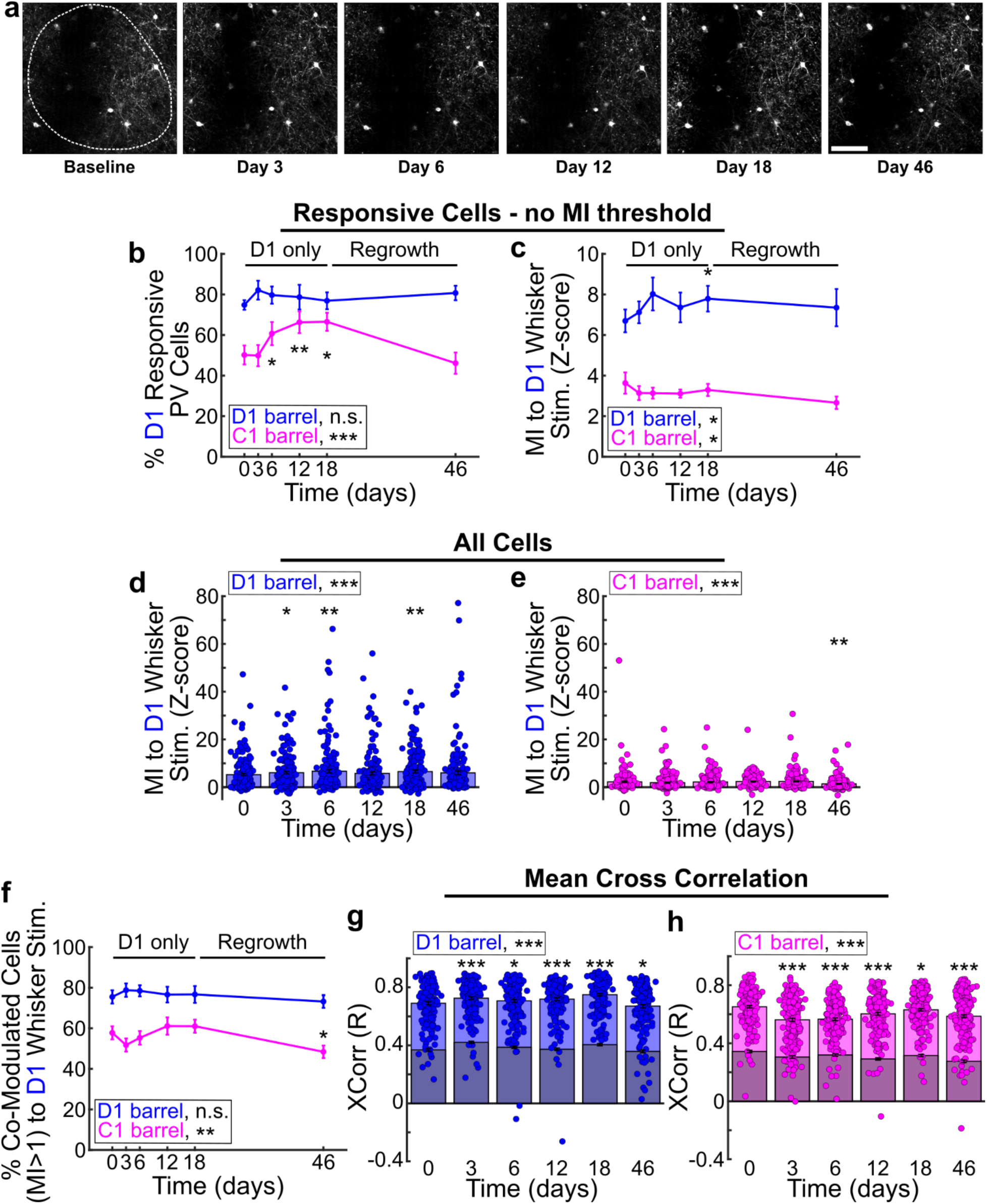
Longitudinal changes in whisker-evoked activity in PV cells during and after whisker trimming. **a.** Average projection 2P image of a representative FOV of trackable PV cells expressing GCaMP6s across imaging sessions during whisker deprivation and regrowth. Dotted outline corresponds to the functional IOSI map of the imaged barrel. Scale bar=100 µm. **b.** Percentage of PV cells classified as D1 whisker responsive (without applying the criteria of MI≧1) in the deprived C1 barrel (magenta) or spared D1 barrel (blue) over time (n=11 mice). GLME binomial model for the spared D1 barrel, fixed effect of timepoint (*p*=0.30). GLME binomial model for the deprived C1 barrel, fixed effect of timepoint (*p*<0.001). Significance for individual coefficients compared to day 0, corrected using Benjamini and Hochberg’s method, are indicated over corresponding data points (*, *p*<0.05; **, *p*<0.01). **c.** MI of PV cells classified as D1 whisker responsive (without applying the criteria of MI≧1) to D1 whisker stimulation in the deprived C1 barrel (magenta) or spared D1 barrel (blue) over time (n=144/109, 160/108, 155/133, 145/148, 149/139, and 151/88 at day 0, 3, 6, 12, 18, and 46 for the D1/C1 barrels, respectively). LME model for the spared D1 barrel, fixed effect of timepoint (*p*=0.04). LME model for the deprived C1 barrel, fixed effect of timepoint (*p*=0.03). Significance for individual coefficients compared to day 0, corrected using Benjamini and Hochberg’s method, are indicated over corresponding data points (*, *p*<0.05). **d-e.** MI of all cells in the spared D1 barrel (**d**) or deprived C1 barrel (**e**) to D1 whisker stimulation over time (n=189 cells for D1 and 210 cells for C1). LME model for the fixed effect of timepoint (***, *p*<0.001) for both **d** and **e**. LME model for the deprived C1 barrel, fixed effect of timepoint (*p*=0.03). Significance for individual coefficients compared to day 0, corrected using Benjamini and Hochberg’s method, are indicated over corresponding data points (*, *p*<0.05; **, *p*<0.01). **f.** Mean percentage of PV cells with MI≧1 on individual trials of D1 whisker stimulation in the deprived C1 barrel (magenta) or spared D1 barrel (blue) over time (n=11 mice). LME model for the spared D1 barrel, fixed effect of timepoint (*p*=0.32). LME model for the deprived C1 barrel, fixed effect of timepoint (*p*<0.01). Significance for individual coefficients compared to day 0, corrected using Benjamini and Hochberg’s method, are indicated over corresponding data points (*, *p*<0.05). **g-h.** Mean cross-correlation of individual PV cells with all other PV cells within the FOV in the spared D1 barrel (**g**) or deprived C1 barrel (**h**) across 20 trials of D1 whisker stimulation over time (n=189 cells for D1 and 210 cells for C1). LME model for the fixed effect of timepoint (***, *p*<0.001) for both **g** and **h**. Significance for individual coefficients compared to day 0, corrected using Benjamini and Hochberg’s method, are indicated over corresponding data points (*, *p*<0.05; ***, *p*<0.001). Overlaid gray bars indicate mean cross correlation values obtained using scrambled traces.

**Figure S4.**
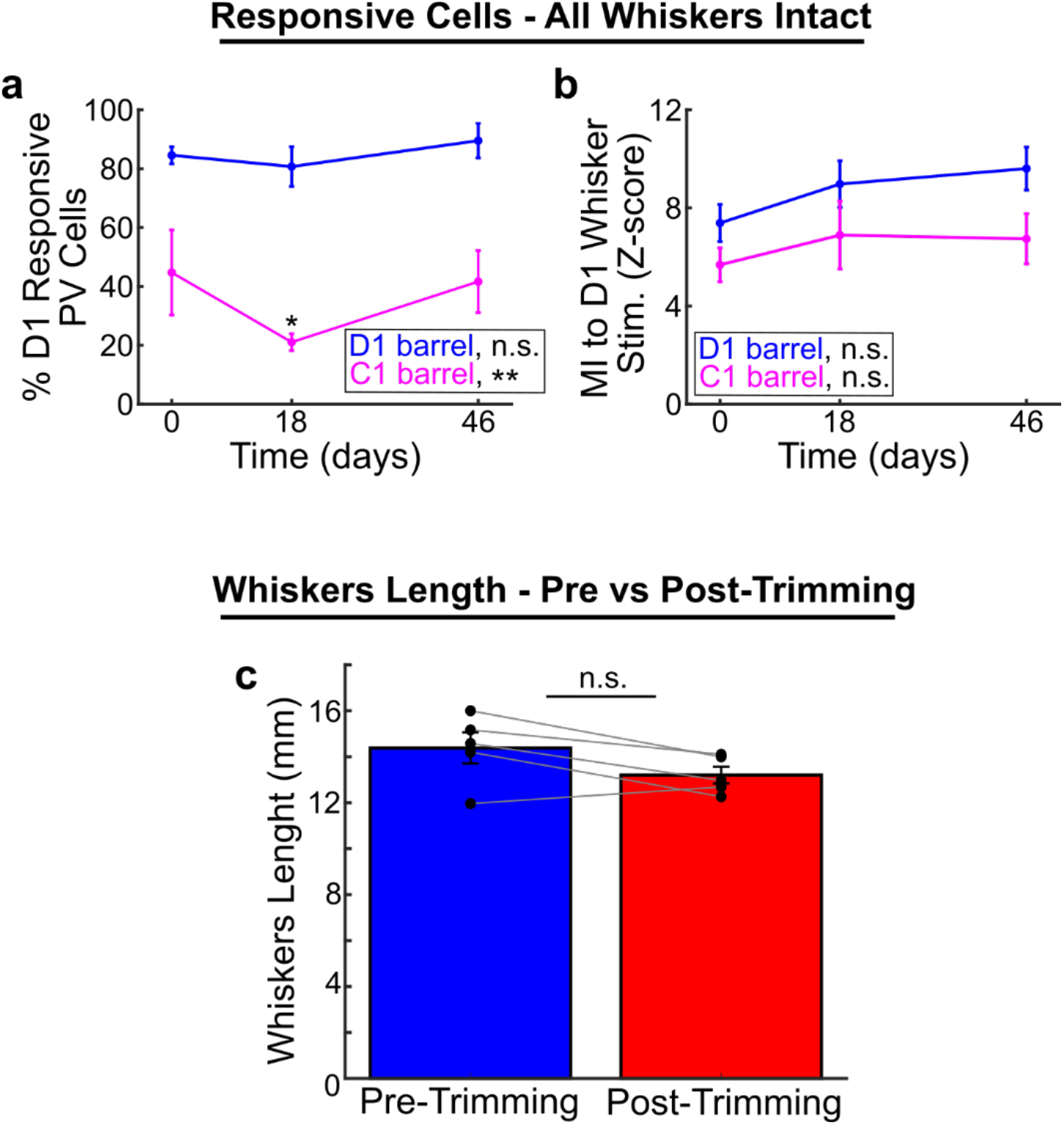
PV cell responses do not increase in control animals without whisker trimming. **a.** Percentage of D1 responsive PV cells in the D1 (blue) or C1 (magenta) barrel over time (n=5 mice). GLME binomial model for the D1 barrel, fixed effect of timepoint (*p*=0.29). GLME binomial model for the C1 barrel, fixed effect of timepoint (**, *p*<0.01). Significance for individual coefficients compared to day 0, corrected using Benjamini and Hochberg’s method, are indicated over corresponding data points (*, *p*<0.05). **b.** MI of D1 responsive PV cells in the D1 (blue) or C1 (magenta) barrel over time (n=92/49, 90/24, and 98/49 at day 0, 18, and 46 for the D1/C1 barrels, respectively). LME for the D1 barrel, fixed effect of timepoint (*p*=0.05). LME for the C1 barrel, fixed effect of timepoint (*p*=0.14). **c.** Averaged whisker length (in mm) pre- and post-trimming and regrowth (n=5 mice, 7 whiskers per mouse). Paired t-test (*p*=0.07).

**Figure S5.**
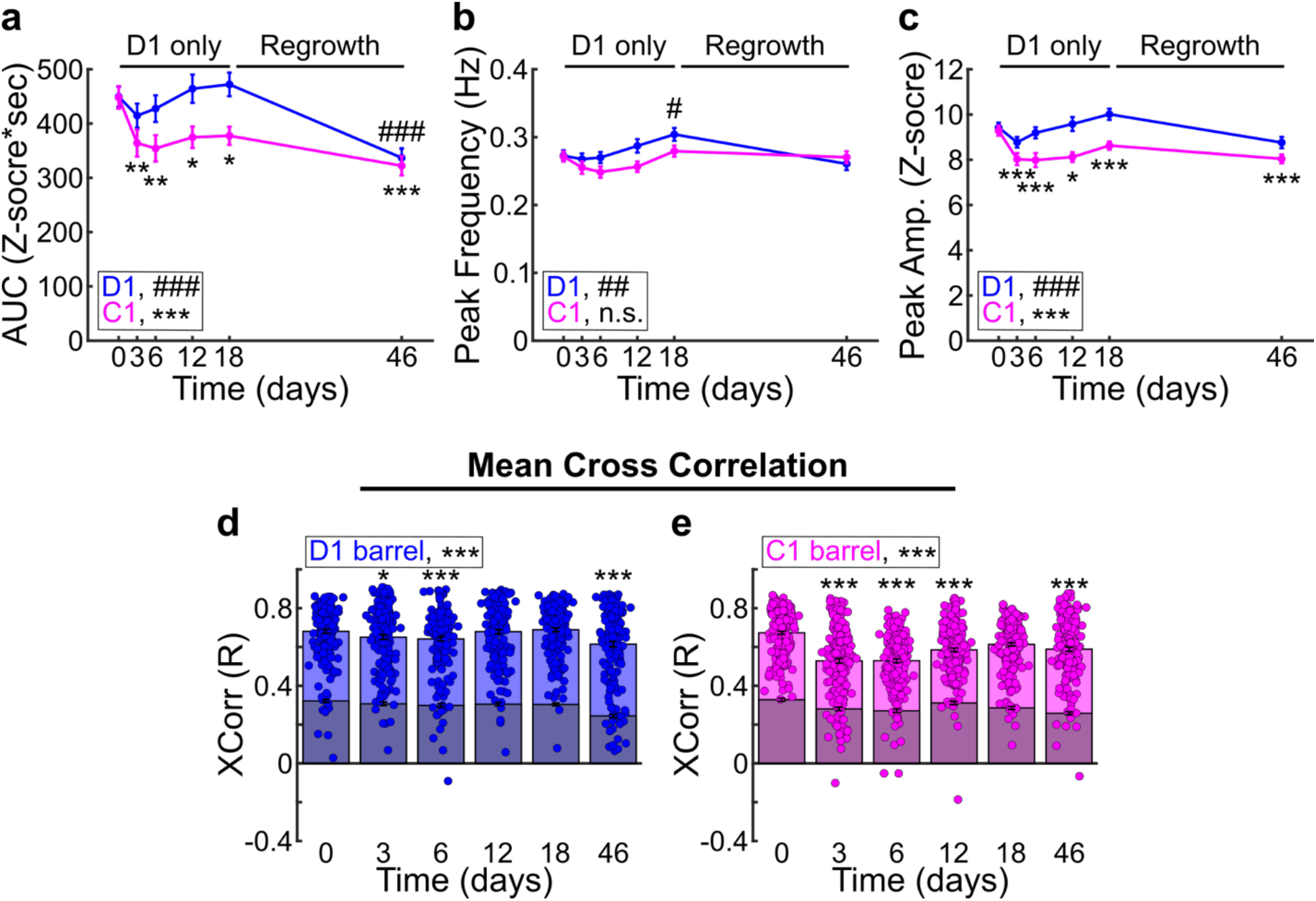
Spontaneous activity changes in PV cells during whisker trimming and regrowth. **a.** Total area-under-the-curve (AUC) of spontaneous activity (no whisker deflection) calcium traces from PV cells recorded in the D1 (blue) or C1 (magenta) barrels over time (n=189 cells for D1 and 210 cells for C1 from 11 mice). LME for the D1 barrel, fixed effect of timepoint (###, *p*<0.001). LME for the C1 barrel, fixed effect of timepoint (***, *p*<0.001). Significance for individual coefficients compared to day 0, corrected using Benjamini and Hochberg’s method, are indicated over corresponding data points (###, *p*<0.001; *, *p*<0.05; **, *p*<0.01; ***, *p*<0.001). **b.** Frequency of calcium transient peaks in the D1 (blue) or C1 (magenta) barrels over time (n=189 cells for D1 and 210 cells for C1 from 11 mice). LME for the D1 barrel, fixed effect of timepoint (##, *p*<0.01). LME for the C1 barrel, fixed effect of timepoint (*p*=0.06). Significance for individual coefficients compared to day 0, corrected using Benjamini and Hochberg’s method, are indicated over corresponding data points (#, *p*<0.05). **c.** Mean amplitude of calcium transient peaks in the D1 (blue) or C1 (magenta) barrels over time (n=189 cells for D1 and 210 cells for C1 from 11 mice). LME for the D1 barrel, fixed effect of timepoint (###, *p*<0.001). LME for the C1 barrel, fixed effect of timepoint (***, *p*=0.001). Significance for individual coefficients compared to day 0, corrected using Benjamini and Hochberg’s method, are indicated over corresponding data points (*, *p*<0.05; ***, *p*<0.001). **d-e.** Mean cross-correlation of individual PV cells with all other PV cells within the FOV in the spared D1 barrel (**d**) or deprived C1 barrel (**e**) across ∼150 s of spontaneous activity over time (n=189 cells for D1 and 210 cells for C1). LME model for the fixed effect of timepoint (***, *p*<0.001) for both **d** and **e**. Significance for individual coefficients compared to day 0, corrected using Benjamini and Hochberg’s method, are indicated over corresponding data points (*, *p*<0.05; ***, *p*<0.001). Overlaid gray bars indicate mean cross correlation values obtained using scrambled traces.

**Figure S6.**
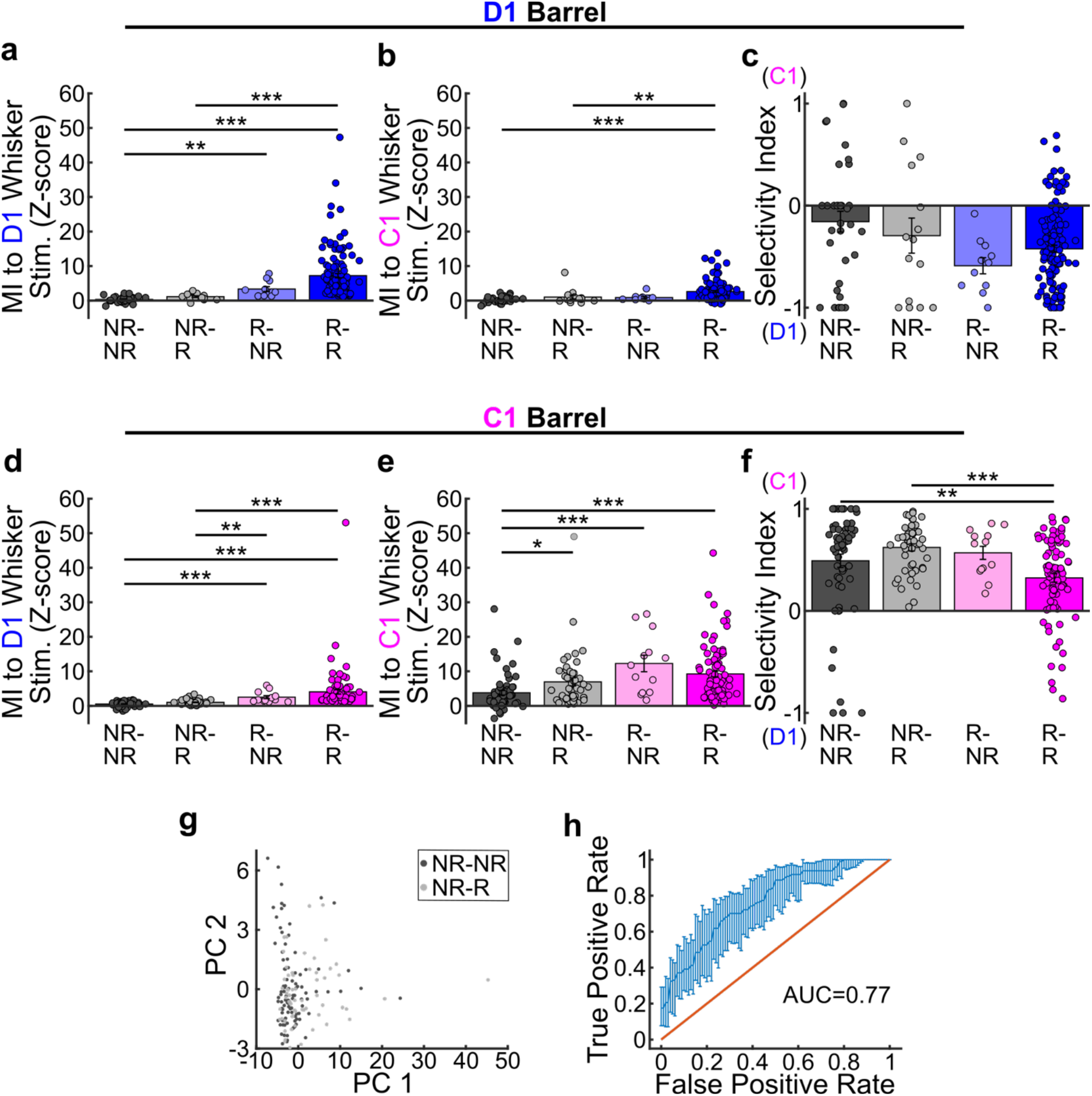
Baseline activity metrics only partially predict which PV cells will change responsivity to the spared D1 whisker during whisker trimming. **a.** MI of PV cells in the spared D1 barrel to the PW D1 at day 0. Cells are grouped by switch category: non-responsive to the PW D1 at day 0 and day 12 (NR-NR, n=35); switched from non-responsive to responsive (NR-R, n=15); switched from responsive to non-responsive (R-NR, n=11); or responsive on both day 0 and day 12 (R-R, n=128). Kruskal-Wallis test (**, *p*<0.01; ***, *p*<0.001). **b.** MI of PV cells in the spared D1 barrel to the SW C1 at day 0, by switch category (n=35, 15, 11, and 128 cells for NR-NR, NR-R, R-NR, and R-R, respectively). Kruskal-Wallis test (**, *p*<0.01; ***, *p*<0.001). **c.** Selectivity index of PV cells in the spared D1 barrel at day 0, by switch category (n=35, 15, 11, and 128 cells for NR-NR, NR-R, R-NR, and R-R, respectively). Kruskal-Wallis test. **d.** MI of PV cells in the deprived C1 barrel to the SW D1 at day 0, by switch category (n=62, 47, 13, and 88 cells for NR-NR, NR-R, R-NR, and R-R, respectively). Kruskal-Wallis test (**, *p*<0.01; ***, *p*<0.001). **e.** MI of PV cells in the derived C1 barrel to the PW C1 at day 0, by switch category (n=62, 47, 13, and 88 cells for NR-NR, NR-R, R-NR, and R-R, respectively). Kruskal-Wallis test (*, *p*<0.05; ***, *p*<0.001). **f.** Selectivity index of PV cells in the deprived C1 barrel at day 0, by switch category (n=62, 47, 13, and 88 cells for NR-NR, NR-R, R-NR, and R-R, respectively). Kruskal-Wallis test (**, *p*<0.01; ***, *p*<0.001). **g.** Plot of the first and second principal components from analysis of non-responder PV cells in both the spared D1 and deprived C1 barrels at day 0. Cells are group by switch category (NR-NR, dark gray; NR-R, light gray). **h.** Receiver-operating characteristic curve for a GLME model of switch category (NR-NR or NR-R) of day 0 non-responder PV cells (AUC=0.77). The data for PCA and the GLME model included barrel location, response magnitude to C1 and D1 whisker deflection, selectivity index, and the mean Z-score of the spontaneous activity trace for each.

**Figure S7.**
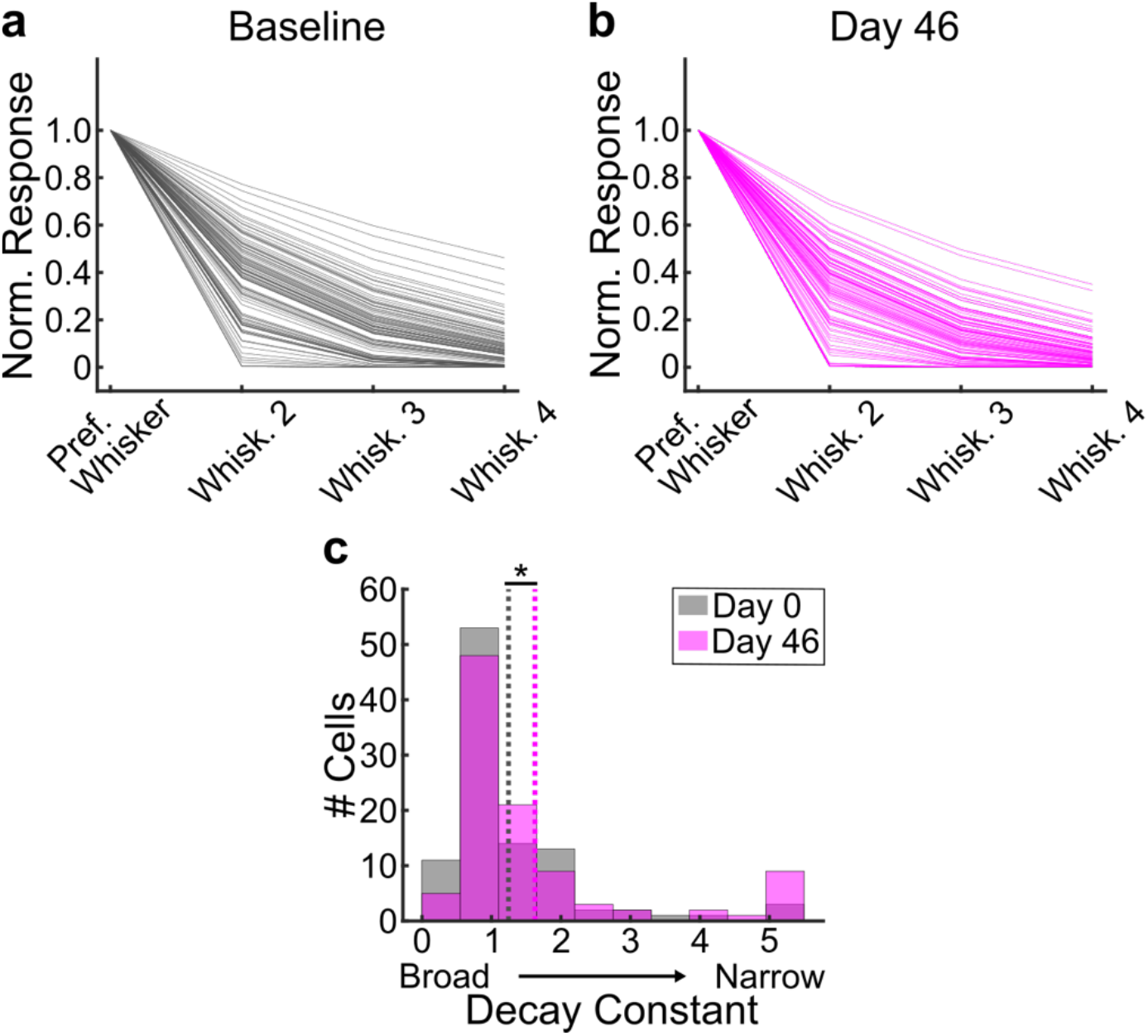
Changes in tuning width of PV cells before and after whisker trimming and regrowth. **a.** Tuning curves of individual PV cells (n=100 cells) in the C1 barrel to the PW C1, and SWs D1, B2, and E3 at baseline. Modulation indices for each whisker were normalized to the response magnitude of the preferred whisker (whisker with the highest MI), sorted in descending order, and fit with a single-term exponential decay. **b.** Tuning curves of individual PV cells (n=100 cells) in the C1 barrel at day 46 after whisker trimming and regrowth. **c.** Histogram of tuning curve decay constants from PV cells (n=100 cells) in the C1 barrel at day 0 (gray) or day 46 (magenta). LME model, fixed effect of timepoint (*, *p*<0.05).

**Figure S8.**
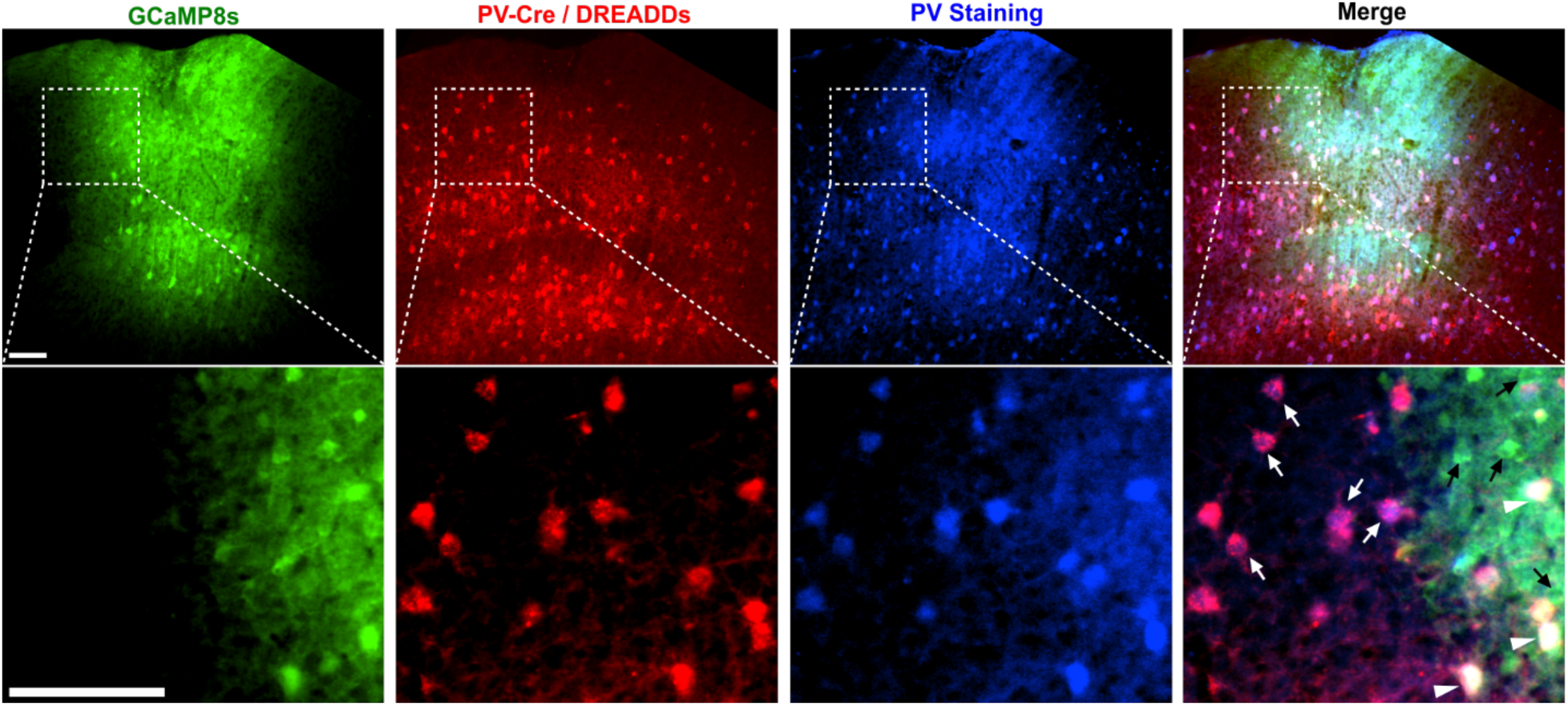
Cre-dependent expression of DREADDs in PV cells. Representative 10X images of PV immunostaining from a PV-Cre mouse injected with AAVs to express jGCaMP8s in all cells and Cre-dependent DREADDs in PV cells in the S1BF. Top row: jGCaMP8s expression (green), Cre-dependent expression of mCherry-tagged DREADDs in PV cells (red), PV immunostaining (blue) and the colocalization (merge). 70.83% of mCherry+ cells expressing DREADDs (red) colocalize with PV immunostaining (blue), and 80.62% of PV immunostained cells (blue) colocalize with mCherry fluorescence (red). n=4 mice, 3 slices per mouse. Scale bar=100µm. Bottom row: Higher magnification view of the regions indicated by squares from each image from the top row. White arrowheads indicate PV cells positive for mCherry-tagged DREADDs, jGCaMP8s and PV immunostaining. White arrows indicate PV cells positive for mCherry-tagged DREADDs and PV immunostaining, but lacking jGCaMP8s expression. Black arrows indicate expression of jGCaMP8s in putative Pyr cells lacking both mCherry tagged DREADDs and PV immunostaining. Scale bar=100µm.

**Figure S9.**
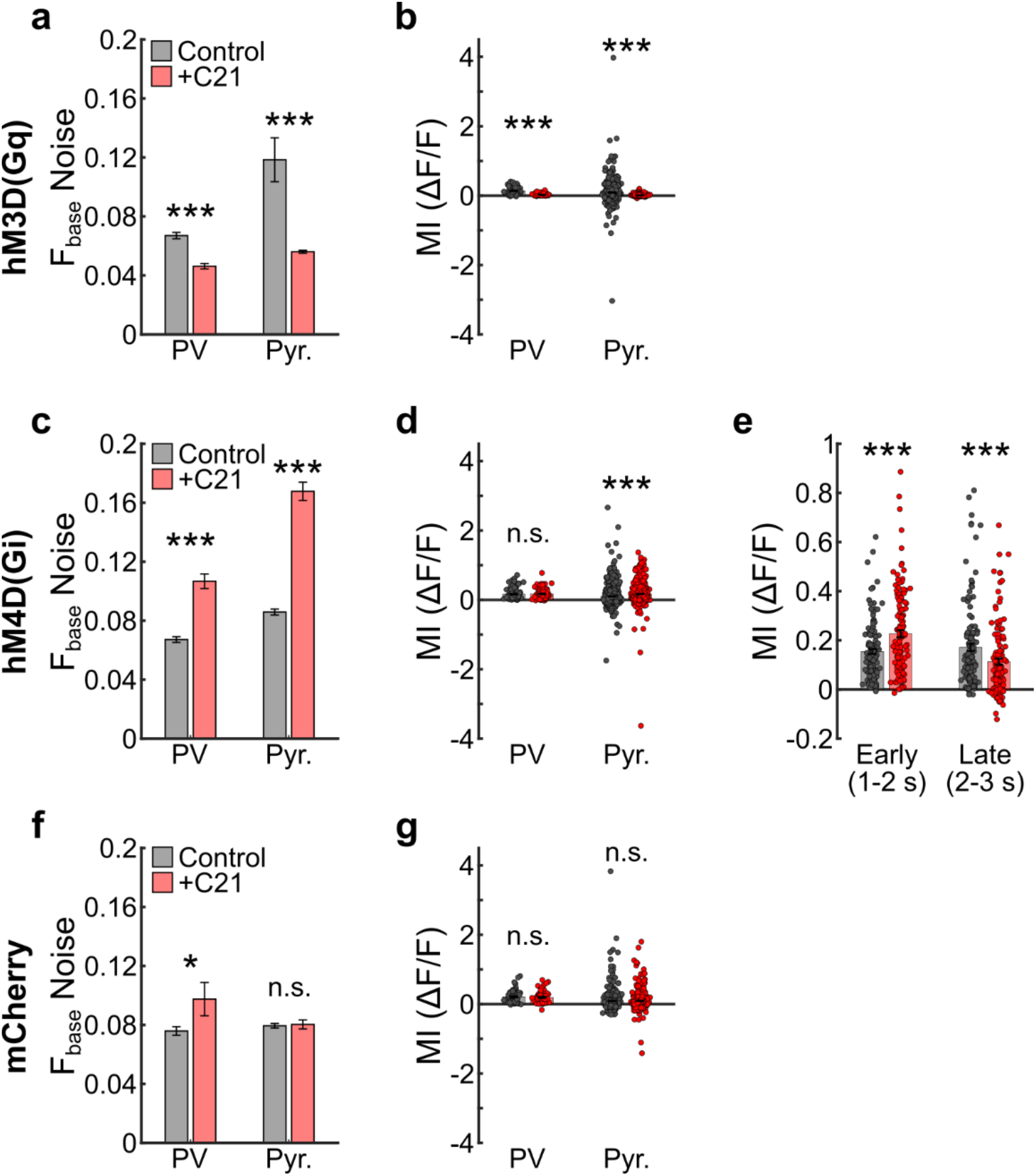
Chemogenetic modulation of PV cells affects the noise of baseline fluorescence and whisker-evoked responses in PV and Pyr cells. **a.** Noise of baseline fluorescence in PV and Pyr cells in mice (n=7) expressing hM3D(Gq) in PV cells before and after C21 (5 mg/kg, i.p.). LME model, fixed effect of condition (before and after C21; PV, ***, *p*<0.001; Pyr, ***, *p*<0.001). **b.** Mean evoked ΔF/F MI for all PV and Pyr cells in mice expressing hM3D(Gq) in PV cells before and after C21 (n=92/92 PV and 813/667 Pyr cells before/after C21). LME models for the MI (Time=1-3 s), fixed effect of condition (PV, ***, *p*<0.001; Pyr, ***, *p*<0.001). **c.** Noise of baseline fluorescence in PV and Pyr cells in mice (n=8) expressing hM4D(Gi) in PV cells before and after C21. LME model, fixed effect of condition (PV, ***, *p*<0.001; Pyr, ***, *p*<0.001). **d.** Mean evoked ΔF/F MI for all PV and Pyr cells in mice expressing hM4D(Gi) in PV cells before and after C21 (n=132/132 PV and 1113/979 Pyr cells before/after C21). LME models for the MI (Time=1-3 s), fixed effect of condition (PV, *p*=0.51; Pyr, ***, *p*<0.001). **e.** Mean evoked ΔF/F MI for all PV cells in mice expressing hM4D(Gi) in PV cells before and after C21 (n=132 cells) early (Time=1-2 s) and late (Time=2-3 s) after onset of whisker stimulation. LME models for the MI, fixed effect of condition (Early, ***, *p*<0.001; Late, ***, *p*<0.001). **f.** Noise of baseline fluorescence in PV and Pyr cells in mice (n=6) expressing mCherry in PV cells before and after C21. LME model, fixed effect of condition (PV, *, *p*<0.05; Pyr, *p*=0.78). **g.** Mean evoked ΔF/F MI for all PV and Pyr cells in mice expressing mCherry in PV cells before and after C21 (n=71/71 PV and 598/617 Pyr cells before/after C21). LME models for the MI (Time=1-3 s), fixed effect of condition (PV, *p*=0.21; Pyr, *p*=0.75).

**Figure S10.**
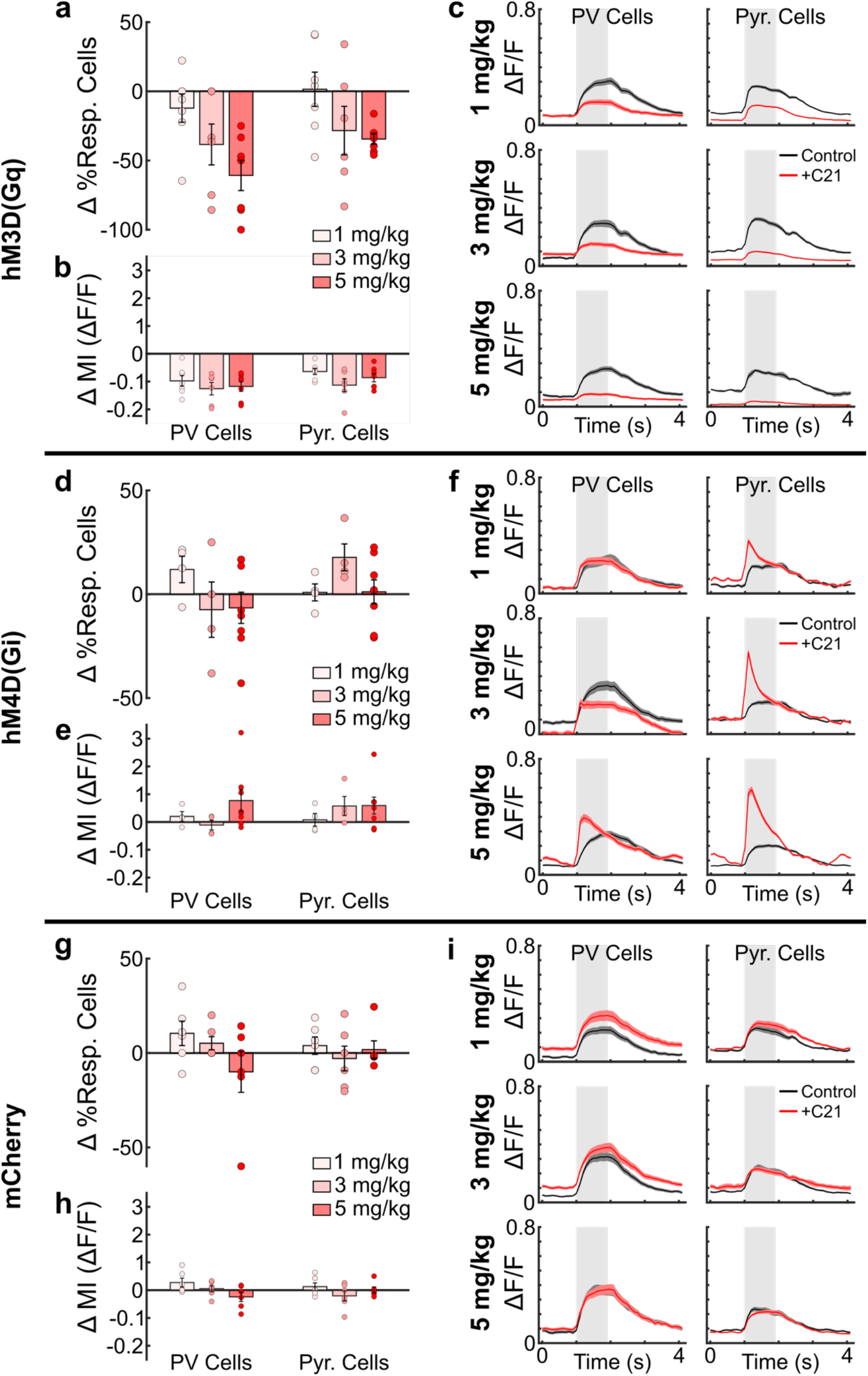
Dose responses of chemogenetic modulation of PV cells on whisker-evoked activity in local PV and Pyr cells. **a.** Change in the percentage of PW responsive PV and Pyr cells in mice (n=7) expressing hM3D(Gq) in PV cells before and after different doses of C21 (1, 3, and 5 mg/kg, i.p.). **b.** Change in the evoked ΔF/F MI of PV and Pyr cells in mice expressing hM3D(Gq) in PV cells before and after different doses of C21. **c.** Mean evoked ΔF/F calcium traces of PV and Pyr cells in mice expressing hM3D(Gq) in PV cells before and after different doses of C21. **d.** Change in the percentage of PW responsive PV and Pyr cells in mice (n=8) expressing hM4D(Gi) in PV cells before and after different doses of C21 (1, 3, and 5 mg/kg, i.p.). **e.** Change in the early (1-2 s) evoked ΔF/F MI of PV and Pyr cells in mice expressing hM4D(Gi) in PV cells before and after different doses of C21. **f.** Mean evoked ΔF/F calcium traces of PV and Pyr cells in mice expressing hM4D(Gi) in PV cells before and after different doses of C21. **g.** Change in the percentage of PW responsive PV and Pyr cells in mice (n=6) expressing mCherry in PV cells before and after different doses of C21 (1, 3, and 5 mg/kg, i.p.). **h.** Change in the evoked ΔF/F MI of PV and Pyr cells in mice expressing mCherry in PV cells before and after different doses of C21. **i.** Mean evoked ΔF/F calcium traces of PV and Pyr cells in mice expressing mCherry in PV cells before and after different doses of C21.

**Figure S11.**
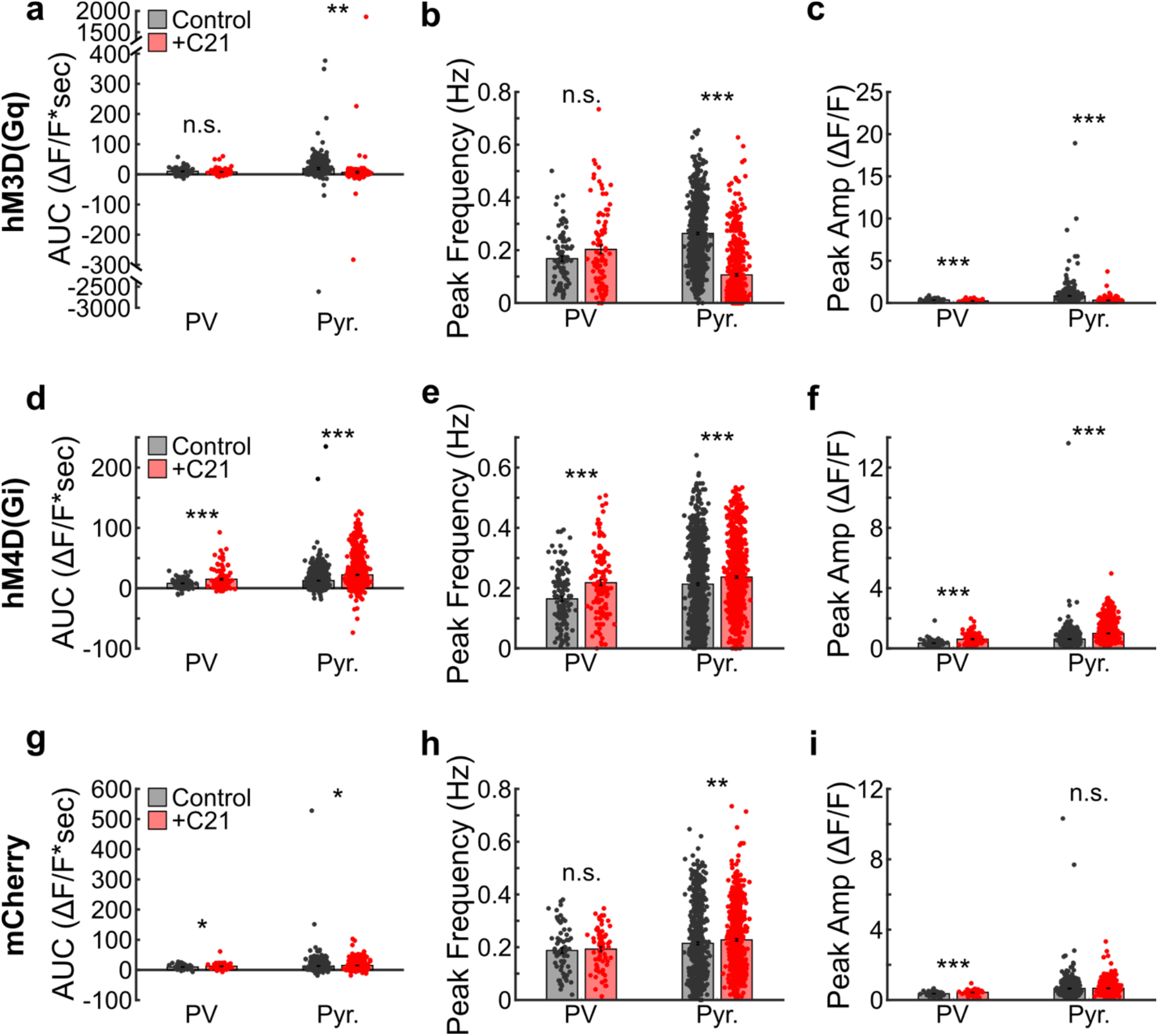
Acute chemogenetic modulation of PV cells affects spontaneous activity of local PV and Pyr cells. **a.** Total area-under-the-curve (AUC) of spontaneous activity (no whisker deflection) ΔF/F calcium traces from PV and Pyr cells in mice expressing hM3D(Gq) in PV cells before and after C21 (5 mg/kg, i.p.; n=92/92 PV and 774/638 Pyr cells before and after C21, respectively, from 7 mice). LME models for fixed effect of timepoint (PV, *p*=0.15; Pyr, **, *p*<0.01). **b.** Frequency of ΔF/F calcium transient peaks from PV and Pyr cells before and after C21. LME models for fixed effect of timepoint (PV, *p*=0.06; Pyr, ***, *p*<0.001). **c.** Mean amplitude of ΔF/F calcium transient peaks from PV and Pyr cells before and after C21. LME models for fixed effect of timepoint (PV, ***, *p*<0.001; Pyr cells, ***, *p*<0.001). **d.** Total AUC of spontaneous activity ΔF/F calcium traces from PV and Pyr cells in mice expressing hM4D(Gi) in PV cells before and after C21 (n=132/132 PV and 1017/910 Pyr cells before and after C21, respectively, from 8 mice). LME models for fixed effect of timepoint (PV, ***, *p*<0.001; Pyr, ***, *p*<0.001). **e.** Frequency of ΔF/F calcium transient peaks from PV and Pyr cells before and after C21. LME models for fixed effect of timepoint (PV, ***, *p*<0.001; Pyr, ***, *p*<0.001). **f.** Mean amplitude of ΔF/F calcium transient peaks from PV and Pyr cells before and after C21. LME models for fixed effect of timepoint (PV, ***, *p*<0.001; Pyr cells, ***, *p*<0.001). **g.** Total AUC of spontaneous activity ΔF/F calcium traces from PV and Pyr cells in mice expressing mCherry in PV cells before and after C21 (n=71/71 PV and 596/673 Pyr cells before and after C21, respectively, from 6 mice). LME models for fixed effect of timepoint (PV, *, *p*<0.05; Pyr, *, *p*<0.05). **h.** Frequency of ΔF/F calcium transient peaks from PV and Pyr cells before and after C21. LME models for fixed effect of timepoint (PV, *p*=0.67; Pyr, **, *p*<0.01). **i.** Mean amplitude of ΔF/F calcium transient peaks from PV and Pyr cells before and after C21. LME models for fixed effect of timepoint (PV, ***, *p*<0.001; Pyr, *p*=0.43).

**Figure S12.**
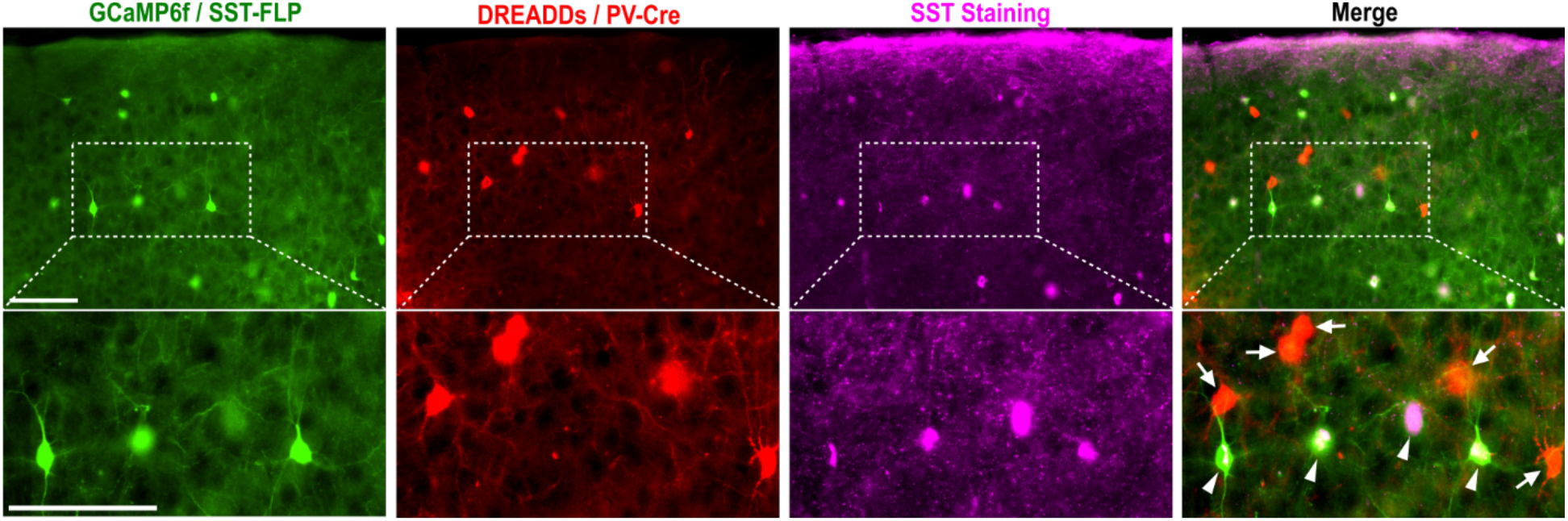
Flp-dependent expression of GCaMP6f in SST cells and Cre-dependent expression of DREADDs in PV cells. Representative 20X images of SST immunostaining from a double transgenic PV-Cre:SST-Flp mouse injected with AAVs to express GCaMP6f Coff/Fon and Cre-dependent mCherry-tagged DREADDs in the S1BF. Top row: Flp-dependent expression of GCaMP6fs only in SST cells (green), Cre-dependent expression of DREADDs only in PV cells (red), SST immunostaining (magenta) and the colocalization (merge). Scale bar=100 µm. Bottom row: Higher magnification view of the regions indicated by rectangles from each image from the top row. White arrowheads indicate SST cells positive for GCaMP6f and SST immunostaining. White arrows indicate PV cells positive for mCherry-tagged DREADDs and negative for SST immunostaining. Scale bar=100 µm.

**Figure S13.**
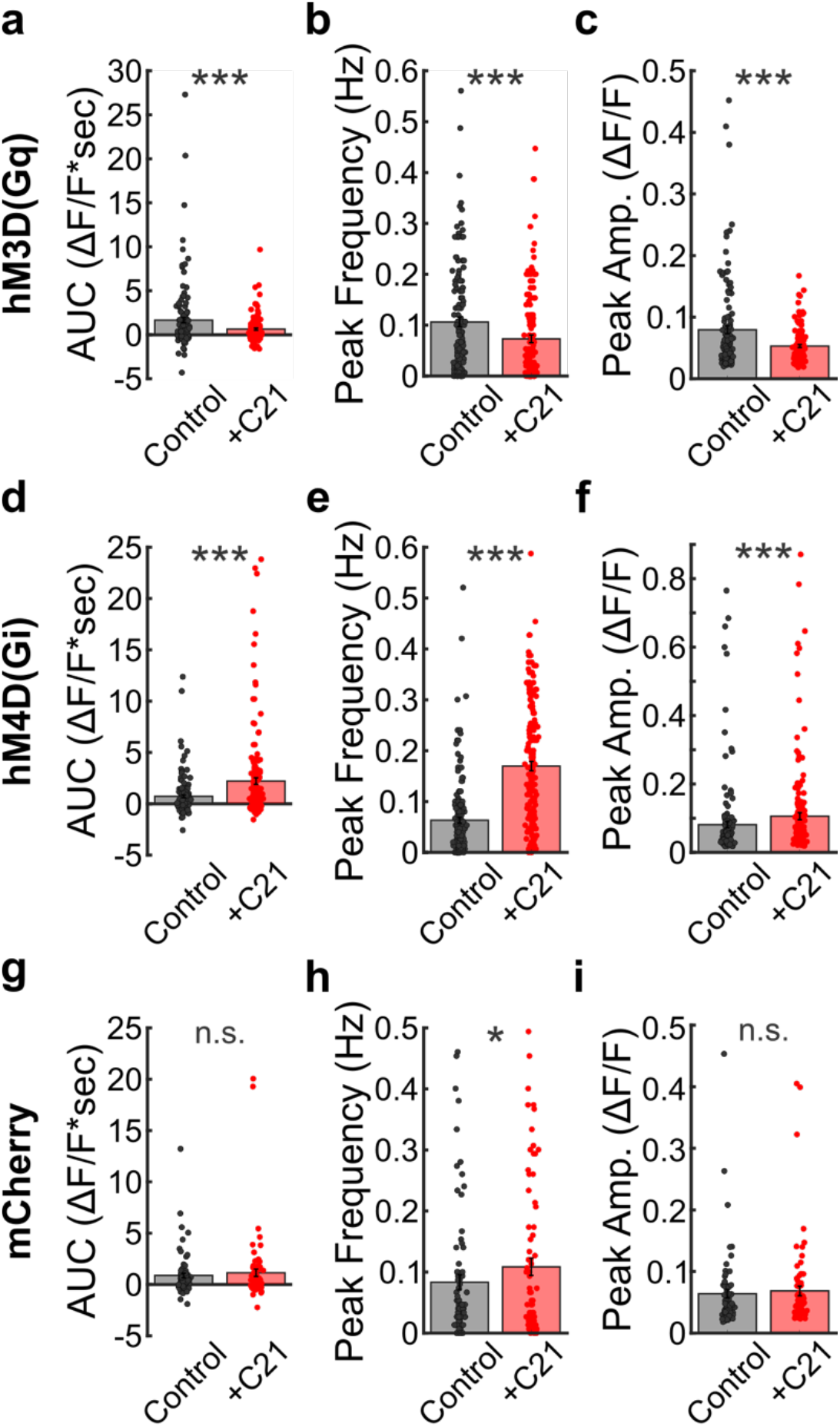
Acute chemogenetic modulation of PV cells affects spontaneous activity of local SST cells. **a.** Total AUC of spontaneous activity (no whisker deflection) ΔF/F calcium traces from SST cells in mice expressing hM3D(Gq) in PV cells before and after C21 (n=137 cells from 14 mice). LME model, fixed effect of timepoint (***, *p*<0.001). **b.** Frequency of ΔF/F calcium transient peaks before and after C21. LME model, fixed effect of timepoint (***, *p*<0.001). **c.** Mean amplitude of ΔF/F calcium transient peaks before and after. LME model, fixed effect of timepoint (***, *p*<0.001). **d.** Total AUC of spontaneous activity ΔF/F calcium traces from SST cells in mice expressing hM4D(Gi) in PV cells before and after C21 (n=173 cells from 16 mice). LME model, fixed effect of timepoint (***, *p*<0.001). **e.** Frequency of ΔF/F calcium transient peaks before and after C21. LME model, fixed effect of timepoint (***, *p*<0.001). **f.** Mean amplitude of ΔF/F calcium transient peaks before and after C21. LME model, fixed effect of timepoint (***, *p*<0.001). **g.** Total AUC of spontaneous activity ΔF/F calcium traces from SST cells in mice expressing mCherry in PV cells before and after C21 (n=80 cells from 12 mice). LME model, fixed effect of timepoint (*p*=0.31). **h.** Frequency of ΔF/F calcium transient peaks before and after C21. LME model, fixed effect of timepoint (*, *p*<0.05). **i.** Mean amplitude of ΔF/F calcium transient peaks before and after C21. LME model, fixed effect of timepoint (*p*=0.42).

**Figure S14.**
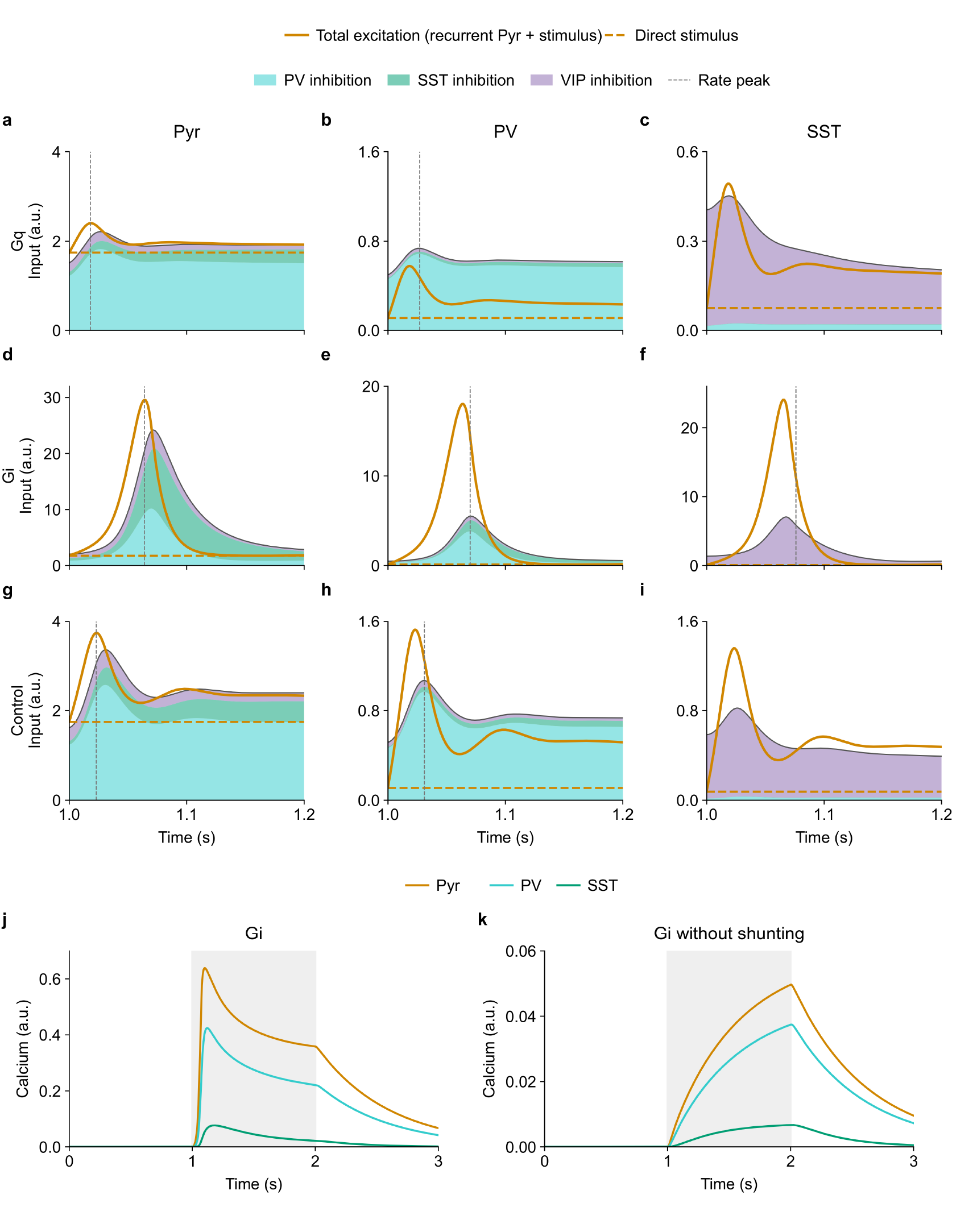
Reduced PV output efficacy lets recurrent excitation outpace inhibition and is necessary for the early hM4D(Gi) calcium peaks. All panels show deterministic simulations of the fitted four-population mean-field model. **a–i.** Inputs to Pyr, PV and SST (columns) under hM3D(Gq) (a–c), hM4D(Gi) (d–f) and Control (g–i). Orange solid lines show recurrent Pyr excitation plus direct stimulus input; orange dashed lines show the direct stimulus component in the same units. Stacked cyan, green and purple areas show the magnitudes of PV, SST and VIP inhibition, respectively; their upper outline gives total recurrent inhibition. Pyr→PV depression and Pyr→SST facilitation are included. Background drive, PV tonic offsets and Pyr adaptation are omitted. Gray vertical dashed lines mark the time of the stimulus-period firing-rate peak when it falls within the displayed window. **j,k.** Baseline-subtracted Pyr, PV and SST calcium responses in fitted hM4D(Gi) and after restoring κ_PV_^(Gi)^ to 1 (hM4D(Gi) without shunting), respectively. All other hM4D(Gi) parameters and the calcium readout are unchanged; each condition starts at its own equilibrium baseline. Stimulus onset is at 1 s in all panels. Panels a–i show the first 0.2 s after onset, and j,k show the 1–2-s stimulus period (gray shading) with one second before and after stimulation.

**Figure S15.**
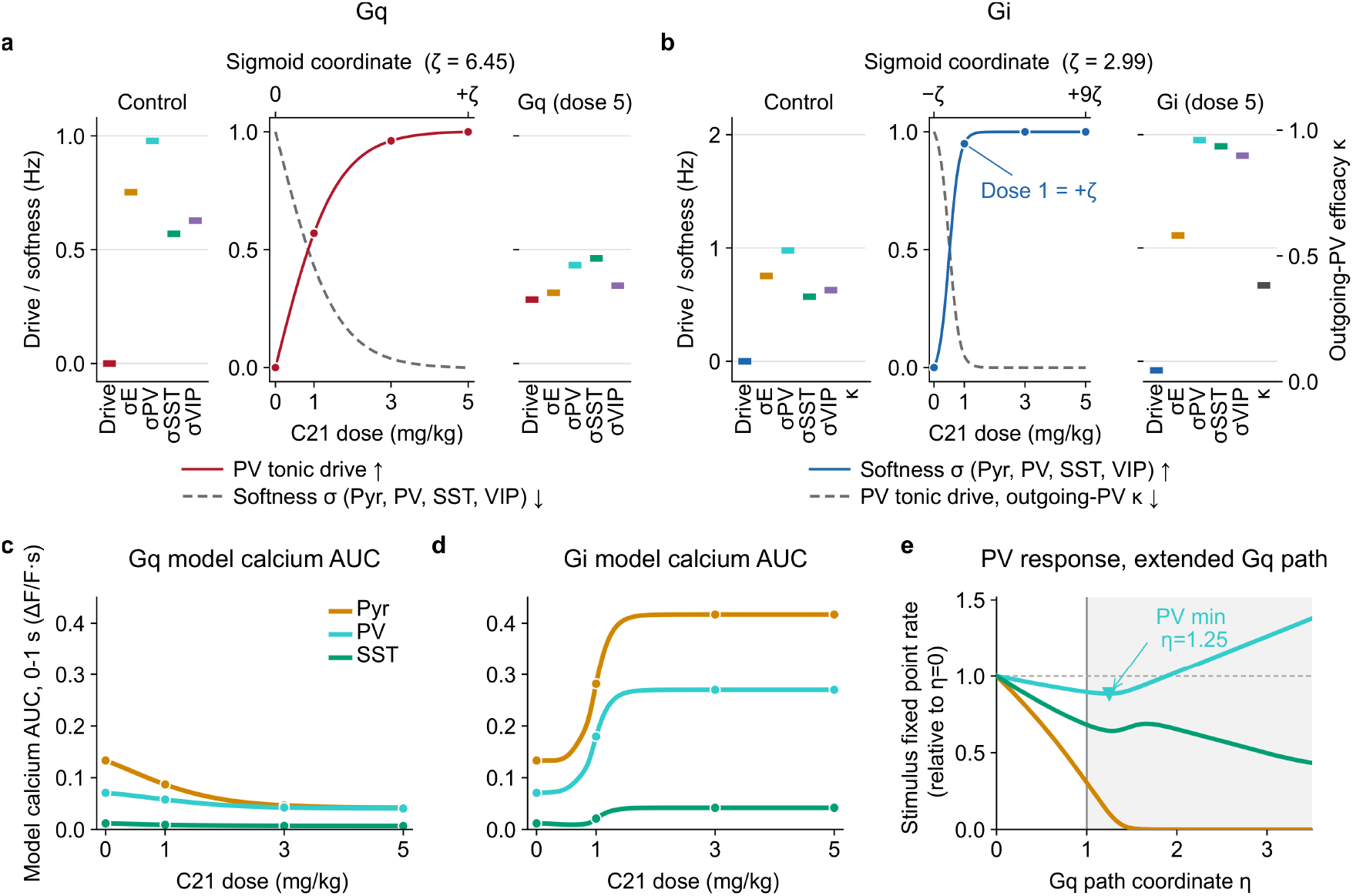
Calibrated dose paths capture the opposing dose dependence of hM3D(Gq) and hM4D(Gi) responses, and extending the hM3D(Gq) path predicts a nonmonotonic PV response. **a, b.** Joint parameter progression for hM3D(Gq) (a) and hM4D(Gi) (b) at C21 doses d=0, 1, 3 and 5 mg/kg. Central plots show the shared sigmoid progress (solid) and its complement (dashed). Side plots show Control and dose-5 values: Drive denotes 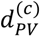, σ denotes 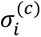, and the outgoing-PV label denotes 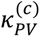. The fitted spans were 6.45 for hM3D(Gq) and 2.99 for hM4D(Gi). **c, d.** Baseline-subtracted model-calcium AUC over 0–1 s after stimulus onset along the hM3D(Gq) (c) and hM4D(Gi) (d) paths. Intermediate-dose SST values are model predictions. **e.** Stimulus-period fixed-point firing rates, normalized to *η* = 0, along the extended joint change in PV tonic input and transfer softness. Pyr, PV and SST are colored as in c. Softness follows the smooth continuation defined in Supplementary Information (Eq. S5), which closely approximates Control at *η* = 0 and the fitted dose-5 state at *η* = 1. Shading marks the uncalibrated region *η* > 1. PV activity first decreases and then increases as Pyr activity approaches zero.

**Figure S16.**
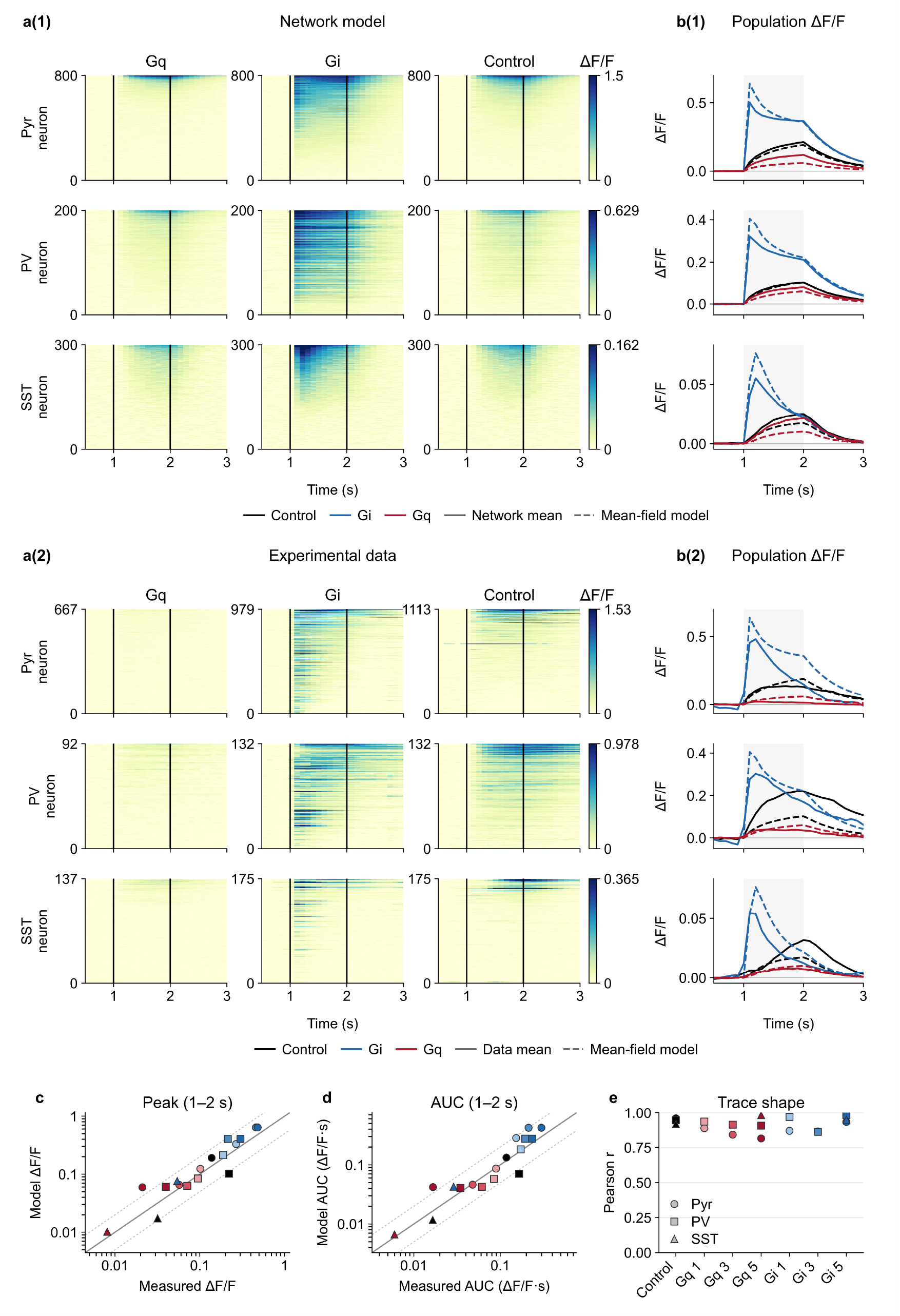
A heterogeneous network model preserves the population-level dynamics of the mean-field model. **a(1), a(2).** Trial-averaged, baseline-subtracted calcium heat maps for the heterogeneous network model (a(1)) and experimental recordings (a(2)), with Pyr, PV and SST in rows and hM3D(Gq), hM4D(Gi) and Control in columns. Baselines were the mean signal over the 10 s (network) or 1 s (experiment) before stimulus onset. Network-model panels contain all 800 Pyr, 200 PV and 300 SST units, averaged over 20 trials per condition. Experimental panels contain all recorded cells from the dose-5 cohorts, without responsiveness selection: Control is the hM4D(Gi)-group pre-C21 baseline (BL) session; hM4D(Gi) and hM3D(Gq) are the C21 sessions of the corresponding groups. Other control recordings showed broadly similar temporal profiles. Experimental cell/mouse counts in hM3D(Gq), hM4D(Gi) and Control, respectively, were 667/7, 979/8 and 1113/8 for Pyr; 92/7, 132/8 and 132/8 for PV; and 137/14, 175/16 and 175/16 for SST. Rows do not identify matched cells across conditions. Cells were ranked independently within each population and condition by their modulation index, defined for recordings in Methods (Two-photon Calcium Imaging) and for model units as the mean calcium increment over 0–1 s after stimulus onset divided by the prestimulus standard deviation. Color bars show ΔF/F. The linear color scale was shared across conditions within each population, with its upper limit set to the maximum of the three condition-specific 99.5th percentiles over displayed cell-by-time values. Limits were set separately for model and experiment. Negative values were clipped for display only and retained in quantitative analyses. **b(1), b(2).** All-cell population means from the network model (b(1), solid) and experimental recordings (b(2), solid), each compared with the same fitted mean-field reference (dashed). Experimental means correspond to the fitting targets. Population means weighted cells equally, pooling experimental cells across mice. Black, blue and red denote Control, hM4D(Gi) and hM3D(Gq). Vertical lines in a and gray shading in b mark the stimulus (1–2 s; onset at 1 s). **c, d.** Measured versus mean-field model peak ΔF/F (c) and AUC (ΔF/F·s; d) during the 1–2-s stimulus for 17 population-by-condition/dose pairs. Shape indicates population; color indicates condition and dose. Solid and dotted lines indicate equality and a twofold difference, respectively. **e.** Pearson correlations between measured and mean-field model traces over 0–3 s after stimulus onset (1–4 s in the figure coordinates). Comparisons in c–e use data that were also used for endpoint fitting or dose calibration.

## References

1. Hooks, B. M. & Chen, C. Circuitry Underlying Experience-Dependent Plasticity in the Mouse Visual System. Neuron 106, 21–36 (2020).

2. Barth, A. L. & Ray, A. Progressive Circuit Changes during Learning and Disease . Neuron 104, 37–46 (2019).

3. Chéreau, R., Williams, L. E., Bawa, T. & Holtmaat, A. Circuit mechanisms for cortical plasticity and learning. Semin Cell Dev Biol 125, 68–75 (2022).

4. Hübener, M. & Bonhoeffer, T. Neuronal plasticity: beyond the critical period. Cell 159, 727–737 (2014).

5. Diamond, M. E. & Arabzadeh, E. Whisker sensory system - from receptor to decision. Prog. Neurobiol. 103, 28–40 (2013).

6. Woolsey, T. A. & Van der Loos, H. The structural organization of layer IV in the somatosensory region (SI) of mouse cerebral cortex. The description of a cortical field composed of discrete cytoarchitectonic units. Brain Res 17, 205–242 (1970).

7. Petersen, C. C. H. Sensorimotor processing in the rodent barrel cortex. Nat. Rev. Neurosci. 20, 533–546 (2019).

8. Oberlaender, M. et al. Cell type-specific three-dimensional structure of thalamocortical circuits in a column of rat vibrissal cortex. Cereb Cortex 22, 2375–2391 (2012).

9. Petersen, C. C. H., Grinvald, A. & Sakmann, B. Spatiotemporal dynamics of sensory responses in layer 2/3 of rat barrel cortex measured in vivo by voltage-sensitive dye imaging combined with whole-cell voltage recordings and neuron reconstructions. J Neurosci 23, 1298–1309 (2003).

10. Narayanan, R. T. et al. Beyond Columnar Organization: Cell Type- and Target Layer-Specific Principles of Horizontal Axon Projection Patterns in Rat Vibrissal Cortex. Cereb Cortex 25, 4450–4468 (2015).

11. Fox, K., Wright, N., Wallace, H. & Glazewski, S. The Origin of Cortical Surround Receptive Fields Studied in the Barrel Cortex. J Neurosci 23, 8380–8391 (2003).

12. Clancy, K. B., Schnepel, P., Rao, A. T. & Feldman, D. E. Structure of a single whisker representation in layer 2 of mouse somatosensory cortex. J. Neurosci. 35, 3946–3958 (2015).

13. Wang, H. C., LeMessurier, A. M. & Feldman, D. E. Tuning instability of non-columnar neurons in the salt-and-pepper whisker map in somatosensory cortex. Nat Commun 13, 6611 (2022).

14. Fox, K. Anatomical pathways and molecular mechanisms for plasticity in the barrel cortex. Neuroscience 111, 799–814 (2002).

15. Polley, D. B., Chen-Bee, C. H. & Frostig, R. D. Two directions of plasticity in the sensory-deprived adult cortex. Neuron 24, 623–637 (1999).

16. Fox, K. A critical period for experience-dependent synaptic plasticity in rat barrel cortex. J Neurosci 12, 1826–1838 (1992).

17. Glazewski, S. & Fox, K. Time course of experience-dependent synaptic potentiation and depression in barrel cortex of adolescent rats. J. Neurophysiol. 75, 1714–1729 (1996).

18. Vasquez, B., Campos, B., Cao, A., Theint, A. T. & Zeiger, W. High-Sensitivity Intrinsic Optical Signal Imaging Through Flexible, Low-Cost Adaptations of an Upright Microscope. eNeuro 10, ENEURO.0046-23.2023 (2023).

19. Margolis, D. J. et al. Reorganization of cortical population activity imaged throughout long-term sensory deprivation. Nat. Neurosci. 15, 1539–1546 (2012).

20. Gainey, M. A. & Feldman, D. E. Multiple shared mechanisms for homeostatic plasticity in rodent somatosensory and visual cortex. Philos. Trans. R. Soc. Lond., B, Biol. Sci. 372, (2017).

21. Margolis, D. J., Lütcke, H. & Helmchen, F. Microcircuit dynamics of map plasticity in barrel cortex. Curr Opin Neurobiol 24, 76–81 (2014).

22. Xu, X., Roby, K. D. & Callaway, E. M. Immunochemical characterization of inhibitory mouse cortical neurons: three chemically distinct classes of inhibitory cells. J Comp Neurol 518, 389–404 (2010).

23. Gonchar, Y., Wang, Q. & Burkhalter, A. Multiple distinct subtypes of GABAergic neurons in mouse visual cortex identified by triple immunostaining. Front Neuroanat 1, 3 (2007).

24. Tamamaki, N. et al. Green fluorescent protein expression and colocalization with calretinin, parvalbumin, and somatostatin in the GAD67-GFP knock-in mouse. J Comp Neurol 467, 60–79 (2003).

25. Celio, M. R. Parvalbumin in most gamma-aminobutyric acid-containing neurons of the rat cerebral cortex. Science 231, 995–997 (1986).

26. Kawaguchi, Y., Katsumaru, H., Kosaka, T., Heizmann, C. W. & Hama, K. Fast spiking cells in rat hippocampus (CA1 region) contain the calcium-binding protein parvalbumin. Brain Res 416, 369–374 (1987).

27. Kawaguchi, Y. & Kubota, Y. Correlation of physiological subgroupings of nonpyramidal cells with parvalbumin- and calbindinD28k-immunoreactive neurons in layer V of rat frontal cortex. J Neurophysiol 70, 387–396 (1993).

28. Kawaguchi, Y. Physiological subgroups of nonpyramidal cells with specific morphological characteristics in layer II/III of rat frontal cortex. J Neurosci 15, 2638– 2655 (1995).

29. Katsumaru, H., Kosaka, T., Heizmann, C. W. & Hama, K. Immunocytochemical study of GABAergic neurons containing the calcium-binding protein parvalbumin in the rat hippocampus. Exp Brain Res 72, 347–362 (1988).

30. Czeiger, D. & White, E. L. Comparison of the distribution of parvalbumin-immunoreactive and other synapses onto the somata of callosal projection neurons in mouse visual and somatosensory cortex. J Comp Neurol 379, 198–210 (1997).

31. Di Cristo, G. et al. Subcellular domain-restricted GABAergic innervation in primary visual cortex in the absence of sensory and thalamic inputs. Nat Neurosci 7, 1184–1186 (2004).

32. Bruno, R. M. & Simons, D. J. Feedforward mechanisms of excitatory and inhibitory cortical receptive fields. J Neurosci 22, 10966–10975 (2002).

33. Swadlow, H. A. Influence of VPM afferents on putative inhibitory interneurons in S1 of the awake rabbit: evidence from cross-correlation, microstimulation, and latencies to peripheral sensory stimulation. J Neurophysiol 73, 1584–1599 (1995).

34. Mateo, C. et al. In vivo optogenetic stimulation of neocortical excitatory neurons drives brain-state-dependent inhibition. Curr Biol 21, 1593–1602 (2011).

35. DeNardo, L. A., Berns, D. S., DeLoach, K. & Luo, L. Connectivity of mouse somatosensory and prefrontal cortex examined with trans-synaptic tracing. Nat. Neurosci. 18, 1687–1697 (2015).

36. Avermann, M., Tomm, C., Mateo, C., Gerstner, W. & Petersen, C. C. H. Microcircuits of excitatory and inhibitory neurons in layer 2/3 of mouse barrel cortex. J Neurophysiol 107, 3116–3134 (2012).

37. McCasland, J. S., Hibbard, L. S., Rhoades, R. W. & Woolsey, T. A. Activation of a wide-spread network of inhibitory neurons in barrel cortex. Somatosens Mot Res 14, 138–147 (1997).

38. Swadlow, H. A. Efferent neurons and suspected interneurons in S-1 vibrissa cortex of the awake rabbit: receptive fields and axonal properties. J Neurophysiol 62, 288–308 (1989).

39. Yu, J., Hu, H., Agmon, A. & Svoboda, K. Recruitment of GABAergic Interneurons in the Barrel Cortex during Active Tactile Behavior. Neuron 104, 412–427.e4 (2019).

40. Yeganeh, F. et al. Effects of optogenetic inhibition of a small fraction of parvalbumin-positive interneurons on the representation of sensory stimuli in mouse barrel cortex. Sci Rep 12, 19419 (2022).

41. Yang, J.-W. et al. Optogenetic Modulation of a Minor Fraction of Parvalbumin-Positive Interneurons Specifically Affects Spatiotemporal Dynamics of Spontaneous and Sensory-Evoked Activity in Mouse Somatosensory Cortex in Vivo. Cereb. Cortex 27, 5784–5803 (2017).

42. Swadlow, H. A. Thalamocortical control of feed-forward inhibition in awake somatosensory ‘barrel’ cortex. Philos Trans R Soc Lond B Biol Sci 357, 1717–1727 (2002).

43. Khatri, V., Hartings, J. A. & Simons, D. J. Adaptation in thalamic barreloid and cortical barrel neurons to periodic whisker deflections varying in frequency and velocity. J Neurophysiol 92, 3244–3254 (2004).

44. Brumberg, J. C., Pinto, D. J. & Simons, D. J. Spatial gradients and inhibitory summation in the rat whisker barrel system. J Neurophysiol 76, 130–140 (1996).

45. Gabernet, L., Jadhav, S. P., Feldman, D. E., Carandini, M. & Scanziani, M. Somatosensory integration controlled by dynamic thalamocortical feed-forward inhibition. Neuron 48, 315–327 (2005).

46. Atallah, B. V., Bruns, W., Carandini, M. & Scanziani, M. Parvalbumin-expressing interneurons linearly transform cortical responses to visual stimuli. Neuron 73, 159– 170 (2012).

47. Cardin, J. A. et al. Driving fast-spiking cells induces gamma rhythm and controls sensory responses. Nature 459, 663–667 (2009).

48. Swadlow, H. A., Beloozerova, I. N. & Sirota, M. G. Sharp, local synchrony among putative feed-forward inhibitory interneurons of rabbit somatosensory cortex. J Neurophysiol 79, 567–582 (1998).

49. Gentet, L. J., Avermann, M., Matyas, F., Staiger, J. F. & Petersen, C. C. H. Membrane potential dynamics of GABAergic neurons in the barrel cortex of behaving mice. Neuron 65, 422–435 (2010).

50. Rupert, D. D. & Shea, S. D. Parvalbumin-Positive Interneurons Regulate Cortical Sensory Plasticity in Adulthood and Development Through Shared Mechanisms. Front Neural Circuits 16, 886629 (2022).

51. Reh, R. K. et al. Critical period regulation across multiple timescales. Proc Natl Acad Sci U S A 117, 23242–23251 (2020).

52. Takesian, A. E. & Hensch, T. K. Balancing plasticity/stability across brain development. Prog Brain Res 207, 3–34 (2013).

53. Yazaki-Sugiyama, Y., Kang, S., Câteau, H., Fukai, T. & Hensch, T. K. Bidirectional plasticity in fast-spiking GABA circuits by visual experience. Nature 462, 218–221 (2009).

54. Kuhlman, S. J. et al. A disinhibitory microcircuit initiates critical-period plasticity in the visual cortex. Nature 501, 543–546 (2013).

55. Kaplan, E. S. et al. Contrasting roles for parvalbumin-expressing inhibitory neurons in two forms of adult visual cortical plasticity. Elife 5, e11450 (2016).

56. Cisneros-Franco, J. M. & de Villers-Sidani, É. Reactivation of critical period plasticity in adult auditory cortex through chemogenetic silencing of parvalbumin-positive interneurons. Proc Natl Acad Sci U S A 116, 26329–26331 (2019).

57. Li, L., Gainey, M. A., Goldbeck, J. E. & Feldman, D. E. Rapid homeostasis by disinhibition during whisker map plasticity. Proc. Natl. Acad. Sci. U.S.A. 111, 1616–1621 (2014).

58. House, D. R. C., Elstrott, J., Koh, E., Chung, J. & Feldman, D. E. Parallel regulation of feedforward inhibition and excitation during whisker map plasticity. Neuron 72, 819– 831 (2011).

59. Zhang, Y. et al. Fast and sensitive GCaMP calcium indicators for imaging neural populations. Nature 10.1038/s41586-023-05828-9 (2023) doi:10.1038/s41586-023-05828-9.

60. Fenno, L. E. et al. Comprehensive Dual- and Triple-Feature Intersectional Single-Vector Delivery of Diverse Functional Payloads to Cells of Behaving Mammals. Neuron 107, 836–853.e11 (2020).

61. Krashes, M. J. et al. Rapid, reversible activation of AgRP neurons drives feeding behavior in mice. J Clin Invest 121, 1424–1428 (2011).

62. Hippenmeyer, S. et al. A developmental switch in the response of DRG neurons to ETS transcription factor signaling. PLoS Biol 3, e159 (2005).

63. Daigle, T. L. et al. A Suite of Transgenic Driver and Reporter Mouse Lines with Enhanced Brain-Cell-Type Targeting and Functionality. Cell 174, 465–480.e22 (2018).

64. Holtmaat, A. et al. Long-term, high-resolution imaging in the mouse neocortex through a chronic cranial window. Nat Protoc 4, 1128–1144 (2009).

65. Mostany, R. & Portera-Cailliau, C. A craniotomy surgery procedure for chronic brain imaging. J Vis Exp 10.3791/680 (2008) doi:10.3791/680.

66. Cantu, D. A. et al. EZcalcium: Open Source Toolbox for Analysis of Calcium Imaging Data. bioRxiv 2020.01.02.893198 (2020) doi:10.1101/2020.01.02.893198.

67. He, C. X. et al. Tactile Defensiveness and Impaired Adaptation of Neuronal Activity in the Fmr1 Knock-Out Mouse Model of Autism. J. Neurosci. 37, 6475–6487 (2017).

68. Pachitariu, M. et al. Suite2p: beyond 10,000 neurons with standard two-photon microscopy. 061507 Preprint at 10.1101/061507 (2017).

69. Thompson, K. J. et al. DREADD Agonist 21 Is an Effective Agonist for Muscarinic-Based DREADDs in Vitro and in Vivo. ACS Pharmacol. Transl. Sci. 10.1021/acsptsci.8b00012 (2018) doi:10.1021/acsptsci.8b00012.

70. Strata, P. & Harvey, R. Dale’s principle. Brain Res Bull 50, 349–350 (1999).

71. Miller, K. D. & Troyer, T. W. Neural noise can explain expansive, power-law nonlinearities in neural response functions. J Neurophysiol 87, 653–659 (2002).

72. Carandini, M. Amplification of trial-to-trial response variability by neurons in visual cortex. PLoS Biol 2, E264 (2004).

73. Jercog, D. et al. UP-DOWN cortical dynamics reflect state transitions in a bistable network. Elife 6, e22425 (2017).

74. Campagnola, L. et al. Local connectivity and synaptic dynamics in mouse and human neocortex. Science 375, eabj5861 (2022).

75. Tsodyks, M., Pawelzik, K. & Markram, H. Neural networks with dynamic synapses. Neural Comput 10, 821–835 (1998).

76. Devienne, G. et al. Regulation of Perineuronal Nets in the Adult Cortex by the Activity of the Cortical Network. J Neurosci 41, 5779–5790 (2021).

77. Stachniak, T. J., Ghosh, A. & Sternson, S. M. Chemogenetic synaptic silencing of neural circuits localizes a hypothalamus→midbrain pathway for feeding behavior. Neuron 82, 797–808 (2014).

78. Graf, J., Samiee, A., Flossmann, T., Holthoff, K. & Kirmse, K. Chemogenetic silencing reveals presynaptic Gi/o protein-mediated inhibition of developing hippocampal synchrony in vivo. iScience 27, 110997 (2024).

79. Chen, T.-W. et al. Ultrasensitive fluorescent proteins for imaging neuronal activity. Nature 499, 295–300 (2013).

80. Kingma, D. P. & Ba, J. Adam: A Method for Stochastic Optimization. Preprint at 10.48550/arXiv.1412.6980 (2017).

81. Zeiger, W. A. et al. Barrel cortex plasticity after photothrombotic stroke involves potentiating responses of pre-existing circuits but not functional remapping to new circuits. Nat Commun 12, 3972 (2021).

82. Yu, Z. et al. Beyond t test and ANOVA: applications of mixed-effects models for more rigorous statistical analysis in neuroscience research. Neuron 110, 21–35 (2022).

83. Benjamini, Y. & Hochberg, Y. Controlling the False Discovery Rate: A Practical and Powerful Approach to Multiple Testing. Journal of the Royal Statistical Society. Series B (Methodological) 57, 289–300 (1995).

84. Gainey, M. A., Aman, J. W. & Feldman, D. E. Rapid Disinhibition by Adjustment of PV Intrinsic Excitability during Whisker Map Plasticity in Mouse S1. J. Neurosci. 38, 4749– 4761 (2018).

85. Sanzeni, A. et al. Inhibition stabilization is a widespread property of cortical networks. Elife 9, e54875 (2020).

86. Rogers, S., Rozman, P. A., Valero, M., Doyle, W. K. & Buzsáki, G. Mechanisms and plasticity of chemogenically induced interneuronal suppression of principal cells. Proc Natl Acad Sci U S A 118, e2014157118 (2021).

87. Zhou, X., Mansori, I., Fischer, T., Witte, M. & Staiger, J. F. Characterizing the morphology of somatostatin-expressing interneurons and their synaptic innervation pattern in the barrel cortex of the GFP-expressing inhibitory neurons mouse. J Comp Neurol 528, 244– 260 (2020).

88. Wang, Y. et al. Anatomical, physiological and molecular properties of Martinotti cells in the somatosensory cortex of the juvenile rat. J Physiol 561, 65–90 (2004).

89. Yavorska, I. & Wehr, M. Somatostatin-Expressing Inhibitory Interneurons in Cortical Circuits. Front Neural Circuits 10, 76 (2016).

90. Preuss, F., Walker, F., Möck, M., Witte, M. & Staiger, J. F. Inhibitory circuit motifs of cortical somatosensory layer 5 SST interneurons are uniform within layers but specific across layers. J Physiol 604, 479–502 (2026).

91. Walker, F. et al. Parvalbumin- and vasoactive intestinal polypeptide-expressing neocortical interneurons impose differential inhibition on Martinotti cells. Nat Commun 7, 13664 (2016).

92. Pfeffer, C. K., Xue, M., He, M., Huang, Z. J. & Scanziani, M. Inhibition of inhibition in visual cortex: the logic of connections between molecularly distinct interneurons. Nat Neurosci 16, 1068–1076 (2013).

93. Bertelsen, N. et al. Electrophysiologically-defined excitation-inhibition autism neurosubtypes. 2023.11.22.23298729 Preprint at 10.1101/2023.11.22.23298729 (2026).

94. Moore, A. K., Weible, A. P., Balmer, T. S., Trussell, L. O. & Wehr, M. Rapid Rebalancing of Excitation and Inhibition by Cortical Circuitry. Neuron 97, 1341–1355.e6 (2018).

95. Billeh, Y. N. et al. Systematic Integration of Structural and Functional Data into Multi-scale Models of Mouse Primary Visual Cortex. Neuron 106, 388–403.e18 (2020).

96. Tsodyks, M. V., Skaggs, W. E., Sejnowski, T. J. & McNaughton, B. L. Paradoxical effects of external modulation of inhibitory interneurons. J Neurosci 17, 4382–4388 (1997).

97. Hafner, G. et al. Mapping Brain-Wide Afferent Inputs of Parvalbumin-Expressing GABAergic Neurons in Barrel Cortex Reveals Local and Long-Range Circuit Motifs. Cell Rep 28, 3450–3461.e8 (2019).

98. Kerlin, A. M., Andermann, M. L., Berezovskii, V. K. & Reid, R. C. Broadly tuned response properties of diverse inhibitory neuron subtypes in mouse visual cortex. Neuron 67, 858–871 (2010).

99. Liu, B. et al. Visual receptive field structure of cortical inhibitory neurons revealed by two-photon imaging guided recording. J Neurosci 29, 10520–10532 (2009).

100. Niell, C. M. & Stryker, M. P. Highly selective receptive fields in mouse visual cortex. J Neurosci 28, 7520–7536 (2008).

101. Guy, J., Möck, M. & Staiger, J. F. Direction selectivity of inhibitory interneurons in mouse barrel cortex differs between interneuron subtypes. Cell Rep 42, 111936 (2023).

102. Swadlow, H. A. & Gusev, A. G. Receptive-field construction in cortical inhibitory interneurons. Nat Neurosci 5, 403–404 (2002).

103. Dobler, Z. et al. Adapting and facilitating responses in mouse somatosensory cortex are dynamic and shaped by experience. Curr Biol 34, 3506–3521.e5 (2024).

104. Finnerty, G. T., Roberts, L. S. & Connors, B. W. Sensory experience modifies the short-term dynamics of neocortical synapses. Nature 400, 367–371 (1999).

105. Marik, S. A., Yamahachi, H., McManus, J. N. J., Szabo, G. & Gilbert, C. D. Axonal dynamics of excitatory and inhibitory neurons in somatosensory cortex. PLoS Biol 8, e1000395 (2010).

106. Albieri, G. et al. Rapid Bidirectional Reorganization of Cortical Microcircuits. Cereb Cortex 25, 3025–3035 (2015).

107. Trakoshis, S. et al. Intrinsic excitation-inhibition imbalance affects medial prefrontal cortex differently in autistic men versus women. Elife 9, e55684 (2020).

108. Pérez-Montoyo, E. et al. Inhibitory tone in the dentate gyrus dynamically prioritizes memory flexibility or stability by tuning a sensitivity-consistency continuum. PLoS Biol 24, e3003956 (2026).

109. Fu, X. et al. Gq neuromodulation of BLA parvalbumin interneurons induces burst firing and mediates fear-associated network and behavioral state transition in mice. Nat Commun 13, 1290 (2022).

110. Yi, F. et al. Direct excitation of parvalbumin-positive interneurons by M1 muscarinic acetylcholine receptors: roles in cellular excitability, inhibitory transmission and cognition. J Physiol 592, 3463–3494 (2014).

111. Huang, L. et al. Relationship between simultaneously recorded spiking activity and fluorescence signal in GCaMP6 transgenic mice. Elife 10, e51675 (2021).

112. Rupprecht, P., Rózsa, M., Fang, X., Svoboda, K. & Helmchen, F. Spike inference from calcium imaging data acquired with GCaMP8 indicators. Nat Methods 10.1038/s41592-026-03183-x (2026) doi:10.1038/s41592-026-03183-x.

113. Goel, A. et al. Impaired perceptual learning in a mouse model of Fragile X syndrome is mediated by parvalbumin neuron dysfunction and is reversible. Nat. Neurosci. 21, 1404–1411 (2018).

114. Kourdougli, N. et al. Improvement of sensory deficits in fragile X mice by increasing cortical interneuron activity after the critical period. Neuron 111, 2863–2880.e6 (2023).

115. Znamenskiy, P. et al. Functional specificity of recurrent inhibition in visual cortex. Neuron 112, 991–1000.e8 (2024).

116. Garcia-Junco-Clemente, P., Tring, E., Ringach, D. L. & Trachtenberg, J. T. State-Dependent Subnetworks of Parvalbumin-Expressing Interneurons in Neocortex. Cell Rep 26, 2282–2288.e3 (2019).

117. Khan, A. G. et al. Distinct learning-induced changes in stimulus selectivity and interactions of GABAergic interneuron classes in visual cortex. Nat Neurosci 21, 851– 859 (2018).

118. Bernardo, K. L., McCasland, J. S., Woolsey, T. A. & Strominger, R. N. Local intra- and interlaminar connections in mouse barrel cortex. J Comp Neurol 291, 231–255 (1990).

119. Hoffer, Z. S., Hoover, J. E. & Alloway, K. D. Sensorimotor corticocortical projections from rat barrel cortex have an anisotropic organization that facilitates integration of inputs from whiskers in the same row. J Comp Neurol 466, 525–544 (2003).

120. Lee, S.-H., Land, P. W. & Simons, D. J. Layer- and cell-type-specific effects of neonatal whisker-trimming in adult rat barrel cortex. J Neurophysiol 97, 4380–4385 (2007).

121. Simons, D. J., Carvell, G. E. & Kyriazi, H. T. Alterations in functional thalamocortical connectivity following neonatal whisker trimming with adult regrowth. J Neurophysiol 114, 1912–1922 (2015).

122. Kalambogias, J. et al. Development and sensory experience dependent regulation of microglia in barrel cortex. J Comp Neurol 528, 559–573 (2020).

123. Barrera, K. et al. Organization of myelin in the mouse somatosensory barrel cortex and the effects of sensory deprivation. Dev Neurobiol 73, 297–314 (2013).

124. Murokh, S. et al. X-ray diffraction reveals alterations in mouse somatosensory cortex following sensory deprivation. J Neurosci Methods 424, 110582 (2025).

125. McRae, P. A., Rocco, M. M., Kelly, G., Brumberg, J. C. & Matthews, R. T. Sensory deprivation alters aggrecan and perineuronal net expression in the mouse barrel cortex. J Neurosci 27, 5405–5413 (2007).

126. Nowicka, D., Soulsby, S., Skangiel-Kramska, J. & Glazewski, S. Parvalbumin-containing neurons, perineuronal nets and experience-dependent plasticity in murine barrel cortex. Eur J Neurosci 30, 2053–2063 (2009).

127. Chen, C.-C., Bajnath, A. & Brumberg, J. C. The impact of development and sensory deprivation on dendritic protrusions in the mouse barrel cortex. Cereb Cortex 25, 1638– 1653 (2015).

128. Miquelajauregui, A. et al. Layer 4 Pyramidal Neurons Exhibit Robust Dendritic Spine Plasticity In Vivo after Input Deprivation. J. Neurosci. 35, 7287–7294 (2015).

129. Dobrzanski, G., Zakrzewska, R., Kossut, M. & Liguz-Lecznar, M. Impact of somatostatin interneurons on interactions between barrels in plasticity induced by whisker deprivation. Sci Rep 12, 17992 (2022).

