## Supplementary Information for "Dynamic recruitment of PV interneurons in the somatosensory cortex induced by experience-dependent plasticity"

Throughout the Supplementary Information, Gq and Gi denote the hM3D(Gq) and hM4D(Gi) conditions, respectively.

### Supplementary Methods

#### Model parameters

The fitted base matrix is given below, with target populations in rows and source populations in columns, ordered Pyr, PV, SST and VIP. Italics mark magnitudes below 0.15, the threshold below which connections are omitted from the schematic in Fig. 6a; this marking is for display only. All connections were retained. Gi scaled every PV output, including PV→SST.

$$W = \begin{pmatrix} 3.36243 & -2.76387 & -1.72472 & -0.86920 \\ 2.40461 & -1.04056 & -0.19945 & \mathit{-0.11977} \\ 2.12219 & \mathit{-0.03426} & \mathit{-0.00863} & -1.67819 \\ 2.05910 & -1.80294 & -1.19315 & \mathit{-0.00109} \end{pmatrix}.$$

Bounds were enforced by a logistic reparameterization, $\theta_k = l_k + (u_k - l_k)\mathrm{expit}(v_k)$, with limits in the released search configuration. SST→SST and VIP→VIP were restricted a priori to $[-0.03, -0.001]$ and $[-0.002, -10^{-6}]$, respectively. We chose these narrow bounds to reflect the sparse recurrent connectivity reported within SST and VIP populations[1–3]. VIP→PV and PV→SST were also bounded a priori, within [−0.696, −0.001] and [−0.800, −0.001], respectively. Although these broader bounds did not require weak coupling, optimization converged on small magnitudes for both connections (−0.120 and −0.034, respectively), consistent with the weak VIP→PV and PV→SST inhibition measured experimentally[2].

The 49 free parameters comprised 16 weights, 4 background inputs, 3 sensory amplitudes, 4 rate time constants, 4 Control softness values, 7 adaptation and short-term plasticity parameters, 2 tonic PV inputs, 8 softness ratios and 1 Gi PV efficacy; fitted values are given in the matrix *W* above and in Supplementary Tables S1–S4. VIP sensory input was fixed at zero. Calcium time constants and population observation gains were fixed at the values in Supplementary Table S1. Control tonic PV input, PV output efficacy and softness ratios were fixed at 0, 1 and 1, respectively.

| Parameter | Pyr | PV | SST | VIP |
|---|---|---|---|---|
| Background input $b_i$ (Hz) | −0.17800 | 0.62139 | 0.01457 | 1.02261 |
| Sensory input $s_i$ (Hz) | 1.74712 | 0.10993 | 0.07622 | 0 (fixed) |
| Rate time constant $\tau_i$ (s) | 0.01137 | 0.01897 | 0.04690 | 0.03381 |
| Control softness $\sigma_i^{(\mathrm{Control})}$ (Hz) | 0.75274 | 0.97828 | 0.56930 | 0.62746 |
| Calcium time constant $\tau_{C,i}$ (s; fixed) | 0.60 | 0.60 | 0.35 | 0.60 |
| Observation gain $\gamma_i$ (fixed) | 1.46145 | 0.75757 | 0.08120 | 1 |

**Supplementary Table S1. Shared population parameters.** Except for Control softness, which the softness ratios in Supplementary Table S3 rescale under Gi and Gq, these parameters were shared across conditions. Parameters marked fixed were not optimized.

| Process | Parameter | Fitted value |
|---|---|---|
| Pyr adaptation | $g_A$ | 0.85082 |
| Pyr adaptation | $\tau_A$ (s) | 0.50203 |
| Pyr→PV depression | $U_D$ | 0.62151 |
| Pyr→PV depression | $\tau_D$ (s) | 0.91216 |
| Pyr→SST facilitation | $U_F$ | 1.11703 |
| Pyr→SST facilitation | $\tau_F$ (s) | 0.50840 |
| Pyr→SST facilitation | $f_{\max}$ | 3.12395 |

**Supplementary Table S2. Adaptation and short-term plasticity parameters.** Symbols follow Eqs. 3 and 4 of the main Methods. All values were fitted and shared across conditions.

| Parameter | Control (fixed) | Gi | Gq |
|---|---|---|---|
| Tonic PV input $d_{\mathrm{PV}}^{(c)}$ (Hz) | 0 | −0.08028 | +0.28067 |
| PV output efficacy $\kappa_{\mathrm{PV}}^{(c)}$ | 1 | 0.38285 | 1 (fixed) |
| Pyr softness ratio $\rho_{\mathrm{E}}^{(c)}$ | 1 | 1.47640 | 0.41277 |
| PV softness ratio $\rho_{\mathrm{PV}}^{(c)}$ | 1 | 1.99484 | 0.44163 |
| SST softness ratio $\rho_{\mathrm{SST}}^{(c)}$ | 1 | 3.33160 | 0.81081 |
| VIP softness ratio $\rho_{\mathrm{VIP}}^{(c)}$ | 1 | 2.89115 | 0.54452 |

**Supplementary Table S3. Condition-dependent parameters.** Gi and Gq values are the fitted 5-mg/kg endpoints. Softness ratios multiply the Control softness in Supplementary Table S1, and intermediate doses follow the dose paths in Supplementary Table S4. Parameters marked fixed were not optimized.

| Condition | Prescribed sigmoid argument at doses 0 and 5 | Fitted span $\zeta_c$ | Derived $h_c(1)$ | Derived $h_c(3)$ |
|---|---|---|---|---|
| Gq | $0, +\zeta_{\mathrm{Gq}}$ | 6.44561 | 0.5698 | 0.9621 |
| Gi | $-\zeta_{\mathrm{Gi}}, +9\zeta_{\mathrm{Gi}}$ | 2.99287 | 0.9499 | 1.0000 |

**Supplementary Table S4. Dose-path parameters.** The span $\zeta_c$ was the only parameter fitted during dose calibration (Eq. S4). The derived progress $h_c(d)$ equals 0 in Control and 1 at the fitted 5-mg/kg endpoint (main Methods, Eqs. 7 and 8).

### Parameter search

We fitted the nine population-mean calcium traces jointly using the bounded circuit parameters and fixed observation model described in the main Methods. At search stage *s*, the objective combined full (0–3 s after stimulus onset) and early (0–1 s after stimulus onset) trace errors from Eq. 6 with additional terms:

$$L_s = \lambda_{s,full} E_{[0,3]} + \lambda_{s,early} E_{[0,1]} + R_s \quad \text{(S1)}$$

The full-trace and early-trace weights in Eq. S1 were (1, 1.5), (4, 6) and (8, 14) in stages 1–3, respectively. The additional term $R_s$ included Gi peak-amplitude error and penalties encouraging consistent prestimulus and stimulus-period equilibria, stable dynamics, limited firing rates and oscillations, and, in Control and Gq, an inhibition-stabilized stimulus-period state in which added tonic PV input lowers Pyr, PV and SST rates. We additionally imposed a low-or-close baseline prior on Control and Gq, allowing either low rates in both conditions or a small observation-scaled difference.

The three stages used 192, 144 and 80 starts, with 1,800, 1,800 and 1,600 iterations per start, respectively. All first-stage starts were random. Stage 2 used 120 perturbed starts from 12 retained candidates plus 24 fresh random starts; stage 3 used 64 perturbed starts from 8 retained candidates plus 16 fresh random starts. The final stage weighted the Gi waveform more heavily, including its early peak, subsequent decay, and lack of a late rate increase. Auxiliary equilibrium variables aided training; equilibria were reconstructed independently for evaluation. Circuit penalties guided the search toward Gq suppression, but the selected model was not required to satisfy the inhibition-stabilization and PV-input criteria.

For reporting, we averaged normalized root-mean-square errors across the nine traces:

$$\overline{\mathrm{NRMSE}}_T = \frac{1}{9}\sum_c \sum_i \frac{\sqrt{N_T^{-1}\sum_{t\in T}[\hat{C}_{ci}(t)-Y_{ci}(t)]^2}}{\max[D_T(Y_{ci}),0.05]}. \quad \text{(S2)}$$

Here, $D_T(Y_{ci})$ is the target range within the scoring window, with a floor of 0.05; training instead used the full 0–3-s target range for all windows. Mean normalized errors were 0.297 over 0–1 s and 0.221 over 0–3 s. The accompanying code specifies the complete objective, parameter bounds, search settings, and candidate evaluations.

#### Dose calibration

Experimental 0–1-s AUCs were calculated after subtracting each trial's prestimulus mean. The common-mouse cohort comprised four Gi and six Gq mice measured at doses 1, 3 and 5. For dose 0, we first averaged each mouse's available baseline sessions. At each dose, mouse-level AUCs were averaged with equal weight across mice. Pyr and PV also contributed equally to dose calibration. SST supplied only endpoint measurements.

To reduce the risk of overfitting the limited dose measurements, we constrained all condition-dependent parameters to a common sigmoidal progression within each condition (Eqs. 7–11). With the Control and dose-5 endpoints fixed, only one shared span was fitted, rather than a separate sigmoid for each parameter. Dose calibration proceeded in two steps. First, we fitted a smooth empirical AUC curve to the mouse-averaged measurements:

$$A_i(d) = A_{i,\mathrm{offset}} + A_{i,\mathrm{span}}\mathrm{expit}[k(d-m)], \quad \text{(S3)}$$

with population-specific offsets and amplitudes and a shared slope and midpoint. The unweighted squared error used eight population-by-dose means. At each $k, m$, population coefficients were solved by least squares, constraining the dose-0-to-5 change negative for Gq and positive for Gi. Differential evolution followed by bounded local minimization optimized $k, m$.

Second, we fitted the span of the model dose path (Eqs. 7–11) by minimizing the mean squared error $E_{\mathrm{dose}}$ in Eq. S4. This compared AUCs from circuit simulations with the smooth empirical curve after normalizing each to its own endpoints:

$$A_i^*(d) = \frac{A_i(d)-A_i(0)}{A_i(5)-A_i(0)},$$
$$E_{\mathrm{dose}} = \frac{1}{2N_D}\sum_{i\in\{\mathrm{E,PV}\}}\sum_{d\in D}[A_i^{*,\mathrm{model}}(d) - A_i^{*,\mathrm{data}}(d)]^2. \quad \text{(S4)}$$

Here $D$ is the interior dose grid, $0 < d < 5$, sampled in 0.1-mg/kg steps and containing $N_D$ = 49 points. Endpoint residuals vanish by construction. Only the dose-path span $\zeta_c$ in Eqs. 7 and 8 was optimized to minimize the discrepancy in Eq. S4, with circuit parameters held fixed. Final dose curves were checked at 1-ms resolution for stable, bounded dynamics.

For the Gq extrapolation shown in Supplementary Fig. S15e, C and Q denote Control and Gq. A separate coordinate $\eta$ ran from 0 to 3.5 in steps of 0.025. Tonic input was $d_{\mathrm{PV}}(\eta) = \eta d_{\mathrm{PV}}^{(Q)}$, with linear softness $\sigma_{i,\mathrm{lin}}(\eta) = \sigma_i^{(C)}[1+\eta(\rho_i^{(Q)}-1)]$ and a smooth positive floor:

$$\sigma_i(\eta) = \sigma_{i,\min} + \delta_i \log\left[1 + \exp\frac{\sigma_{i,\mathrm{lin}}(\eta)-\sigma_{i,\min}}{\delta_i}\right]. \quad \text{(S5)}$$

We used $\sigma_{i,\min} = 0.40\sigma_i^{(Q)}$ and $\delta_i = 0.05\sigma_i^{(C)}$. PV efficacy and all other parameters remained fixed. This floor only approximates the fitted softness at $\eta = 0{,}1$. Stimulus-period fixed-point rates were normalized to $\eta = 0$. The region $\eta > 1$ was not calibrated to C21 doses.

#### Expansion into a finite heterogeneous network

We expanded the four mean-field populations into corresponding Pyr, PV, SST and VIP populations containing 1,400 rate units in total, with sizes listed in Supplementary Table S5. This retained the mean-field circuit organization while allowing individual neurons within each population to differ. The mean-field parameters were not refitted, so the network tested whether the fitted population dynamics persisted in a finite, heterogeneous circuit. The units formed a Bernoulli graph without autapses. Nonzero connections were normalized by expected presynaptic indegree, recovering homogeneous mean-field input in expectation. The graph and static cell properties were shared across conditions. For neuron $n$ in population $i$,

$$\tau_i \frac{dr_{in}}{dt} = -r_{in} + \phi_{\sigma_{i,\mathrm{tr}}^{(c)}} \Big( \sum_j \sum_{m\in j} J_{in,jm}^{(c)} Q_{in,jm}(t) r_{jm} + b_i + s_{in}(t) + \mathbb{1}_{i=\mathrm{PV}} d_{\mathrm{PV}}^{(c)} - \mathbb{1}_{i=E} a_{En}(t) + \xi_{in}(t) \Big) \quad \text{(S6)}$$

Each Pyr neuron followed Eqs. 3 and 4 using its own firing rate. The factor $Q$ in Eq. S6 describes synaptic short-term plasticity. It equals the presynaptic Pyr neuron's depression state on Pyr→PV projections and its facilitation state on Pyr→SST projections; on projections without short-term plasticity, it equals one. Each presynaptic Pyr neuron shared its plasticity states across the corresponding terminals. Independent zero-mean Ornstein–Uhlenbeck (OU) fluctuations $\xi_{in}$ resolved part of the fitted softness:

$$\sigma_{i,\mathrm{OU}}^{(c)} = q\sigma_i^{(c)}, \sigma_{i,\mathrm{tr}}^{(c)} = \sqrt{1-q^2}\,\sigma_i^{(c)}, q = 0.5. \quad \text{(S7)}$$

A fraction $q^2 = 0.25$ of the input variance was therefore explicit; the remaining 75% was represented by the residual transfer-function softness in Eq. S6. The partition exactly recovers the transfer function for static Gaussian input. Equivalence in the recurrent dynamics is approximate and was tested by simulation. We selected $q = 0.5$ by qualitative comparison with the recorded single-cell responses, without fitting it to individual traces. To initialize trials in each condition, we first computed the network's steady state in the absence of sensory input. This deterministic calculation used the full fitted transfer-function softness $\sigma_i^{(\mathrm{c})}$ and no explicit OU noise to initialize rates, adaptation and plasticity states. We then simulated 10 s of prestimulus activity followed by the stimulus and poststimulus periods, using residual softness and explicit OU input throughout.

In exploratory simulations, sparse connectivity and independent noise left single-neuron responses more uniform than those observed experimentally. We therefore explored additional sources of heterogeneity in input strength and time course, as well as in calcium/fluorescence readout, to examine whether the network could better capture this diversity while retaining the fitted population dynamics. These additions were phenomenological and were not estimated by the mean-field fit.

We first varied how strongly each neuron received sensory drive and recurrent excitation from Pyr neurons, representing differences in coupling to the stimulus-recruited ensemble. In Eq. S6, sensory gains $g_{in}$ multiply the population input amplitude $s_i$ within $s_{in}(t)$ (Eq. S9). Recurrent Pyr-input gains $h_{in}$ multiply all incoming Pyr connections to each target neuron, so $J_{in,Em}^{(\mathrm{c})} = h_{in}\bar{J}_{in,Em}^{(\mathrm{c})}$, where $\bar{J}_{in,Em}^{(\mathrm{c})}$ denotes the sparse weights before this gain is applied. Initial sensory gains were lognormal and normalized within each population. Recurrent Pyr-input gains were generated from normalized beta draws with coefficient of variation (CV):

$$B_n \sim \mathrm{Beta}(\alpha, 1), \alpha = \sqrt{1+\mathrm{CV}^{-2}} - 1, u_n = B_n/\bar{B}. \quad \text{(S8)}$$

For a gain type with zero CV, all gains equalled one, leaving that component homogeneous. CVs in Supplementary Table S5 describe underlying draws, before recruitment and balancing. To generate a continuous tail of weakly recruited SST cells, we assigned each SST neuron n a fixed, positive recruitment factor $R_n$, drawn from the normalized beta distribution in Eq. S8 with CV 0.80. The same factor scaled both input pathways of that neuron: it multiplied the initial sensory gain by $R_n$ and the initial recurrent Pyr-input gain by $R_n^{0.25}$. Thus, $R_n$ is absorbed into the final gains $g_{\mathrm{SST},n}$ and $h_{\mathrm{SST},n}$, which enter Eq. S6 through sensory input (Eq. S9) and recurrent Pyr coupling $J^{(\mathrm{c})}_{\mathrm{SST}n,Em}$, respectively. The smaller exponent made recruitment heterogeneity weaker for recurrent than for sensory input; it was chosen phenomenologically rather than inferred from single-cell measurements. Recurrent gains were renormalized to mean one.

Neurons in sensory cortex can respond at different times during stimulation. To represent diversity in sensory recruitment, we relaxed the mean-field assumption of a common rectangular input profile within each population and allowed sensory input to vary in both amplitude and time course. During the 1-s stimulus,

$$s_{in}(t) = s_i g_{in}[1 + \mu_i z_{in}(2e^{-t/\tau_s} - 1)], \quad \text{(S9)}$$

with $t = 0$ at onset and coordinates $z_{in}$ evenly spaced on $[-1,1]$. The signed temporal modulation produces declining input for positive coordinates and rising input for negative coordinates; the total sensory input remains non-negative. Paired SST units at opposite sensory coordinates shared $R_n$, and initial sensory gains were orthogonalized against that coordinate. Final sensory gains satisfied $\langle g \rangle_i = 1$ and $\langle gz \rangle_i = 0$, so the population mean remained the same rectangular input as in the mean-field model at every time. These operations preserve gain means, but not necessarily response means. Input was zero outside the stimulus.

Similar firing activity can also yield different fluorescence amplitudes across neurons because the mapping from spiking to calcium and fluorescence varies between cells. We therefore introduced cell-specific readout gains to capture this source of variability. For each condition and trial (indices omitted), we first applied the same calcium filter as in the mean-field model (Eq. 5) to each neuron's absolute firing rate:

$$\tau_{\mathrm{C},i} \frac{dC_{in}}{dt} = r_{in} - C_{in}. \quad \text{(S10)}$$

We then applied heterogeneous SST calcium gains and additional Pyr and PV observation gains, using the normalized beta distribution in Eq. S8. Because these gains were constant in time, their effect can be written as multiplicative scaling after this linear filter. The observed and baseline-subtracted signals were

$$\begin{aligned} C^{\mathrm{obs}}_{in}(t) &= \gamma_i g^{\mathrm{Ca}}_{in} g^{\mathrm{obs}}_{in} C_{in}(t) + \varepsilon_{in}(t), \\ \Delta C^{\mathrm{obs}}_{in}(t) &= C^{\mathrm{obs}}_{in}(t) - \langle C^{\mathrm{obs}}_{in} \rangle_{\mathrm{pre}}. \end{aligned} \quad \text{(S11)}$$

Here $g^{\mathrm{Ca}}_{in}$ is the heterogeneous SST calcium gain and equals one in the other populations; $g^{\mathrm{obs}}_{in}$ is the additional heterogeneous Pyr or PV observation gain and equals one in SST and VIP. SST calcium gains were drawn independently of the recruitment factor $R_n$. The population gain $\gamma_i$ and calcium time constant were inherited from the mean-field model. The additive term $\varepsilon_{in}(t)$ is zero-mean, temporally correlated Gaussian readout noise. It represents variability in the fluorescence measurement and affects the observed signal without entering the rate dynamics in Eq. S6. Gains and noise amplitudes were shared across conditions, with independent noise realizations.

The prestimulus mean in Eq. S11 was calculated separately for each neuron and trial over the simulated 10 s before stimulus onset. These baseline-subtracted traces were then averaged across trials. Simulations used Euler integration with the settings in Supplementary Table S6.

| Setting | Pyr | PV | SST | VIP |
|---|---|---|---|---|
| Number of units | 800 | 200 | 300 | 100 |
| Sensory-gain CV | 0.25 | 0.80 | 0.80 | 0.25 |
| Recurrent Pyr-input gain CV | 0 | 0.15 | 0.15 | 0 |
| Recruitment-factor CV | 0 | 0 | 0.80 | 0 |
| Calcium-gain CV | 0 | 0 | 0.80 | 0 |
| Additional observation-gain CV | 0.75 | 0.50 | 0 | 0 |
| Readout-noise SD (model-calcium units) | 0.090 | 0.040 | 0.025 | 0.040 |
| Temporal loading $\mu_i$ | 0.4 | 0.4 | 0.4 | 0 |

**Supplementary Table S5. Population-specific settings of the heterogeneous network.** Settings were fixed after exploratory comparisons and were not fitted. CV, coefficient of variation.

| Setting | Value |
|---|---|
| Connection probability | 0.25 |
| OU amplitude fraction $q$ | 0.50 |
| OU correlation time | 30 ms |
| SST recruitment exponent | 0.25 |
| Sensory time constant $\tau_s$ | 50 ms |
| Readout-noise correlation time | 50 ms |
| Integration step | 1 ms |
| Saved frame rate | 10 Hz |
| Prestimulus / stimulus / poststimulus | 10 / 1 / 1 s |
| Trials per condition | 20 |

**Supplementary Table S6. Shared settings of the heterogeneous network.** OU, Ornstein–Uhlenbeck.

### Supplementary Results

#### Tonic PV drive and transfer softness shape Gq responses

We modeled Gq as increased tonic input to PV neurons combined with reduced transfer softness in all four populations (Supplementary Table S3). Because calcium responses are measured relative to the prestimulus baseline, a smaller PV response could reflect either lower PV firing or a baseline that rises more than the stimulus-evoked activity. To distinguish these possibilities and to separate the contributions of the two parameter changes, we compared three model states: Control; tonic only, which combines the fitted Gq tonic PV input with Control softness; and fitted Gq. Three paths connect these states: (i) from Control to tonic only, changing tonic input alone; (ii) from Control to fitted Gq, changing tonic input and softness together; and (iii) from tonic only to fitted Gq, changing softness alone at the fitted Gq tonic input. Along each path, the changing parameters varied linearly and were sampled at five evenly spaced points, including both endpoints; these points are not calibrated to C21 doses. All other model and observation parameters were unchanged. For every parameter setting, we recomputed the prestimulus equilibrium before applying the stimulus, so that each response was measured relative to its own baseline. Along all three paths, early baseline-subtracted calcium responses (0–200-ms AUC) decreased monotonically in Pyr, PV and SST, consistent with the experimentally observed Gq suppression.

Supplementary Table S7 lists the values at the three model states. Along path (i), tonic input alone raised PV baseline and stimulus-period fixed-point firing rates but lowered the early PV firing-rate peak. The PV calcium increment at 200 ms fell by 35% relative to Control, even though absolute PV calcium at that time rose by 16%, because the baseline rose more than the stimulus-evoked level. Along path (iii), which ends at the Gq condition of the model presented in the main text (Fig. 6b), reducing softness at the fitted Gq tonic input lowered PV baseline and stimulus-period fixed-point rates and further reduced the calcium increment, to 45% below Control. Thus, tonic drive alone was sufficient to reduce the early PV calcium increment, whereas the return of PV baseline firing to near its Control level and the lower stimulus-period PV firing of fitted Gq arose only when softness was also reduced. A smaller calcium increment alone does not establish classical paradoxical suppression of absolute PV firing[4,5].

| Measure | Control | Tonic only | Fitted Gq |
|---|---|---|---|
| PV baseline firing rate (Hz) | 0.4479 | 0.5506 | 0.4435 |
| PV stimulus-period fixed-point firing rate (Hz) | 0.6003 | 0.6642 | 0.5376 |
| PV peak firing rate, 0–200 ms (Hz) | 0.9312 | 0.7605 | 0.6605 |
| PV absolute calcium at 200 ms (a.u.) | 0.38606 | 0.44733 | 0.36173 |
| PV calcium increment at 200 ms (a.u.) | 0.04672 | 0.03018 | 0.02575 |
| PV calcium increment AUC, 0–200 ms (a.u.·s) | 0.005374 | 0.003251 | 0.002892 |
| Pyr calcium increment AUC, 0–200 ms (a.u.·s) | 0.010785 | 0.005646 | 0.003668 |
| SST calcium increment AUC, 0–200 ms (a.u.·s) | 0.000598 | 0.000388 | 0.000276 |

**Supplementary Table S7. PV firing rates and calcium responses under isolated and combined Gq parameter changes.** Values are from deterministic mean-field simulations with a 1-ms step. Tonic only combines the fitted Gq tonic PV input with Control softness, and fitted Gq uses the fitted Gq values of both. The three columns are therefore the endpoints of the paths described in the text: path (i) runs from Control to tonic only, path (ii) from Control to fitted Gq, and path (iii) from tonic only to fitted Gq. Times are relative to stimulus onset, and the stimulus remains on throughout 0–200 ms. Absolute calcium includes the fixed PV observation gain $\gamma_{PV}$ (Supplementary Table S1). Because model calcium equals the firing rate at the prestimulus equilibrium, absolute calcium equals the calcium increment plus $\gamma_{PV}$ times the PV baseline firing rate. All calcium increments, including the Pyr and SST AUCs, include the corresponding population's fixed observation gain. The stimulus-period fixed-point rate was solved separately. Calcium values at 200 ms and AUC over 0–200 ms are distinct from the 0–1-s peak and AUC measures used elsewhere.

#### Complete PV silencing produces disinhibition rather than Gq-associated suppression

To test whether complete loss of PV firing could account for Gq-associated suppression, we clamped PV firing to zero throughout the prestimulus and stimulus periods, eliminating all recurrent PV output. We compared the unmodified and clamped models under Control and fitted dose-5 Gq. All other model and observation parameters retained their condition-specific values, and each simulation started from its own equilibrium baseline. No parameters were refitted.

PV silencing produced large stimulus-onset transients in Pyr, SST and VIP, followed by a decline in activity (Supplementary Table S8). Under Gq, these populations crossed the prespecified 40-Hz validity bound within 28–37 ms of onset. The same qualitative failure occurred under Control. The clamped prestimulus equilibria remained stable, but the only stimulus fixed point located in each condition by the bounded multistart search was unstable. Reducing the integration step from 1 ms to 0.5 or 0.25 ms, or increasing it to 2 ms, preserved the rate-bound violations in all three populations.

Thus, within the fitted circuit, complete PV firing loss produces a disinhibited onset response rather than the observed Gq-associated suppression. Rates beyond the model's validity bound indicate departure from its intended operating regime and are not physiological predictions.

| Quantity | Control | Control + PV clamp | Gq | Gq + PV clamp |
|---|---|---|---|---|
| Peak Pyr rate (Hz) | 0.591 | 312 | 0.195 | 321 |
| Peak PV rate (Hz) | 0.931 | 0 | 0.660 | 0 |
| Peak SST rate (Hz) | 0.281 | 578 | 0.187 | 595 |
| Peak VIP rate (Hz) | 0.471 | 112 | 0.256 | 115 |
| First crossing of 40 Hz, Pyr / SST / VIP (ms) | — / — / — | 28 / 35 / 37 | — / — / — | 28 / 35 / 37 |
| Baseline $\lambda_{max}$ $(s^{-1})$ | −1.098 | −1.113 | −1.096 | −1.096 |
| Stimulus $\lambda_{max}$ $(s^{-1})$ | −1.150 | +117.947 | −1.111 | +118.853 |

**Supplementary Table S8. Mean-field model response to complete PV firing loss.** Values are deterministic simulations of the same fitted parameter set, with Gq denoting fitted dose 5. The clamp imposes $r_{PV}(t) = 0$, including during baseline equilibration. Peaks are maxima of absolute latent population rates over the first 1 s after onset, computed at a 1-ms step and rounded to three significant digits. They are neither baseline-subtracted responses nor steady-state rates. Crossing times are the first sampled latencies above 40 Hz, and a dash indicates no crossing during the stimulus. This model-validity bound is not a physiological firing-rate ceiling. $\lambda_{max}$ is the largest real part of the rate–adaptation–plasticity Jacobian eigenvalues at the indicated fixed point. Negative values indicate local stability. The clamped Jacobian excludes the constrained PV rate.

### Contribution of reduced PV output efficacy to Gi-associated early calcium peaks

Under Gi, measured Pyr, PV and SST calcium responses showed enhanced early peaks (Figs. 4h–i and 5f), and the fitted model reproduced these peaks (Fig. 6c). Because Gi changed tonic PV input, PV output efficacy and softness together in the model, we asked how the fitted circuit generates these transients and whether reduced PV output efficacy is required for them. We first decomposed the inputs to Pyr, PV and SST in the fitted model (Supplementary Fig. S14a–i). Direct stimulus input predominantly targets Pyr, with weak inputs to PV and SST and none to VIP, so stimulus-evoked excitation of these interneurons depends strongly on recurrent Pyr activity. Under Gi, with the fitted PV efficacy factor $\kappa_{PV}^{(Gi)} \approx 0.383$, SST recruitment follows the initial Pyr response and recurrent excitation peaks before total recurrent inhibition. Gq shows no comparable transient amplification and retains a more Control-like temporal profile at reduced amplitude, qualitatively consistent with the measured response shapes.

We assessed the contribution of shunting by restoring $\kappa_{PV}^{(Gi)}$ to 1 without refitting the remaining Gi parameters or changing the calcium readout. After re-equilibrating the baseline, Pyr, PV and SST calcium responses lost their early peaks, decreased markedly in amplitude and rose gradually to stimulus offset (Supplementary Fig. S14j,k). Thus, attenuated PV output is required for the early calcium peaks of this fitted circuit.

We then tested whether other parameters could compensate by refitting the model with $\kappa_{PV}^{(Gi)}$ fixed at 1, using the same multistart optimization budget, objectives and bounds for all remaining trainable parameters. No candidate within the prescribed rate bound recovered early calcium peaks in all three populations. The best aggregate fit had an average error comparable to that of the fitted model but peaked at stimulus offset in every Gi trace. The only two candidates meeting all three early-peak criteria exhibited runaway dynamics and very large errors (Supplementary Table S9). Together, these interventions and refits support a key role for PV-output attenuation in reproducing the Gi early-peak phenotype within the tested model family.

| **Model or refit candidate** | **Mean NRMSE 0–1 s** | **Mean NRMSE 0–3 s** | **Gi calcium peak times Pyr / PV / SST (s)** | **Outcome** |
|---|---|---|---|---|
| Fitted model $\kappa_{PV}^{(Gi)} \approx 0.383$ | 0.2971 | 0.2212 | 0.101 / 0.121 / 0.179 | Early peaks; bounded dynamics |
| Without shunting: best aggregate fit | 0.3068 | 0.2168 | 1.000 / 1.000 / 1.000 | All peaks at stimulus offset |
| Without shunting: lowest-error candidate meeting all early-peak criteria | 2061.46 | 1112.80 | 0.193 / 0.212 / 0.186 | Runaway dynamics; invalid candidate |

**Supplementary Table S9. Refitting with Gi PV output efficacy fixed at the Control value.** Normalized root-mean-square error (NRMSE) is averaged across the nine Pyr/PV/SST × Control/Gi/Gq calcium traces; each trace's RMSE is normalized by its target range, with a floor of 0.05. Times and error windows are relative to stimulus onset. Early-peak criteria are ≤0.30 s for Pyr/PV and ≤0.20 s for SST. The invalid candidate reached $4.58 \times 10^5$ Hz, exceeding the prescribed 40-Hz rate limit. The best aggregate fit minimizes the sum of the two reported mean NRMSE values. NRMSE is a common reconstruction metric, not the composite optimization loss.

Both searches used Adam with 192, 144 and 80 starts and 1,800, 1,800 and 1,600 iterations in successive stages. The first stage was entirely random; the refinement stages included 24 and 16 fresh random starts, respectively. Remaining starts refined candidates from the preceding stage. All entries use independent simulations with a 1-ms time step rather than training-time trajectories.

### References


1. Campagnola, L. *et al.* Local connectivity and synaptic dynamics in mouse and human neocortex. *Science* **375**, eabj5861 (2022).
2. Pfeffer, C. K., Xue, M., He, M., Huang, Z. J. & Scanziani, M. Inhibition of inhibition in visual cortex: the logic of connections between molecularly distinct interneurons. *Nat Neurosci* **16**, 1068–1076 (2013).
3. Billeh, Y. N. *et al.* Systematic Integration of Structural and Functional Data into Multi-scale Models of Mouse Primary Visual Cortex. *Neuron* **106**, 388-403.e18 (2020).
4. Tsodyks, M. V., Skaggs, W. E., Sejnowski, T. J. & McNaughton, B. L. Paradoxical effects of external modulation of inhibitory interneurons. *J Neurosci* **17**, 4382–4388 (1997).
5. Sanzeni, A. *et al.* Inhibition stabilization is a widespread property of cortical networks. *Elife* **9**, e54875 (2020).

**Figure 1d - Number of Responsive Cells**

Model information:

| | |
|---|---|
| Number of observations | 34 |
| Fixed effects coefficients | 4 |
| Random effects coefficients | 11 |
| Covariance parameters | 1 |
| Distribution | Binomial |
| Link | Logit |
| FitMethod | REMPL |

Formula:

NumRespCells ~ 1 + Whisker_stim + (1 | MouseID)

Model fit statistics:

| AIC | BIC | LogLikelihood | Deviance |
|---|---|---|---|
| 114.14 | 121.15 | -52.071 | 104.14 |

Fixed effects coefficients (9 CIs):

| Name | Estimate | SE | tStat | DF | pValue | Lower | Upper | adj p |
|---|---|---|---|---|---|---|---|---|
| {'(Intercept)' } | 1.9473 | 0.30332 | 6.42 | 30 | 4.34E-07 | 1.3278 | 2.5668 | 4.34E-07 |
| {'Whisker_stim_Stm_D1'} | -2.1236 | 0.25846 | -8.2165 | 30 | 3.59E-09 | -2.6515 | -1.5958 | 7.17E-09 |
| {'Whisker_stim_Stm_B2'} | -2.5742 | 0.32875 | -7.8304 | 30 | 9.74E-09 | -3.2456 | -1.9028 | 1.30E-08 |
| {'Whisker_stim_Stm_E3'} | -5.2222 | 0.531 | -9.8346 | 30 | 6.74E-11 | -6.3067 | -4.1377 | 2.70E-10 |

**Figure 1h - MI of Responsive Cells**

Model information:

| | |
|---|---|
| Number of observations | 328 |
| Fixed effects coefficients | 4 |
| Random effects coefficients | 11 |
| Covariance parameters | 2 |

Formula:

MI ~ 1 + Whisker_stim + (1 | MouseID)

Model fit statistics:

| AIC | BIC | LogLikelihood | Deviance |
|---|---|---|---|
| 2172.3 | 2195 | -1080.2 | 2160.3 |

Fixed effects coefficients (9 CIs):

| Name | Estimate | SE | tStat | DF | pValue | Lower | Upper | adj p |
|---|---|---|---|---|---|---|---|---|
| {'(Intercept)' } | 8.3337 | 0.4901 | 17.004 | 324 | 8.53E-47 | 7.3696 | 9.2979 | 0 |
| {'Whisker_stim_Stm_D1'} | -4.4784 | 0.82183 | -5.4494 | 324 | 1.00E-07 | -6.0952 | -2.8617 | 0 |
| {'Whisker_stim_Stm_B2'} | -5.1862 | 1.1693 | -4.4353 | 324 | 1.26E-05 | -7.4866 | -2.8859 | 0 |
| {'Whisker_stim_Stm_E3'} | -3.6248 | 3.0052 | -1.2062 | 324 | 0.22864 | -9.537 | 2.2874 | 0.2286 |

**Figure 1i - MI of Non-Responsive Cells**

Model information:

| | |
|---|---|
| Number of observations | 292 |
| Fixed effects coefficients | 4 |
| Random effects coefficients | 11 |
| Covariance parameters | 2 |

Formula:

MI ~ 1 + Whisker_stim + (1 | MouseID)

Model fit statistics:

| AIC | BIC | LogLikelihood | Deviance |
|---|---|---|---|
| 656.62 | 678.59 | -322.31 | 644.62 |

Fixed effects coefficients (9 CIs):

| Name | Estimate | SE | tStat | DF | pValue | Lower | Upper | adj p |
|---|---|---|---|---|---|---|---|---|
| {'(Intercept)' } | 0.38062 | 0.18198 | 2.0915 | 288 | 0.037357 | 0.022436 | 0.73881 | 0.0498 |
| {'Whisker_stim_Stm_D1'} | 0.33018 | 0.1513 | 2.1824 | 288 | 0.02989 | 0.032397 | 0.62797 | 0.0498 |
| {'Whisker_stim_Stm_B2'} | -0.043217 | 0.1687 | -0.25617 | 288 | 0.798 | -0.37526 | 0.28883 | 0.798 |
| {'Whisker_stim_Stm_E3'} | -0.55869 | 0.16066 | -3.4775 | 288 | 0.00058415 | -0.8749 | -0.24248 | 0.0023 |

**Figure 2b - SI C1 vs D1 barrel @ baseline**

Linear mixed-effects model fit by ML                    *Adding random slopes did not improve model

Model information:

| | |
|---|---|
| Number of observations | 399 |
| Fixed effects coefficients | 2 |
| Random effects coefficients | 399 |
| Covariance parameters | 2 |

Formula:
SI_MI_BL ~ 1 + Barrel_imaged + (1 | MouseID:Index)

Model fit statistics:

| AIC | BIC | LogLikelihood | Deviance |
|---|---|---|---|
| 486.39 | 502.34 | -239.19 | 478.39 |

| Name | Estimate | SE | tStat | DF | pValue | Lower | Upper |
|---|---|---|---|---|---|---|---|
| {'(Intercept)' } | 0.45504 | 0.030409 | 14.964 | 397 | 1.84E-40 | 0.39526 | 0.51483 |
| {'Barrel_imaged_D1'} | -0.82844 | 0.044184 | -18.75 | 397 | 1.23E-56 | -0.9153 | -0.74158 |

**Figure 2c - % D1 Responsive cells in D1 barrel**

Model information:

| | |
|---|---|
| Number of observations | 66 |
| Fixed effects coefficients | 6 |
| Random effects coefficients | 11 |
| Covariance parameters | 1 |
| Distribution | Binomial |
| Link | Logit |
| FitMethod | MPL |

**ANOVA marginal tests: DFMethod = 'residual'**

| Term | FStat | DF1 | DF2 | pValue |
|---|---|---|---|---|
| {'(Intercept)' | 18.056 | 1 | 60 | 7.59E-05 |
| {'Timepoint' | 1.0245 | 5 | 60 | 0.41142 |

Formula:
NumRespCells ~ 1 + Timepoint + (1 | MouseID)

Model fit statistics:

| AIC | BIC | LogLikelihood | Deviance |
|---|---|---|---|
| 161.76 | 177.09 | -73.88 | 147.76 |

Fixed effects coefficients (95% CIs):

| Name | Estimate | SE | tStat | DF | pValue | Lower | Upper | adj p |
|---|---|---|---|---|---|---|---|---|
| {'(Intercept)' } | 1.0218 | 0.24047 | 4.2493 | 60 | 7.59E-05 | 0.54081 | 1.5028 | 0.0005 |
| {'Timepoint_Day3' } | 0.4804 | 0.25519 | 1.8825 | 60 | 0.064618 | -0.030058 | 0.99087 | 0.1939 |
| {'Timepoint_Day6' } | 0.30678 | 0.2485 | 1.2345 | 60 | 0.22182 | -0.19029 | 0.80385 | 0.4436 |
| {'Timepoint_Day12'} | 0.11741 | 0.2424 | 0.48438 | 60 | 0.62988 | -0.36747 | 0.6023 | 0.8106 |
| {'Timepoint_Day18'} | 0.057939 | 0.24073 | 0.24068 | 60 | 0.81062 | -0.42359 | 0.53947 | 0.8106 |
| {'Timepoint_Day46'} | 0.057939 | 0.24073 | 0.24068 | 60 | 0.81062 | -0.42359 | 0.53947 | 0.8106 |

**Figure 2c - % D1 Responsive cells in C1 barrel**

Model information:

| | |
|---|---|
| Number of observations | 66 |
| Fixed effects coefficients | 6 |
| Random effects coefficients | 11 |
| Covariance parameters | 1 |
| Distribution | Binomial |
| Link | Logit |
| FitMethod | MPL |

**ANOVA marginal tests: DFMethod = 'residual'**

| Term | FStat | DF1 | DF2 | pValue |
|---|---|---|---|---|
| {'(Intercept)' | 0.35327 | 1 | 60 | 0.55451 |
| {'Timepoint' | 12.031 | 5 | 60 | 4.37E-08 |

Formula:
NumRespCells ~ 1 + Timepoint + (1 | MouseID)

Model fit statistics:

| AIC | BIC | LogLikelihood | Deviance |
|---|---|---|---|
| 149.84 | 165.17 | -67.92 | 135.84 |

Fixed effects coefficien (95% CIs):

| Name | Estimate | SE | tStat | DF | pValue | Lower | Upper | adj p |
|---|---|---|---|---|---|---|---|---|
| {'(Intercept)' } | -0.10564 | 0.17774 | -0.59436 | 60 | 0.55451 | -0.46116 | 0.24989 | 0.5545 |
| {'Timepoint_Day3' } | -0.35644 | 0.19954 | -1.7863 | 60 | 0.079108 | -0.75558 | 0.042707 | 0.1187 |
| {'Timepoint_Day6' } | 0.2927 | 0.1979 | 1.4791 | 60 | 0.14435 | -0.10315 | 0.68855 | 0.1732 |
| {'Timepoint_Day12'} | 0.67814 | 0.20167 | 3.3626 | 60 | 0.001349 | 0.27473 | 1.0815 | 0.0081 |
| {'Timepoint_Day18'} | 0.49204 | 0.19939 | 2.4677 | 60 | 0.016468 | 0.093193 | 0.89088 | 0.0329 |
| {'Timepoint_Day46'} | -0.62946 | 0.2033 | -3.0962 | 60 | 0.0029786 | -1.0361 | -0.2228 | 0.0089 |

**Figure 2d - MI of D1 responsive cells in D1 barrel**

Linear mixed-effects model fit by REML

Model information:

| | |
|---|---|
| Number of observations | 867 |
| Fixed effects coefficients | 6 |
| Random effects coefficients | 1032 |
| Covariance parameters | 22 |

**ANOVA marginal tests: DFMethod = 'residual'**

| Term | FStat | DF1 | DF2 | pValue |
|---|---|---|---|---|
| {'(Intercept)' | 135.07 | 1 | 861 | 4.17E-29 |
| {'Timepoint' | 2.4741 | 5 | 861 | 0.030852 |

Formula:
MI ~ 1 + Timepoint + (1 + Timepoint | MouseID:Index)

Model fit statistics:

| AIC | BIC | LogLikelihood | Deviance |
|---|---|---|---|
| 5568.4 | 5701.6 | -2756.2 | 5512.4 |

Fixed effects coefficients (95% CIs):

| Name | Estimate | SE | tStat | DF | pValue | Lower | Upper | adj p |
|---|---|---|---|---|---|---|---|---|
| {'(Intercept)' } | 6.189 | 0.53253 | 11.622 | 861 | 4.17E-29 | 5.1438 | 7.2342 | 0 |
| {'Timepoint_Day3' } | 0.8028 | 0.46014 | 1.7447 | 861 | 0.081398 | -0.10033 | 1.7059 | 0.1221 |
| {'Timepoint_Day6' } | 1.3761 | 0.62983 | 2.1849 | 861 | 0.029167 | 0.13991 | 2.6123 | 0.0583 |
| {'Timepoint_Day12'} | 0.48865 | 0.58217 | 0.83935 | 861 | 0.4015 | -0.654 | 1.6313 | 0.4015 |
| {'Timepoint_Day18'} | 1.3325 | 0.49115 | 2.713 | 861 | 0.0067999 | 0.36852 | 2.2965 | 0.0204 |
| {'Timepoint_Day46'} | 0.79954 | 0.85735 | 0.93257 | 861 | 0.3513 | -0.88319 | 2.4823 | 0.4015 |

**Figure 2d - MI of D1 responsive cells in C1 barrel**

Model information:

| | |
|---|---|
| Number of observations | 631 |
| Fixed effects coefficients | 6 |
| Random effects coefficients | 1044 |
| Covariance parameters | 22 |

**ANOVA marginal tests: DFMethod = 'residual'**

| Term | FStat | DF1 | DF2 | pValue |
|---|---|---|---|---|
| {'(Intercept)' | 39.835 | 1 | 625 | 5.24E-10 |
| {'Timepoint' | 1.0989 | 5 | 625 | 0.35977 |

Formula:
MI ~ 1 + Timepoint + (1 + Timepoint | MouseID:Index)

Model fit statistics:

| AIC | BIC | LogLikelihood | Deviance |
|---|---|---|---|
| 2941.7 | 3065.9 | -1442.8 | 2885.7 |

Fixed effects coefficien (95% CIs):

| Name | Estimate | SE | tStat | DF | pValue | Lower | Upper | adj p |
|---|---|---|---|---|---|---|---|---|
| {'(Intercept)' } | 2.8232 | 0.44731 | 6.3115 | 625 | 5.24E-10 | 1.9448 | 3.7016 | 0 |
| {'Timepoint_Day3' } | 0.19054 | 0.45109 | 0.4224 | 625 | 0.67288 | -0.69529 | 1.0764 | 0.7084 |
| {'Timepoint_Day6' } | 0.29629 | 0.41073 | 0.72139 | 625 | 0.47094 | -0.51028 | 1.1029 | 0.7064 |
| {'Timepoint_Day12'} | 0.38835 | 0.34674 | 1.12 | 625 | 0.26314 | -0.29256 | 1.0693 | 0.7064 |
| {'Timepoint_Day18'} | 0.35627 | 0.40197 | 0.88632 | 625 | 0.37579 | -0.43311 | 1.1457 | 0.7064 |
| {'Timepoint_Day46'} | -0.17592 | 0.47008 | -0.37422 | 625 | 0.70836 | -1.099 | 0.74722 | 0.7084 |

**Figure 2g - delta MI By Switch category D1 barrel - BL vs D12**

Kruskal Wallis 6.44E-04

| Group A | Group B | Lower Limit | A-B | Upper Limit | P-value |
|---|---|---|---|---|---|
| NR to NR | NR to R | -88.770248 | -45.4 | -2.029752 | 0.03604257 |
| NR to NR | R to NR | -4.6865848 | 43.8909091 | 92.468403 | 0.09321837 |
| NR to NR | R to R | -29.983139 | -3.1765625 | 23.6300138 | 0.99021131 |
| NR to R | R to NR | 33.5041339 | 89.2909091 | 145.077684 | 0.00022949 |
| NR to R | R to R | 3.87002959 | 42.2234375 | 80.5768454 | 0.02419532 |
| R to NR | R to R | -91.223767 | -47.067472 | -2.9111764 | 0.03138255 |

**Figure 2h - delta MI By Switch category C1 barrel - BL vs D12**

Kruskal Wallis 6.50E-10

| Group A | Group B | Lower Limit | A-B | Upper Limit | P-value |
|---|---|---|---|---|---|
| NR to NR | NR to R | -92.899028 | -62.706589 | -32.51415 | 5.72E-07 |
| NR to NR | R to NR | -0.693579 | 46.926799 | 94.547177 | 0.05517776 |
| NR to NR | R to R | -43.281032 | -17.396628 | 8.48777683 | 0.30973966 |
| NR to R | R to NR | 60.7135795 | 109.633388 | 158.553196 | 5.47E-08 |
| NR to R | R to R | 17.1062444 | 45.3099613 | 73.5136782 | 0.00021489 |
| R to NR | R to R | -110.70841 | -64.323427 | -17.938444 | 0.00208074 |

**Figure 2i - MI @ D12 By Switch category D1 barrel**

Kruskal Wallis 7.01E-21

| Group A | Group B | Lower Limit | A-B | Upper Limit | P-value |
|---|---|---|---|---|---|
| NR to NR | NR to R | -96.360724 | -52.990476 | -9.6202282 | 0.0091962 |
| NR to NR | R to NR | -72.36191 | -23.784416 | 24.7930783 | 0.58989451 |
| NR to NR | R to R | -123.47309 | -96.666518 | -69.859942 | 3.77E-09 |
| NR to R | R to NR | -26.580715 | 29.2060606 | 84.9928357 | 0.53416811 |
| NR to R | R to R | -82.02945 | -43.676042 | -5.3226338 | 0.01807376 |
| R to NR | R to R | -117.0384 | -72.882102 | -28.725807 | 0.00013116 |

**Figure 2j - MI @ D12 By Switch category C1 barrel**

Kruskal Wallis 1.58E-29

| Group A | Group B | Lower Limit | A-B | Upper Limit | P-value |
|---|---|---|---|---|---|
| NR to NR | NR to R | -107.56375 | -77.371311 | -47.178872 | 4.03E-09 |
| NR to NR | R to NR | -68.292835 | -20.672457 | 26.9479214 | 0.68011937 |
| NR to NR | R to R | -139.96945 | -114.08504 | -88.20064 | 3.77E-09 |
| NR to R | R to NR | 7.77904592 | 56.6988543 | 105.618663 | 0.01539962 |
| NR to R | R to R | -64.91745 | -36.713733 | -8.5100162 | 0.00458286 |
| R to NR | R to R | -139.79757 | -93.412587 | -47.027604 | 1.37E-06 |

**Figure 3c - Change in SI BL vs D46 - D1 Barrel**

Linear mixed-effects model fit by ML

Model information:

| | |
|---|---|
| Number of observations | 378 |
| Fixed effects coefficients | 2 |
| Random effects coefficients | 378 |
| Covariance parameters | 4 |

Formula:

SI ~ 1 + Timepoint + (1 + Timepoint | MouseID:Index)

Model fit statistics:

| AIC | BIC | LogLikelihood | Deviance |
|---|---|---|---|
| 431.82 | 455.43 | -209.91 | 419.82 |

| Name | Estimate | SE | tStat | DF | pValue | Lower | Upper |
|---|---|---|---|---|---|---|---|
| {'(Intercept)' } | -0.3734 | 0.034053 | -10.965 | 376 | 1.84E-24 | -0.44035 | -0.30644 |
| {'Timepoint_D46'} | -0.28972 | 0.037199 | -7.7882 | 376 | 6.68E-14 | -0.36286 | -0.21657 |

**Figure 3d - Change in SI BL vs D46 - C1 Barrel**

Linear mixed-effects model fit by ML

Model information:

| | |
|---|---|
| Number of observations | 420 |
| Fixed effects coefficients | 2 |
| Random effects coefficients | 420 |
| Covariance parameters | 4 |

Formula:

SI ~ 1 + Timepoint + (1 + Timepoint | MouseID:Index)

Model fit statistics:

| AIC | BIC | LogLikelihood | Deviance |
|---|---|---|---|
| 499.25 | 523.49 | -243.62 | 487.25 |

| Name | Estimate | SE | tStat | DF | pValue | Lower | Upper |
|---|---|---|---|---|---|---|---|
| {'(Intercept)' } | 0.45504 | 0.028596 | 15.913 | 418 | 6.48E-45 | 0.39883 | 0.51125 |
| {'Timepoint_D46'} | -0.066696 | 0.032832 | -2.0315 | 418 | 0.04284 | -0.13123 | -0.0021607 |

**Figure 3e - Change in %Responsive cells by whisker - BL vs D46 - D1 Barrel**

GLME

Model information:

| | |
|---|---|
| Number of observations | 44 |
| Fixed effects coefficients | 4 |
| Random effects coefficients | 11 |
| Covariance parameters | 1 |
| Distribution | Binomial |
| Link | Logit |
| FitMethod | REMPL |

ANOVA marg = 'residual'

| Term | FStat | DF1 | DF2 | pValue |
|---|---|---|---|---|
| {'(Intercept)' } | 8.2822 | 1 | 40 | 0.0063956 |
| {'Timepoint' } | 15.629 | 1 | 40 | 0.00030598 |
| {'Whisker_stim' } | 53.739 | 1 | 40 | 6.52E-09 |
| {'Timepoint:Whisker_stim | 9.1048 | 1 | 40 | 0.0044192 |

Formula:

NumRespCells ~ 1 + Timepoint*Whisker_stim + (1 | MouseID)

Model fit statistics:

| AIC | BIC | LogLikelihood | Deviance |
|---|---|---|---|
| 123.97 | 132.41 | -56.983 | 113.97 |

Fixed effects coefficients (95% CIs):

| Name | Estimate | SE | tStat | DF | pValue | Lower | Upper | adj p |
|---|---|---|---|---|---|---|---|---|
| {'(Intercept)' } | -0.59996 | 0.20847 | -2.8779 | 40 | 0.0063956 | -1.0213 | -0.17862 | 0.0064 |
| {'Timepoint_Day46' } | -0.98048 | 0.24802 | -3.9533 | 40 | 0.00030598 | -1.4817 | -0.47922 | 0.0006 |
| {'Whisker_stim_Stm_D1' } | 1.6758 | 0.2286 | 7.3307 | 40 | 6.52E-09 | 1.2138 | 2.1378 | 0 |
| {'Timepoint_Day46:Whisker_stim_Stm_D1'} | 1.0371 | 0.3437 | 3.0174 | 40 | 0.0044192 | 0.34244 | 1.7317 | 0.0059 |

**Figure 3f - Change in MI of responsive cells BL vs D46 - D1 barrel**

Model information:

| | |
|---|---|
| Number of observations | 756 |
| Fixed effects coefficients | 4 |
| Random effects coefficients | 11 |
| Covariance parameters | 2 |

ANOVA marg = 'Residual'

| Term | FStat | DF1 | DF2 | pValue |
|---|---|---|---|---|
| {'(Intercept)' } | 17.972 | 1 | 752 | 2.52E-05 |
| {'Timepoint' } | 3.4121 | 1 | 752 | 0.065111 |
| {'Whisker_stim' } | 25.784 | 1 | 752 | 4.82E-07 |
| {'Timepoint:Whisker_stim | 4.514 | 1 | 752 | 0.033945 |

Formula:

MI ~ 1 + Timepoint*Whisker_stim + (1 | MouseID)

Model fit statistics:

| AIC | BIC | LogLikelihood | Deviance |
|---|---|---|---|
| 4938.1 | 4965.8 | -2463 | 4926.1 |

Fixed effects coefficients (95% CIs):

| Name | Estimate | SE | tStat | DF | pValue | Lower | Upper | adj p |
|---|---|---|---|---|---|---|---|---|
| {'(Intercept)' } | 1.9464 | 0.45913 | 4.2393 | 752 | 2.52E-05 | 1.0451 | 2.8477 | 0.0001 |
| {'Timepoint_Day46' } | -1.1994 | 0.64931 | -1.8472 | 752 | 0.065111 | -2.4741 | 0.075273 | 0.0651 |
| {'Whisker_stim_Stm_D1' } | 3.297 | 0.64931 | 5.0777 | 752 | 4.82E-07 | 2.0224 | 4.5717 | 0 |
| {'Timepoint_Day46:Whisker_stim_Stm_D1'} | 1.9509 | 0.91826 | 2.1246 | 752 | 0.033945 | 0.14829 | 3.7536 | 0.0453 |

**Figure 3e - Change in %Responsive cells by whisker - BL vs D46 - C1 Barrel**

GLME

Model information:

| | |
|---|---|
| Number of observations | 68 |
| Fixed effects coefficients | 8 |
| Random effects coefficients | 11 |
| Covariance parameters | 1 |
| Distribution | Binomial |
| Link | Logit |
| FitMethod | REMPL |

ANOVA marg = 'residual'

| Term | FStat | DF1 | DF2 | pValue |
|---|---|---|---|---|
| {'(Intercept)' } | 62.856 | 1 | 60 | 6.50E-11 |
| {'Timepoint' } | 10.948 | 1 | 60 | 0.001588 |
| {'Whisker_stim' } | 43.553 | 3 | 60 | 4.51E-15 |
| {'Timepoint:Whisker_stim | 4.6229 | 3 | 60 | 0.0056436 |

Formula:

NumRespCells ~ 1 + Timepoint*Whisker_stim + (1 | MouseID)

Model fit statistics:

| AIC | BIC | LogLikelihood | Deviance |
|---|---|---|---|
| 394.71 | 413.56 | -188.35 | 376.71 |

Fixed effects coefficients (95% CIs):

| Name | Estimate | SE | tStat | DF | pValue | Lower | Upper | adj p |
|---|---|---|---|---|---|---|---|---|
| {'(Intercept)' } | 1.931 | 0.24357 | 7.9282 | 60 | 6.50E-11 | 1.4438 | 2.4183 | 0 |
| {'Timepoint_Day46' } | -0.86522 | 0.2615 | -3.3087 | 60 | 0.001588 | -1.3883 | -0.34215 | 0.0021 |
| {'Whisker_stim_Stm_D1' } | -2.0269 | 0.2504 | -8.0946 | 60 | 3.38E-11 | -2.5277 | -1.526 | 0 |
| {'Whisker_stim_Stm_B2' } | -2.4497 | 0.30408 | -8.0561 | 60 | 3.93E-11 | -3.0579 | -1.8414 | 0 |
| {'Whisker_stim_Stm_E3' } | -5.0054 | 0.51213 | -9.7735 | 60 | 5.05E-14 | -6.0298 | -3.9809 | 0 |
| {'Timepoint_Day46:Whisker_stim_Stm_D1'} | 0.2346 | 0.33128 | 0.70816 | 60 | 0.48159 | -0.42807 | 0.89727 | 0.5504 |
| {'Timepoint_Day46:Whisker_stim_Stm_B2'} | -2.6427 | 0.79022 | -3.3443 | 60 | 0.0014262 | -4.2234 | -1.062 | 0.0021 |
| {'Timepoint_Day46:Whisker_stim_Stm_E3'} | -99.708 | 6.71E+06 | -1.49E-05 | 60 | 0.99999 | -1.34E+07 | 1.34E+07 | 1 |

**Figure 3f - Change in MI of responsive cells BL vs D46 - C1 barrel**

Model information:

| | |
|---|---|
| Number of observations | 1240 |
| Fixed effects coefficients | 8 |
| Random effects coefficients | 11 |
| Covariance parameters | 2 |

ANOVA marg = 'Residual'

| Term | FStat | DF1 | DF2 | pValue |
|---|---|---|---|---|
| {'(Intercept)' } | 543.91 | 1 | 1232 | 6.09E-100 |
| {'Timepoint' } | 32.962 | 1 | 1232 | 1.18E-08 |
| {'Whisker_stim' } | 79.005 | 3 | 1232 | 9.58E-47 |
| {'Timepoint:Whisker_stim | 3.5724 | 3 | 1232 | 0.013618 |

Formula:

MI ~ 1 + Timepoint*Whisker_stim + (1 | MouseID)

Model fit statistics:

| AIC | BIC | LogLikelihood | Deviance |
|---|---|---|---|
| 7284.1 | 7335.2 | -3632 | 7264.1 |

Fixed effects coefficients (95% CIs):

| Name | Estimate | SE | tStat | DF | pValue | Lower | Upper | adj p |
|---|---|---|---|---|---|---|---|---|
| {'(Intercept)' } | 7.307 | 0.31331 | 23.322 | 1232 | 6.09E-100 | 6.6923 | 7.9216 | 0 |
| {'Timepoint_Day46' } | -2.5439 | 0.44308 | -5.7413 | 1232 | 1.18E-08 | -3.4132 | -1.6746 | 0 |
| {'Whisker_stim_Stm_D1' } | -5.0767 | 0.44308 | -11.458 | 1232 | 5.94E-29 | -5.946 | -4.2074 | 0 |
| {'Whisker_stim_Stm_B2' } | -5.8373 | 0.55164 | -10.582 | 1232 | 4.17E-25 | -6.9195 | -4.755 | 0 |
| {'Whisker_stim_Stm_E3' } | -7.1107 | 0.55164 | -12.89 | 1232 | 9.33E-36 | -8.193 | -6.0285 | 0 |
| {'Timepoint_Day46:Whisker_stim_Stm_D1'} | 1.7073 | 0.62662 | 2.7246 | 1232 | 0.0065289 | 0.47793 | 2.9366 | 0.0075 |
| {'Timepoint_Day46:Whisker_stim_Stm_B2'} | 1.2495 | 0.78013 | 1.6016 | 1232 | 0.10949 | -0.28104 | 2.78 | 0.1095 |
| {'Timepoint_Day46:Whisker_stim_Stm_E3'} | 2.1415 | 0.78013 | 2.745 | 1232 | 0.0061386 | 0.61095 | 3.672 | 0.0075 |

**Figure 4d - hM3D(Gq) - PV Cells - % Responsive**
Paired t-test 0.0015

**Figure 4d - hM3D(Gq) - Pyr Cells - % Responsive**
Paired t-test 9.79E-05

**Figure 4e - hM3D(Gq) - PV Cells - MI**
Linear mixed-effects model fit by REML

Model information:
Number of observations 184
Fixed effects coefficients 2
Random effects coefficients 92
Covariance parameters 2

Formula:
MI_dF ~ 1 + condition + (1 | MouseID:index)

Model fit statistics:

| AIC | BIC | LogLikelihood | Deviance |
|---|---|---|---|
| -399.87 | -387.05 | 203.93 | -407.87 |

| Name | Estimate | SE | tStat | DF | pValue | Lower | Upper |
|---|---|---|---|---|---|---|---|
| {'(Intercept)' } | 0.13664 | 0.0082346 | 16.593 | 182 | 2.92E-38 | 0.12039 | 0.15288 |
| {'condition_C21'} | -0.11042 | 0.0096522 | -11.44 | 182 | 3.52E-23 | -0.12946 | -0.091373 |

**Figure 4f - hM3D(Gq) - Pyr Cells - MI**
Linear mixed-effects model fit by REML

Model information:
Number of observations 1480
Fixed effects coefficients 2
Random effects coefficients 7
Covariance parameters 2

Formula:
MI_dF ~ 1 + condition + (1 | MouseID)

Model fit statistics:

| AIC | BIC | LogLikelihood | Deviance |
|---|---|---|---|
| -429.85 | -408.66 | 218.93 | -437.85 |

| Name | Estimate | SE | tStat | DF | pValue | Lower | Upper |
|---|---|---|---|---|---|---|---|
| {'(Intercept)' } | 0.092244 | 0.0072853 | 12.662 | 1478 | 5.87E-35 | 0.077953 | 0.10653 |
| {'condition_C21'} | -0.081541 | 0.010852 | -7.5138 | 1478 | 9.90E-14 | -0.10283 | -0.060253 |

**Figure 4g - hM4D(Gi) - PV Cells - % Responsive**
Paired t-test 0.4097

**Figure 4g - hM4D(Gi) - Pyr Cells - % Responsive**
Paired t-test 0.8479

**Figure 4h - hM4D(Gi) - PV Cells - MI 1-2 s**
Linear mixed-effects model fit by REML

Model information:
Number of observations 264
Fixed effects coefficients 2
Random effects coefficients 132
Covariance parameters 2

Formula:
MIearly_dF ~ 1 + condition + (1 | MouseID:index)

Model fit statistics:

| AIC | BIC | LogLikelihood | Deviance |
|---|---|---|---|
| -268.05 | -253.77 | 138.02 | -276.05 |

| Name | Estimate | SE | tStat | DF | pValue | Lower | Upper |
|---|---|---|---|---|---|---|---|
| {'(Intercept)' } | 0.1545 | 0.012677 | 12.187 | 262 | 2.29E-27 | 0.12954 | 0.17946 |
| {'condition_C21'} | 0.072483 | 0.014176 | 5.1131 | 262 | 6.12E-07 | 0.04457 | 0.1004 |

**Figure 4h - hM4D(Gi) - PV Cells - MI 2-3 s**
Linear mixed-effects model fit by REML

Model information:
Number of observations 264
Fixed effects coefficients 2
Random effects coefficients 132
Covariance parameters 2

Formula:
MIlate_dF ~ 1 + condition + (1 | MouseID:index)

Model fit statistics:

| AIC | BIC | LogLikelihood | Deviance |
|---|---|---|---|
| -288.02 | -273.74 | 148.01 | -296.02 |

| Name | Estimate | SE | tStat | DF | pValue | Lower | Upper |
|---|---|---|---|---|---|---|---|
| {'(Intercept)' } | 0.17171 | 0.0139 | 12.353 | 262 | 6.23E-28 | 0.14434 | 0.19908 |
| {'condition_C21'} | -0.058605 | 0.010775 | -5.4391 | 262 | 1.23E-07 | -0.079822 | -0.037389 |

**Figure 4i - hM4D(Gi) - Pyr Cells - MI**
Linear mixed-effects model fit by REML

Model information:
Number of observations 2092
Fixed effects coefficients 2
Random effects coefficients 8
Covariance parameters 2

Formula:
MI_dF ~ 1 + condition + (1 | MouseID)

Model fit statistics:

| AIC | BIC | LogLikelihood | Deviance |
|---|---|---|---|
| -123.33 | -100.75 | 65.666 | -131.33 |

| Name | Estimate | SE | tStat | DF | pValue | Lower | Upper |
|---|---|---|---|---|---|---|---|
| {'(Intercept)' } | 0.089146 | 0.020289 | 4.3938 | 2090 | 1.17E-05 | 0.049357 | 0.12894 |
| {'condition_C21'} | 0.073513 | 0.010208 | 7.2016 | 2090 | 8.26E-13 | 0.053495 | 0.093532 |

**Figure 4j - mCherry - PV Cells - % Responsive**
Paired t-test 0.3999

**Figure 4j - mCherry - Pyr Cells - % Responsive**
Paired t-test 0.7089

**Figure 4k - mCherry - PV Cells - MI**
Linear mixed-effects model fit by REML

Model information:
Number of observations 142
Fixed effects coefficients 2
Random effects coefficients 71
Covariance parameters 2

Formula:
MI_dF ~ 1 + condition + (1 | MouseID:index)

Model fit statistics:

| AIC | BIC | LogLikelihood | Deviance |
|---|---|---|---|
| -134.77 | -123.01 | 71.387 | -142.77 |

| Name | Estimate | SE | tStat | DF | pValue | Lower | Upper |
|---|---|---|---|---|---|---|---|
| {'(Intercept)' } | 0.21105 | 0.020543 | 10.274 | 140 | 8.45E-19 | 0.17044 | 0.25166 |
| {'condition_C21'} | -0.018158 | 0.014569 | -1.2463 | 140 | 2.15E-01 | -0.046962 | 0.010646 |

**Figure 4l - mCherry - Pyr Cells - MI**
Linear mixed-effects model fit by REML

Model information:
Number of observations 1215
Fixed effects coefficients 2
Random effects coefficients 6
Covariance parameters 2

Formula:
MI_dF ~ 1 + condition + (1 | MouseID)

Model fit statistics:

| AIC | BIC | LogLikelihood | Deviance |
|---|---|---|---|
| 27.756 | 48.16 | -9.8781 | 19.756 |

| Name | Estimate | SE | tStat | DF | pValue | Lower | Upper |
|---|---|---|---|---|---|---|---|
| {'(Intercept)' } | 0.093275 | 0.0099233 | 9.3996 | 1213 | 2.63E-20 | 0.073806 | 0.11274 |
| {'condition_C21'} | 0.0043966 | 0.013925 | 0.31573 | 1213 | 0.75226 | -0.022923 | 0.031717 |

**Figure 5c - hM3D(Gq) - SST - % Responsive**

Wilcoxin Signed Ra 9.77E-04

**Figure 5d - hM3D(Gq) - MI**

Linear mixed-effects model fit by REML

Model information:
Number of observations 274
Fixed effects coefficients 2
Random effects coefficients 14
Covariance parameters 2

Formula:
MI_dF ~ 1 + condition + (1 | MouseID)

Model fit statistics:

| AIC | BIC | LogLikelihood | Deviance |
|---|---|---|---|
| -515.09 | -500.66 | 261.54 | -523.09 |

| Name | Estimate | SE | tStat | DF | pValue | Lower | Upper |
|---|---|---|---|---|---|---|---|
| {'(Intercept)' } | 0.056226 | 0.0077616 | 7.2442 | 272 | 4.49E-12 | 0.040946 | 0.071506 |
| {'condition_C21'} | -0.05144 | 0.010976 | -4.6864 | 272 | 4.40E-06 | -0.07305 | -0.02983 |

**Figure 5e - hM4D(Gi) - SST - % Responsive**

Wilcoxin Signed Ra 0.332

**Figure 5f - hM4D(Gi) - MI 1-2 s**

Linear mixed-effects model fit by REML

Model information:
Number of observations 350
Fixed effects coefficients 2
Random effects coefficients 16
Covariance parameters 2

Formula:
MIearly_dF ~ 1 + condition + (1 | MouseID)

Model fit statistics:

| AIC | BIC | LogLikelihood | Deviance |
|---|---|---|---|
| -1210.6 | -1195.2 | 609.32 | -1218.6 |

| Name | Estimate | SE | tStat | DF | pValue | Lower | Upper |
|---|---|---|---|---|---|---|---|
| {'(Intercept)' } | 0.01227 | 0.005311 | 2.3103 | 348 | 0.021457 | 0.0018242 | 0.022716 |
| {'condition_C21'} | 0.013651 | 0.0042814 | 3.1884 | 348 | 0.0015606 | 0.00523 | 0.022071 |

**Figure 5f - hM4D(Gi) - MI 2-3 s**

Linear mixed-effects model fit by REML

Model information:
Number of observations 350
Fixed effects coefficients 2
Random effects coefficients 16
Covariance parameters 2

Formula:
MIlate_dF ~ 1 + condition + (1 | MouseID)

Model fit statistics:

| AIC | BIC | LogLikelihood | Deviance |
|---|---|---|---|
| -1163 | -1147.6 | 585.52 | -1171 |

| Name | Estimate | SE | tStat | DF | pValue | Lower | Upper |
|---|---|---|---|---|---|---|---|
| {'(Intercept)' } | 0.018144 | 0.0033503 | 5.4156 | 348 | 1.14E-07 | 0.011554 | 0.024733 |
| {'condition_C21'} | -0.01201 | 0.004738 | -2.5349 | 348 | 0.011685 | -0.021329 | -0.0026917 |

**Figure 5g - mCherry - SST - % Responsive**

Paired T-test 0.6147

**Figure 5h - hM3D(Gq) - MI**

Linear mixed-effects model fit by REML

Model information:
Number of observations 160
Fixed effects coefficients 2
Random effects coefficients 12
Covariance parameters 2

Formula:
MI_dF ~ 1 + condition + (1 | MouseID)

Model fit statistics:

| AIC | BIC | LogLikelihood | Deviance |
|---|---|---|---|
| -258.11 | -245.85 | 133.05 | -266.11 |

| Name | Estimate | SE | tStat | DF | pValue | Lower | Upper |
|---|---|---|---|---|---|---|---|
| {'(Intercept)' } | 0.036708 | 0.011336 | 3.2382 | 158 | 1.47E-03 | 0.014319 | 0.059097 |
| {'condition_C21'} | 0.0060808 | 0.016031 | 0.37931 | 158 | 0.70497 | -0.025582 | 0.037744 |

**Figure S1e - Percent Responsive Cells by stimulus type**

Model information:

| | |
|---|---|
| Number of observations | 72 |
| Fixed effects coefficients | 12 |
| Random effects coefficients | 6 |
| Covariance parameters | 1 |
| Distribution | Binomial |
| Link | Logit |
| FitMethod | REMPL |

ANOVA

| Term | FStat | DF1 | DF2 | pValue |
|---|---|---|---|---|
| {'(Intercept)' | 0.51501 | 1 | 60 | 0.47576 |
| {'condition' | 2.8846 | 1 | 60 | 0.094609 |
| {'stim_amp' | 11.747 | 5 | 60 | 6.15E-08 |
| {'condition:st | 2.9015 | 5 | 60 | 0.020646 |

Formula:

NumRespCells ~ 1 + condition*stim_amp + (1 | MouseID)

Model fit statistics:

| AIC | BIC | LogLikelihood | Deviance |
|---|---|---|---|
| 211.92 | 239.14 | -92.958 | 185.92 |

| Name | Estimate | SE | tStat | DF | pValue | Lower | Upper | adj p |
|---|---|---|---|---|---|---|---|---|
| {'(Intercept)' } | -0.29043 | 0.4047 | -0.71764 | 60 | 0.47576 | -1.1 | 0.51909 | 0.5581 |
| {'condition_Awake' } | 0.45066 | 0.26534 | 1.6984 | 60 | 0.094609 | -0.080101 | 0.98142 | 0.1622 |
| {'stim_amp_Amp2_Spd0.01' } | -0.61677 | 0.27127 | -2.2736 | 60 | 0.02658 | -1.1594 | -0.07415 | 0.0532 |
| {'stim_amp_Amp2' } | -1.6163 | 0.30623 | -5.2779 | 60 | 1.90E-06 | -2.2288 | -1.0037 | 0 |
| {'stim_amp_Amp4' } | -1.1557 | 0.2854 | -4.0492 | 60 | 0.00014959 | -1.7266 | -0.58477 | 0.0009 |
| {'stim_amp_Amp6' } | -0.69395 | 0.27272 | -2.5445 | 60 | 0.013534 | -1.2395 | -0.14842 | 0.0406 |
| {'stim_amp_Amp8' } | 0.35381 | 0.26658 | 1.3272 | 60 | 0.18947 | -0.17943 | 0.88705 | 0.2526 |
| {'condition_Awake:stim_amp_Amp2_Spd0.01'} | -0.91527 | 0.39055 | -2.3436 | 60 | 0.022432 | -1.6965 | -0.13406 | 0.0532 |
| {'condition_Awake:stim_amp_Amp2' } | -1.6373 | 0.51431 | -3.1834 | 60 | 0.0023077 | -2.666 | -0.60848 | 0.0092 |
| {'condition_Awake:stim_amp_Amp4' } | -0.65603 | 0.40757 | -1.6096 | 60 | 0.11273 | -1.4713 | 0.15924 | 0.1691 |
| {'condition_Awake:stim_amp_Amp6' } | -0.20938 | 0.38133 | -0.54908 | 60 | 0.58499 | -0.97214 | 0.55339 | 0.585 |
| {'condition_Awake:stim_amp_Amp8' } | -0.24829 | 0.37604 | -0.66027 | 60 | 0.51161 | -1.0005 | 0.5039 | 0.5581 |

**Figure S1g - MI for Awake vs Anesth.**

Model information:

| | |
|---|---|
| Number of observations | 136 |
| Fixed effects coefficients | 2 |
| Random effects coefficients | 6 |
| Covariance parameters | 2 |

Formula:

MI ~ 1 + condition + (1 | MouseID)

Model fit statistics:

| AIC | BIC | LogLikelihood | Deviance |
|---|---|---|---|
| 654.79 | 666.38 | -323.39 | 646.79 |

| Name | Estimate | SE | tStat | DF | pValue | Lower | Upper | adj p |
|---|---|---|---|---|---|---|---|---|
| {'(Intercept)' } | 3.1054 | 0.50613 | 6.1357 | 134 | 8.95E-09 | 2.1044 | 4.1065 | 0 |
| {'condition_Awake'} | -0.20844 | 0.44346 | -0.47003 | 134 | 0.6391 | -1.0855 | 0.66865 | 0.6391 |

**Figure S2a - %Whisker Responsive Cells in D1 Barrel**
Generalized linear mixed-effects model fit by PL

Model information:
Number of observations 22
Fixed effects coefficients 2
Random effects coefficients 11
Covariance parameters 1
Distribution Binomial
Link Logit
FitMethod REMPL

Formula:
NumRespCells ~ 1 + Whisker_stim + (1 | MouseID)

Model fit statistics:

| AIC | BIC | LogLikelihood | Deviance |
|---|---|---|---|
| 58.686 | 61.673 | -26.343 | 52.686 |

| Name | Estimate | SE | tStat | DF | pValue | Lower | Upper |
|---|---|---|---|---|---|---|---|
| {'(Intercept)' } | -0.61164 | 0.20182 | -3.0305 | 20 | 0.0066052 | -1.0326 | -0.19064 |
| {'Whisker_stim_Stm_D1'} | 1.6578 | 0.22745 | 7.2887 | 20 | 4.76E-07 | 1.1834 | 2.1323 |

**Figure S2d - MI of Responsive Cells**
Linear mixed-effects model fit by REML

Model information:
Number of observations 206
Fixed effects coefficients 2
Random effects coefficients 11
Covariance parameters 2

Formula:
MI ~ 1 + Whisker_stim + (1 | MouseID)

Model fit statistics:

| AIC | BIC | LogLikelihood | Deviance |
|---|---|---|---|
| 1312.7 | 1325.9 | -652.34 | 1304.7 |

Fixed effects coefficients (9 CIs):

| Name | Estimate | SE | tStat | DF | pValue | Lower | Upper | adj. p |
|---|---|---|---|---|---|---|---|---|
| {'(Intercept)' } | 4.084 | 0.70754 | 5.7721 | 204 | 2.88E-08 | 2.689 | 5.4791 | 0 |
| {'Whisker_stim_Stm_D1'} | 2.834 | 0.86135 | 3.2902 | 204 | 0.0011796 | 1.1357 | 4.5323 | 0.0012 |

**Figure S2e - MI of Non-Responsive Cells**
Linear mixed-effects model fit by REML

Model information:
Number of observations 172
Fixed effects coefficients 2
Random effects coefficients 11
Covariance parameters 2

Formula:
MI ~ 1 + Whisker_stim + (1 | MouseID)

Model fit statistics:

| AIC | BIC | LogLikelihood | Deviance |
|---|---|---|---|
| 519.39 | 531.93 | -255.69 | 511.39 |

Fixed effects coefficients (9 CIs):

| Name | Estimate | SE | tStat | DF | pValue | Lower | Upper | adj. p |
|---|---|---|---|---|---|---|---|---|
| {'(Intercept)' } | 0.69139 | 0.16499 | 4.1905 | 170 | 4.47E-05 | 0.3657 | 1.0171 | 0.0001 |
| {'Whisker_stim_Stm_D1'} | -0.12752 | 0.17424 | -0.73188 | 170 | 0.46525 | -0.47147 | 0.21643 | 0.4653 |

**Figure S3b - % D1 Responsive Cells - D1 barrel**

Model information:

| | |
|---|---|
| Number of observations | 66 |
| Fixed effects coefficients | 6 |
| Random effects coefficients | 11 |
| Covariance parameters | 1 |
| Distribution | Binomial |
| Link | Logit |
| FitMethod | MPL |

**ANOVA marginal tests: DFMethod = 'residual'**

| Term | FStat | DF1 | DF2 | pValue |
|---|---|---|---|---|
| {'(Intercept)' | 24.57 | 1 | 60 | 6.19E-06 |
| {'Timepoint' | 1.2445 | 5 | 60 | 0.29987 |

Formula:
NumRespCells ~ 1 + Timepoint + (1 | MouseID)

Model fit statistics:

| AIC | BIC | LogLikelihood | Deviance |
|---|---|---|---|
| 181.71 | 197.04 | -83.854 | 167.71 |

Fixed effects coefficie (95% CIs):

| Name | Estimate | SE | tStat | DF | pValue | Lower | Upper | adj p |
|---|---|---|---|---|---|---|---|---|
| {'(Intercept)' } | 1.1722 | 0.23649 | 4.9568 | 60 | 6.19E-06 | 0.69918 | 1.6453 | 0 |
| {'Timepoint_Day3' } | 0.56352 | 0.26885 | 2.0961 | 60 | 0.040301 | 0.025751 | 1.1013 | 0.1209 |
| {'Timepoint_Day6' } | 0.36686 | 0.25961 | 1.4131 | 60 | 0.16279 | -0.15244 | 0.88616 | 0.3256 |
| {'Timepoint_Day12'} | 0.030574 | 0.24729 | 0.12364 | 60 | 0.90202 | -0.46408 | 0.52523 | 0.902 |
| {'Timepoint_Day18'} | 0.15781 | 0.25147 | 0.62755 | 60 | 0.53268 | -0.34521 | 0.66084 | 0.6392 |
| {'Timepoint_Day46'} | 0.22479 | 0.2539 | 0.88534 | 60 | 0.37951 | -0.28309 | 0.73268 | 0.5693 |

**Figure S3b - % D1 Responsive Cells - C1 barrel**

Model information:

| | |
|---|---|
| Number of observations | 66 |
| Fixed effects coefficients | 6 |
| Random effects coefficients | 11 |
| Covariance parameters | 1 |
| Distribution | Binomial |
| Link | Logit |
| FitMethod | MPL |

**ANOVA marginal tests: DFMethod = 'residual'**

| Term | FStat | DF1 | DF2 | pValue |
|---|---|---|---|---|
| {'(Intercept)' | 0.079956 | 1 | 60 | 0.77833 |
| {'Timepoint' | 10.073 | 5 | 60 | 4.92E-07 |

Formula:
NumRespCells ~ 1 + Timepoint + (1 | MouseID)

Model fit statistics:

| AIC | BIC | LogLikelihood | Deviance |
|---|---|---|---|
| 162.19 | 177.52 | -74.094 | 148.19 |

Fixed effects coefficients (95% CIs):

| Name | Estimate | SE | tStat | DF | pValue | Lower | Upper | adj p |
|---|---|---|---|---|---|---|---|---|
| {'(Intercept)' } | 0.049793 | 0.17609 | 0.28277 | 60 | 0.77833 | -0.30245 | 0.40203 | 0.9218 |
| {'Timepoint_Day3' } | -0.019458 | 0.19727 | -0.098636 | 60 | 0.92176 | -0.41407 | 0.37515 | 0.9218 |
| {'Timepoint_Day6' } | 0.48 | 0.201 | 2.3881 | 60 | 0.020102 | 0.077942 | 0.88206 | 0.0402 |
| {'Timepoint_Day12'} | 0.80995 | 0.20694 | 3.914 | 60 | 0.00023443 | 0.39602 | 1.2239 | 0.0014 |
| {'Timepoint_Day18'} | 0.60778 | 0.20295 | 2.9947 | 60 | 0.0039869 | 0.20182 | 1.0137 | 0.012 |
| {'Timepoint_Day46'} | -0.41089 | 0.1985 | -2.0699 | 60 | 0.042772 | -0.80795 | -0.013822 | 0.0642 |

**Figure S3c - MI of D1 responsive Cells - D1 barrel**

Model information:

| | |
|---|---|
| Number of observations | 904 |
| Fixed effects coefficients | 6 |
| Random effects coefficients | 1092 |
| Covariance parameters | 22 |

**ANOVA marginal tests: DFMethod = 'residual'**

| Term | FStat | DF1 | DF2 | pValue |
|---|---|---|---|---|
| {'(Intercept)' | 127.47 | 1 | 898 | 9.78E-28 |
| {'Timepoint' | 2.315 | 5 | 898 | 0.041989 |

Formula:
MI ~ 1 + Timepoint + (1 + Timepoint | MouseID:Index)

Model fit statistics:

| AIC | BIC | LogLikelihood | Deviance |
|---|---|---|---|
| 5784.9 | 5919.3 | -2864.5 | 5728.9 |

Fixed effects coefficie (95% CIs):

| Name | Estimate | SE | tStat | DF | pValue | Lower | Upper | adj p |
|---|---|---|---|---|---|---|---|---|
| {'(Intercept)' } | 5.8546 | 0.51854 | 11.29 | 898 | 9.78E-28 | 4.8369 | 6.8723 | 0 |
| {'Timepoint_Day3' } | 0.7804 | 0.44701 | 1.7459 | 898 | 0.081179 | -0.096892 | 1.6577 | 0.1218 |
| {'Timepoint_Day6' } | 1.2951 | 0.60766 | 2.1312 | 898 | 0.03334 | 0.10247 | 2.4877 | 0.0667 |
| {'Timepoint_Day12'} | 0.46071 | 0.5678 | 0.8114 | 898 | 0.41735 | -0.65366 | 1.5751 | 0.4174 |
| {'Timepoint_Day18'} | 1.2445 | 0.4754 | 2.6177 | 898 | 0.0090011 | 0.31143 | 2.1775 | 0.027 |
| {'Timepoint_Day46'} | 0.71498 | 0.81452 | 0.8778 | 898 | 0.38029 | -0.88359 | 2.3136 | 0.4174 |

**Figure S3c - MI of D1 responsive Cells - C1 barrel**

Model information:

| | |
|---|---|
| Number of observations | 725 |
| Fixed effects coefficients | 6 |
| Random effects coefficients | 1122 |
| Covariance parameters | 22 |

**ANOVA marginal tests: DFMethod = 'residual'**

| Term | FStat | DF1 | DF2 | pValue |
|---|---|---|---|---|
| {'(Intercept)' | 42.569 | 1 | 719 | 1.29E-10 |
| {'Timepoint' | 2.5187 | 5 | 719 | 0.028426 |

Formula:
MI ~ 1 + Timepoint + (1 + Timepoint | MouseID:Index)

Model fit statistics:

| AIC | BIC | LogLikelihood | Deviance |
|---|---|---|---|
| 3316.3 | 3444.5 | -1630.1 | 3260.3 |

Fixed effects coefficients (95% CIs):

| Name | Estimate | SE | tStat | DF | pValue | Lower | Upper | adj p |
|---|---|---|---|---|---|---|---|---|
| {'(Intercept)' } | 2.7505 | 0.42157 | 6.5245 | 719 | 1.29E-10 | 1.9229 | 3.5782 | 0 |
| {'Timepoint_Day3' } | -0.24798 | 0.40753 | -0.60848 | 719 | 0.54306 | -1.0481 | 0.55212 | 0.7535 |
| {'Timepoint_Day6' } | 0.0071336 | 0.38221 | 0.018664 | 719 | 0.98511 | -0.74324 | 0.75751 | 0.9851 |
| {'Timepoint_Day12'} | 0.21083 | 0.32916 | 0.6405 | 719 | 0.52205 | -0.43541 | 0.85706 | 0.7535 |
| {'Timepoint_Day18'} | 0.18178 | 0.37492 | 0.48484 | 719 | 0.62794 | -0.5543 | 0.91785 | 0.7535 |
| {'Timepoint_Day46'} | -0.53224 | 0.42979 | -1.2384 | 719 | 0.21599 | -1.376 | 0.31156 | 0.648 |

**Figure S3d - MI of all cells - D1 barrel**

Model information:

| | |
|---|---|
| Number of observations | 1134 |
| Fixed effects coefficients | 6 |
| Random effects coefficients | 1134 |
| Covariance parameters | 22 |

**ANOVA marginal tests: DFMethod = 'residual'**

| Term | FStat | DF1 | DF2 | pValue |
|---|---|---|---|---|
| {'(Intercept)' | 125.16 | 1 | 1128 | 1.26E-27 |
| {'Timepoint' | 4.4118 | 5 | 1128 | 0.00055307 |

Formula:
MI ~ 1 + Timepoint + (1 + Timepoint | MouseID:Index)

Model fit statistics:

| AIC | BIC | LogLikelihood | Deviance |
|---|---|---|---|
| 6975.6 | 7116.3 | -3459.8 | 6919.6 |

Fixed effects coefficie (95% CIs):

| Name | Estimate | SE | tStat | DF | pValue | Lower | Upper | adj p |
|---|---|---|---|---|---|---|---|---|
| {'(Intercept)' } | 5.2434 | 0.46868 | 11.188 | 1128 | 1.26E-27 | 4.3238 | 6.163 | 0 |
| {'Timepoint_Day3' } | 0.82622 | 0.34762 | 2.3768 | 1128 | 0.01763 | 0.14416 | 1.5083 | 0.0264 |
| {'Timepoint_Day6' } | 1.4409 | 0.49389 | 2.9175 | 1128 | 0.0035983 | 0.47188 | 2.41 | 0.0072 |
| {'Timepoint_Day12'} | 0.53355 | 0.44688 | 1.194 | 1128 | 0.23275 | -0.34325 | 1.4104 | 0.2604 |
| {'Timepoint_Day18'} | 1.2965 | 0.37273 | 3.4783 | 1128 | 0.00052388 | 0.56514 | 2.0278 | 0.0016 |
| {'Timepoint_Day46'} | 0.75155 | 0.66746 | 1.126 | 1128 | 0.26041 | -0.55805 | 2.0611 | 0.2604 |

**Figure S3e - MI of all cells - C1 barrel**

Model information:

| | |
|---|---|
| Number of observations | 1260 |
| Fixed effects coefficients | 6 |
| Random effects coefficients | 1260 |
| Covariance parameters | 22 |

**ANOVA marginal tests: DFMethod = 'residual'**

| Term | FStat | DF1 | DF2 | pValue |
|---|---|---|---|---|
| {'(Intercept)' | 58.534 | 1 | 1254 | 3.97E-14 |
| {'Timepoint' | 13.722 | 5 | 1254 | 4.56E-13 |

Formula:
MI ~ 1 + Timepoint + (1 + Timepoint | MouseID:Index)

Model fit statistics:

| AIC | BIC | LogLikelihood | Deviance |
|---|---|---|---|
| 5168.6 | 5312.4 | -2556.3 | 5112.6 |

Fixed effects coefficie (95% CIs):

| Name | Estimate | SE | tStat | DF | pValue | Lower | Upper | adj p |
|---|---|---|---|---|---|---|---|---|
| {'(Intercept)' } | 2.2303 | 0.29151 | 7.6507 | 1254 | 3.97E-14 | 1.6584 | 2.8021 | 0 |
| {'Timepoint_Day3' } | -0.3227 | 0.23115 | -1.3961 | 1254 | 0.16293 | -0.77618 | 0.13078 | 0.3259 |
| {'Timepoint_Day6' } | -0.045041 | 0.21962 | -0.20508 | 1254 | 0.83754 | -0.47591 | 0.38583 | 0.8375 |
| {'Timepoint_Day12'} | 0.16416 | 0.19407 | 0.84589 | 1254 | 0.39777 | -0.21658 | 0.54491 | 0.4773 |
| {'Timepoint_Day18'} | 0.20212 | 0.21334 | 0.9474 | 1254 | 0.34362 | -0.21643 | 0.62067 | 0.4773 |
| {'Timepoint_Day46'} | -0.83659 | 0.2427 | -3.447 | 1254 | 0.00058572 | -1.3127 | -0.36044 | 0.0018 |

**Figure S3f - % Co-Modulated Cells/Stim- D1 barrel**

Linear mixed-effects model fit by REML

Model information:

| | |
|---|---|
| Number of observations | 66 |
| Fixed effects coefficients | 6 |
| Random effects coefficients | 11 |
| Covariance parameters | 2 |

**ANOVA marginal tests: DFMethod = 'residual'**

| Term | FStat | DF1 | DF2 | pValue |
|---|---|---|---|---|
| {'(Intercept)' | 477.59 | 1 | 60 | 2.93E-30 |
| {'Timepoint' | 1.2034 | 5 | 60 | 0.3186 |

Formula:
stim_PrctCellMod ~ 1 + Timepoint + (1 | MouseID)

Model fit statistics:

| AIC | BIC | LogLikelihood | Deviance |
|---|---|---|---|
| -105.54 | -88.783 | 60.769 | -121.54 |

Fixed effects coefficie (95% CIs):

| Name | Estimate | SE | tStat | DF | pValue | Lower | Upper | adj p |
|---|---|---|---|---|---|---|---|---|
| {'(Intercept)' } | 0.75466 | 0.034532 | 21.854 | 60 | 2.93E-30 | 0.68559 | 0.82374 | 0 |
| {'Timepoint_Day3' } | 0.03312 | 0.026461 | 1.2516 | 60 | 0.21556 | -0.01981 | 0.086049 | 0.531 |
| {'Timepoint_Day6' } | 0.02974 | 0.026461 | 1.1239 | 60 | 0.26552 | -0.02319 | 0.08267 | 0.531 |
| {'Timepoint_Day12'} | 0.011064 | 0.026461 | 0.41811 | 60 | 0.67736 | -0.041866 | 0.063993 | 0.6774 |
| {'Timepoint_Day18'} | 0.0121 | 0.026461 | 0.45726 | 60 | 0.64913 | -0.04083 | 0.065029 | 0.6774 |
| {'Timepoint_Day46'} | -0.022839 | 0.026461 | -0.86312 | 60 | 0.39151 | -0.075769 | 0.030091 | 0.5873 |

**Figure S3f - % Co-Modulated Cells/Stim- C1 barrel**

Linear mixed-effects model fit by REML

Model information:

| | |
|---|---|
| Number of observations | 66 |
| Fixed effects coefficients | 6 |
| Random effects coefficients | 11 |
| Covariance parameters | 2 |

**ANOVA marginal tests: DFMethod = 'residual'**

| Term | FStat | DF1 | DF2 | pValue |
|---|---|---|---|---|
| {'(Intercept)' | 284.51 | 1 | 60 | 1.90E-24 |
| {'Timepoint' | 4.1318 | 5 | 60 | 0.0027268 |

Formula:
stim_PrctCellMod ~ 1 + Timepoint + (1 | MouseID)

Model fit statistics:

| AIC | BIC | LogLikelihood | Deviance |
|---|---|---|---|
| -79.35 | -62.595 | 47.675 | -95.35 |

Fixed effects coefficients (95% CIs):

| Name | Estimate | SE | tStat | DF | pValue | Lower | Upper | adj p |
|---|---|---|---|---|---|---|---|---|
| {'(Intercept)' } | 0.57743 | 0.034233 | 16.868 | 60 | 1.90E-24 | 0.50896 | 0.64591 | 0 |
| {'Timepoint_Day3' } | -0.060115 | 0.035548 | -1.6911 | 60 | 0.096007 | -0.13122 | 0.010991 | 0.192 |
| {'Timepoint_Day6' } | -0.024673 | 0.035548 | -0.69408 | 60 | 0.49031 | -0.095779 | 0.046433 | 0.4903 |
| {'Timepoint_Day12'} | 0.033445 | 0.035548 | 0.94084 | 60 | 0.35056 | -0.037661 | 0.10455 | 0.4344 |
| {'Timepoint_Day18'} | 0.032652 | 0.035548 | 0.91853 | 60 | 0.36202 | -0.038454 | 0.10376 | 0.4344 |
| {'Timepoint_Day46'} | -0.093508 | 0.035548 | -2.6305 | 60 | 0.010818 | -0.16461 | -0.022402 | 0.0325 |

**Figure S3g - Xcorr of all cells in D1 barrel**

Linear mixed-effects model fit by REML

Model information:

| | |
|---|---|
| Number of observations | 1128 |
| Fixed effects coefficients | 6 |
| Random effects coefficients | 1128 |
| Covariance parameters | 22 |

**ANOVA marginal tests: DFMethod = 'residual'**

| Term | FStat | DF1 | DF2 | pValue |
|---|---|---|---|---|
| {'(Intercept)' | 4774.8 | 1 | 1122 | 0.00E+00 |
| {'Timepoint' | 22.15 | 5 | 1122 | 3.22E-21 |

Formula:
Rreal_avg ~ 1 + Timepoint + (1 + Timepoint | MouseID:Index)

Model fit statistics:

| AIC | BIC | LogLikelihood | Deviance |
|---|---|---|---|
| -2001.2 | -1860.5 | 1028.6 | -2057.2 |

Fixed effects coefficie (95% CIs):

| Name | Estimate | SE | tStat | DF | pValue | Lower | Upper | adj p |
|---|---|---|---|---|---|---|---|---|
| {'(Intercept)' } | 0.69036 | 0.0099908 | 69.1 | 1122 | 0.00E+00 | 0.67076 | 0.70996 | 0 |
| {'Timepoint_Day3' } | 0.035127 | 0.0070026 | 5.0163 | 1122 | 6.12E-07 | 0.021387 | 0.048866 | 0 |
| {'Timepoint_Day6' } | 0.016546 | 0.0080369 | 2.0587 | 1122 | 0.03975 | 0.00077683 | 0.032315 | 0.0397 |
| {'Timepoint_Day12'} | 0.028355 | 0.0072836 | 3.893 | 1122 | 0.00010486 | 0.014064 | 0.042646 | 0.0002 |
| {'Timepoint_Day18'} | 0.059048 | 0.0076847 | 7.6839 | 1122 | 3.35E-14 | 0.04397 | 0.074126 | 0 |
| {'Timepoint_Day46'} | -0.019731 | 0.0094856 | -2.0801 | 1122 | 0.037744 | -0.038342 | -0.0011194 | 0.0397 |

**Figure S3h - Xcorr of all cells in C1 barrel**

Linear mixed-effects model fit by REML

Model information:

| | |
|---|---|
| Number of observations | 1260 |
| Fixed effects coefficients | 6 |
| Random effects coefficients | 1260 |
| Covariance parameters | 22 |

**ANOVA marginal tests: DFMethod = 'residual'**

| Term | FStat | DF1 | DF2 | pValue |
|---|---|---|---|---|
| {'(Intercept)' | 6042.9 | 1 | 1254 | 0.00E+00 |
| {'Timepoint' | 22.181 | 5 | 1254 | 2.36E-21 |

Formula:
Rreal_avg ~ 1 + Timepoint + (1 + Timepoint | MouseID:Index)

Model fit statistics:

| AIC | BIC | LogLikelihood | Deviance |
|---|---|---|---|
| -1715.7 | -1572 | 885.87 | -1771.7 |

Fixed effects coefficie (95% CIs):

| Name | Estimate | SE | tStat | DF | pValue | Lower | Upper | adj p |
|---|---|---|---|---|---|---|---|---|
| {'(Intercept)' } | 0.65152 | 0.0083813 | 77.736 | 1254 | 0.00E+00 | 0.63508 | 0.66797 | 0 |
| {'Timepoint_Day3' } | -0.089843 | 0.01195 | -7.5182 | 1254 | 1.05E-13 | -0.11329 | -0.066398 | 0 |
| {'Timepoint_Day6' } | -0.085795 | 0.010232 | -8.3853 | 1254 | 1.34E-16 | -0.10587 | -0.065722 | 0 |
| {'Timepoint_Day12'} | -0.048672 | 0.0094939 | -5.1267 | 1254 | 3.41E-07 | -0.067298 | -0.030047 | 0 |
| {'Timepoint_Day18'} | -0.02076 | 0.0096714 | -2.1466 | 1254 | 0.03202 | -0.039734 | -0.0017863 | 0.032 |
| {'Timepoint_Day46'} | -0.065152 | 0.010436 | -6.2427 | 1254 | 5.87E-10 | -0.085627 | -0.044677 | 0 |

**Figure S4a - % D1 whisker responsive cells - D1 barrel**

Model information:

| | |
|---|---|
| Number of observations | 15 |
| Fixed effects coefficients | 3 |
| Random effects coefficients | 5 |
| Covariance parameters | 1 |
| Distribution | Binomial |
| Link | Logit |
| FitMethod | MPL |

ANOVA marginal tests: DFMethod = 'residual'

| Term | FStat | DF1 | DF2 | pValue |
|---|---|---|---|---|
| {'(Intercept)' | 21.878 | 1 | 12 | 0.00053463 |
| {'Timepoint' | 1.3744 | 2 | 12 | 0.29009 |

Formula:
NumRespCells ~ 1 + Timepoint + (1 | MouseID)

Model fit statistics:

| AIC | BIC | LogLikelihood | Deviance |
|---|---|---|---|
| 41.402 | 44.234 | -16.701 | 33.402 |

| Name | Estimate | SE | tStat | DF | pValue | Lower | Upper | Adj p. |
|---|---|---|---|---|---|---|---|---|
| {'(Intercept)' } | 1.7334 | 0.37059 | 4.6774 | 12 | 0.00053463 | 0.92596 | 2.5409 | 0.0016 |
| {'Timepoint_18'} | -0.14418 | 0.38009 | -0.37934 | 12 | 0.71106 | -0.97232 | 0.68396 | 0.7111 |
| {'Timepoint_46'} | 0.54709 | 0.43394 | 1.2607 | 12 | 0.23136 | -0.39838 | 1.4926 | 0.347 |

**Figure S4b - MI of D1 whisker responsive cells in D1 barrel**

Model information:

| | |
|---|---|
| Number of observations | 280 |
| Fixed effects coefficients | 3 |
| Random effects coefficients | 309 |
| Covariance parameters | 7 |

ANOVA marginal tests: DFMethod = 'residual'

| Term | FStat | DF1 | DF2 | pValue |
|---|---|---|---|---|
| {'(Intercept)' | 92.842 | 1 | 277 | 3.88E-19 |
| {'Timepoint' | 2.957 | 2 | 277 | 0.05362 |

Formula:
MI ~ 1 + Timepoint + (1 + Timepoint | MouseID:index)

Model fit statistics:

| AIC | BIC | LogLikelihood | Deviance |
|---|---|---|---|
| 1942.1 | 1978.4 | -961.03 | 1922.1 |

| Name | Estimate | SE | tStat | DF | pValue | Lower | Upper | Adj p |
|---|---|---|---|---|---|---|---|---|
| {'(Intercept)' } | 7.2338 | 0.75075 | 9.6354 | 277 | 3.88E-19 | 5.7559 | 8.7117 | 0 |
| {'Timepoint_18'} | 1.2298 | 1.0225 | 1.2027 | 277 | 0.23011 | -0.78304 | 3.2426 | 0.2301 |
| {'Timepoint_46'} | 2.2931 | 0.96535 | 2.3754 | 277 | 0.018209 | 0.39277 | 4.1935 | 0.0273 |

**Figure S4c - Whisker Length Pre vs Post Trimming**

Paired t-test p=0.07

**Figure S4a - % D1 whisker responsive cells -C1 barrel**

Model information:

| | |
|---|---|
| Number of observations | 15 |
| Fixed effects coefficients | 3 |
| Random effects coefficients | 5 |
| Covariance parameters | 1 |
| Distribution | Binomial |
| Link | Logit |
| FitMethod | MPL |

ANOVA marginal tests: DFMethod = 'residual'

| Term | FStat | DF1 | DF2 | pValue |
|---|---|---|---|---|
| {'(Intercept)' | 0.61493 | 1 | 12 | 0.44814 |
| {'Timepoint' | 8.2153 | 2 | 12 | 0.0056543 |

Formula:
NumRespCells ~ 1 + Timepoint + (1 | MouseID)

Model fit statistics:

| AIC | BIC | LogLikelihood | Deviance |
|---|---|---|---|
| 46.212 | 49.044 | -19.106 | 38.212 |

| Name | Estimate | SE | tStat | DF | pValue | Lower | Upper | Adj p. |
|---|---|---|---|---|---|---|---|---|
| {'(Intercept)' } | -0.32518 | 0.41468 | -0.78418 | 12 | 0.44814 | -1.2287 | 0.57833 | 0.6722 |
| {'Timepoint_18'} | -1.1316 | 0.31204 | -3.6264 | 12 | 0.0034734 | -1.8115 | -0.45172 | 0.0104 |
| {'Timepoint_46'} | -1.55E-16 | 0.27882 | -5.55E-16 | 12 | 1 | -0.60749 | 0.60749 | 1 |

**Figure S4b - MI of D1 whisker responsive cells in C1 barrel**

**Model information:**

| | |
|---|---|
| Number of observations | 122 |
| Fixed effects coefficients | 3 |
| Random effects coefficients | 192 |
| Covariance parameters | 7 |

ANOVA marginal tests: DFMethod = 'residual'

| Term | FStat | DF1 | DF2 | pValue |
|---|---|---|---|---|
| {'(Intercept)' | 68.858 | 1 | 119 | 1.89E-13 |
| {'Timepoint' | 1.9672 | 2 | 119 | 0.14437 |

Formula:
MI ~ 1 + Timepoint + (1 + Timepoint | MouseID:index)

Model fit statistics:

| AIC | BIC | LogLikelihood | Deviance |
|---|---|---|---|
| 760.03 | 788.07 | -370.02 | 740.03 |

| Name | Estimate | SE | tStat | DF | pValue | Lower | Upper | Adj p |
|---|---|---|---|---|---|---|---|---|
| {'(Intercept)' } | 5.2884 | 0.63731 | 8.298 | 119 | 1.89E-13 | 4.0265 | 6.5503 | 0 |
| {'Timepoint_18'} | -1.0494 | 0.98687 | -1.0634 | 119 | 0.28977 | -3.0035 | 0.90471 | 0.3566 |
| {'Timepoint_46'} | 0.81283 | 0.87829 | 0.92547 | 119 | 0.3566 | -0.92627 | 2.5519 | 0.3566 |

**Figure S5a - AUC of spont. Activity of all cells in D1 barrel**
Linear mixed-effects model fit by ML

Model information:
Number of observations 1134
Fixed effects coefficients 6
Random effects coefficients 22
Covariance parameters 4

Formula:
AUC ~ 1 + Timepoint + (1 + index | MouseID)

ANOVA marginal tests: DFMethod = 'Residual'

| Term | FStat | DF1 | DF2 | pValue |
|---|---|---|---|---|
| {'(Intercept)' | 264.71 | 1 | 1128 | 1.26E-53 |
| {'Timepoint' | 5.851 | 5 | 1128 | 2.40E-05 |

Model fit statistics:

| AIC | BIC | LogLikelihood | Deviance |
|---|---|---|---|
| 16049 | 16100 | -8014.7 | 16029 |

| Name | Estimate | SE | tStat | DF | pValue | Lower | Upper | adj p |
|---|---|---|---|---|---|---|---|---|
| {'(Intercept)' } | 439.16 | 26.992 | 16.27 | 1128 | 1.26E-53 | 386.2 | 492.12 | 0 |
| {'Timepoint_3' } | -34.833 | 28.879 | -1.2062 | 1128 | 0.228 | -91.496 | 21.829 | 0.456 |
| {'Timepoint_6' } | -21.941 | 28.879 | -0.75975 | 1128 | 0.44756 | -78.604 | 34.722 | 0.5371 |
| {'Timepoint_12'} | 14.712 | 28.879 | 0.50944 | 1128 | 0.61055 | -41.951 | 71.375 | 0.6105 |
| {'Timepoint_18'} | 22.616 | 28.879 | 0.78311 | 1128 | 0.43372 | -34.047 | 79.278 | 0.5371 |
| {'Timepoint_46'} | -112.62 | 28.879 | -3.8998 | 1128 | 0.00010196 | -169.28 | -55.959 | 0.0003 |

**Figure S5b- Peak Freq. of spont. Activity of all cells in D1 barrel**
Linear mixed-effects model fit by ML

Model information:
Number of observations 1134
Fixed effects coefficients 6
Random effects coefficients 22
Covariance parameters 4

Formula:
PeakFreq ~ 1 + Timepoint + (1 + index | MouseID)

ANOVA marginal tests: DFMethod = 'Residual'

| Term | FStat | DF1 | DF2 | pValue |
|---|---|---|---|---|
| {'(Intercept)' | 681.5 | 1 | 1128 | 6.70E-118 |
| {'Timepoint' | 3.5891 | 5 | 1128 | 0.0031702 |

Model fit statistics:

| AIC | BIC | LogLikelihood | Deviance |
|---|---|---|---|
| -1639.6 | -1589.3 | 829.81 | -1659.6 |

| Name | Estimate | SE | tStat | DF | pValue | Lower | Upper | adj p |
|---|---|---|---|---|---|---|---|---|
| {'(Intercept)' } | 0.28361 | 0.010864 | 26.106 | 1128 | 6.70E-118 | 0.2623 | 0.30493 | 0 |
| {'Timepoint_3' } | -0.0051573 | 0.011848 | -0.43528 | 1128 | 0.66344 | -0.028404 | 0.01809 | 0.7961 |
| {'Timepoint_6' } | -0.0027199 | 0.011848 | -0.22957 | 1128 | 0.81847 | -0.025967 | 0.020527 | 0.8185 |
| {'Timepoint_12'} | 0.014483 | 0.011848 | 1.2224 | 1128 | 0.22182 | -0.0087641 | 0.03773 | 0.4436 |
| {'Timepoint_18'} | 0.031368 | 0.011848 | 2.6475 | 1128 | 0.0082226 | 0.0081208 | 0.054615 | 0.0247 |
| {'Timepoint_46'} | -0.01201 | 0.011848 | -1.0137 | 1128 | 0.31096 | -0.035257 | 0.011237 | 0.4664 |

**Figure S5c- Peak Amp. of spont. Activity of all cells in D1 barrel**
Linear mixed-effects model fit by ML

Model information:
Number of observations 1133
Fixed effects coefficients 6
Random effects coefficients 22
Covariance parameters 4

Formula:
PeaksAmp_mean ~ 1 + Timepoint + (1 + index | MouseI

ANOVA marginal tests: DFMethod = 'Residual'

| Term | FStat | DF1 | DF2 | pValue |
|---|---|---|---|---|
| {'(Intercept)' | 903.99 | 1 | 1127 | 2.62E-146 |
| {'Timepoint' | 4.3002 | 5 | 1127 | 0.00070255 |

Model fit statistics:

| AIC | BIC | LogLikelihood | Deviance |
|---|---|---|---|
| 5960.7 | 6011 | -2970.3 | 5940.7 |

| Name | Estimate | SE | tStat | DF | pValue | Lower | Upper | adj p |
|---|---|---|---|---|---|---|---|---|
| {'(Intercept)' } | 12.016 | 0.39965 | 30.066 | 1127 | 2.62E-146 | 11.232 | 12.8 | 0 |
| {'Timepoint_3' } | -0.64696 | 0.33139 | -1.9522 | 1127 | 0.051158 | -1.2972 | 0.0032597 | 0.1023 |
| {'Timepoint_6' } | -0.2416 | 0.33184 | -0.72807 | 1127 | 0.46672 | -0.8927 | 0.40949 | 0.5601 |
| {'Timepoint_12'} | 0.15963 | 0.33139 | 0.48169 | 1127 | 0.63012 | -0.49059 | 0.80985 | 0.6301 |
| {'Timepoint_18'} | 0.59023 | 0.33139 | 1.7811 | 1127 | 0.075173 | -0.059988 | 1.2404 | 0.1128 |
| {'Timepoint_46'} | -0.66062 | 0.33139 | -1.9935 | 1127 | 0.046451 | -1.3108 | -0.010404 | 0.1023 |

**Figure S5d- Xcorr of spont. Activity of all cells in D1 barrel**
Linear mixed-effects model fit by REML

Model information:
Number of observations 1128
Fixed effects coefficients 6
Random effects coefficients 1128
Covariance parameters 22

Formula:
Rreal_avg ~ 1 + Timepoint + (1 + Timepoint | MouseID:ir

ANOVA marginal tests: DFMethod = 'Residual'

| Term | FStat | DF1 | DF2 | pValue |
|---|---|---|---|---|
| {'(Intercept)' | 3968.6 | 1 | 1122 | 0.00E+00 |
| {'Timepoint' | 9.5091 | 5 | 1122 | 6.77E-09 |

Model fit statistics:

| AIC | BIC | LogLikelihood | Deviance |
|---|---|---|---|
| -1457.9 | -1317.3 | 756.95 | -1513.9 |

| Name | Estimate | SE | tStat | DF | pValue | Lower | Upper | adj p |
|---|---|---|---|---|---|---|---|---|
| {'(Intercept)' } | 0.68069 | 0.010805 | 62.997 | 1122 | 0.00E+00 | 0.65949 | 0.70189 | 0 |
| {'Timepoint_3' } | -0.03033 | 0.011918 | -2.545 | 1122 | 0.011062 | -0.053714 | -0.0069467 | 0.0166 |
| {'Timepoint_6' } | -0.039317 | 0.0088549 | -4.4401 | 1122 | 9.88E-06 | -0.056691 | -0.021943 | 0 |
| {'Timepoint_12'} | -0.0016508 | 0.0085674 | -0.19268 | 1122 | 0.84725 | -0.018461 | 0.015159 | 0.8472 |
| {'Timepoint_18'} | 0.0083906 | 0.0096979 | 0.8652 | 1122 | 0.38711 | -0.010637 | 0.027419 | 0.4645 |
| {'Timepoint_46'} | -0.066063 | 0.014944 | -4.4208 | 1122 | 1.08E-05 | -0.095384 | -0.036742 | 0 |

**Figure S5e - Xcorr of spont. Activity of all cells in C1 barrel**
Linear mixed-effects model fit by REML

Model information:
Number of observations 1260
Fixed effects coefficients 6
Random effects coefficients 1260
Covariance parameters 22

Formula:
Rreal_avg ~ 1 + Timepoint + (1 + Timepoint | MouseID:ir

ANOVA marginal tests: DFMethod = 'Residual'

| Term | FStat | DF1 | DF2 | pValue |
|---|---|---|---|---|
| {'(Intercept)' | 6981.1 | 1 | 1254 | 0.00E+00 |
| {'Timepoint' | 43.397 | 5 | 1254 | 2.37E-41 |

Model fit statistics:

| AIC | BIC | LogLikelihood | Deviance |
|---|---|---|---|
| -1506.2 | -1362.4 | 781.08 | -1562.2 |

| Name | Estimate | SE | tStat | DF | pValue | Lower | Upper | adj p |
|---|---|---|---|---|---|---|---|---|
| {'(Intercept)' } | 0.67318 | 0.008057 | 83.553 | 1254 | 0.00E+00 | 0.65737 | 0.68899 | 0 |
| {'Timepoint_3' } | -0.14473 | 0.012657 | -11.435 | 1254 | 7.13E-29 | -0.16956 | -0.1199 | 0 |
| {'Timepoint_6' } | -0.14398 | 0.011655 | -12.353 | 1254 | 3.66E-33 | -0.16684 | -0.12111 | 0 |
| {'Timepoint_12'} | -0.087922 | 0.010054 | -8.745 | 1254 | 7.04E-18 | -0.10765 | -0.068198 | 0 |
| {'Timepoint_18'} | -0.059592 | 0.0099782 | -5.9722 | 1254 | 3.04E-09 | -0.079168 | -0.040016 | 0.3044 |
| {'Timepoint_46'} | -0.084505 | 0.009349 | -9.0389 | 1254 | 5.85E-19 | -0.10285 | -0.066163 | 0 |

**Figure S5a - AUC of spont. Activity of all cells in C1 barrel**
Linear mixed-effects model fit by ML

Model information:
Number of observations 1260
Fixed effects coefficients 6
Random effects coefficients 22
Covariance parameters 4

Formula:
AUC ~ 1 + Timepoint + (1 + index | MouseID)

ANOVA marginal tests: DFMethod = 'Residual'

| Term | FStat | DF1 | DF2 | pValue |
|---|---|---|---|---|
| {'(Intercept)' | 1.9289 | 1 | 1254 | 0.16512 |
| {'Timepoint' | 4.6823 | 5 | 1254 | 0.00030597 |

Model fit statistics:

| AIC | BIC | LogLikelihood | Deviance |
|---|---|---|---|
| 17889 | 17941 | -8934.6 | 17869 |

| Name | Estimate | SE | tStat | DF | pValue | Lower | Upper | adj p |
|---|---|---|---|---|---|---|---|---|
| {'(Intercept)' } | 494.69 | 356.18 | 1.3889 | 1254 | 0.16512 | -204.09 | 1193.5 | 0.1651 |
| {'Timepoint_3' } | -84.136 | 27.233 | -3.0895 | 1254 | 0.0020493 | -137.56 | -30.708 | 0.0041 |
| {'Timepoint_6' } | -93.949 | 27.233 | -3.4498 | 1254 | 0.00057965 | -147.38 | -40.521 | 0.0017 |
| {'Timepoint_12'} | -73.437 | 27.233 | -2.6966 | 1254 | 0.0070988 | -126.86 | -20.01 | 0.0106 |
| {'Timepoint_18'} | -70.631 | 27.233 | -2.5936 | 1254 | 0.0096089 | -124.06 | -17.204 | 0.0115 |
| {'Timepoint_46'} | -125.92 | 27.233 | -4.6238 | 1254 | 4.16E-06 | -179.35 | -72.494 | 0 |

**Figure S5b- Peak Freq. of spont. Activity of all cells inC1 barrel**
Linear mixed-effects model fit by ML

Model information:
Number of observations 1260
Fixed effects coefficients 6
Random effects coefficients 22
Covariance parameters 4

Formula:
PeakFreq ~ 1 + Timepoint + (1 + index | MouseID)

ANOVA marginal tests: DFMethod = 'Residual'

| Term | FStat | DF1 | DF2 | pValue |
|---|---|---|---|---|
| {'(Intercept)' | 904.1 | 1 | 1254 | 5.13E-150 |
| {'Timepoint' | 2.1463 | 5 | 1254 | 0.057699 |

Model fit statistics:

| AIC | BIC | LogLikelihood | Deviance |
|---|---|---|---|
| -1784.5 | -1733.1 | 902.27 | -1804.5 |

| Name | Estimate | SE | tStat | DF | pValue | Lower | Upper | adj p |
|---|---|---|---|---|---|---|---|---|
| {'(Intercept)' } | 0.27943 | 0.0092931 | 30.068 | 1254 | 5.13E-150 | 0.2612 | 0.29766 | 0 |
| {'Timepoint_3' } | -0.016309 | 0.011462 | -1.4229 | 1254 | 0.15502 | -0.038796 | 0.006178 | 0.2875 |
| {'Timepoint_6' } | -0.02289 | 0.011462 | -1.997 | 1254 | 0.046041 | -0.045377 | -0.0004028 | 0.1381 |
| {'Timepoint_12'} | -0.014974 | 0.011462 | -1.3064 | 1254 | 0.19167 | -0.037461 | 0.0075133 | 0.2875 |
| {'Timepoint_18'} | 0.0079161 | 0.011462 | 0.69063 | 1254 | 0.48993 | -0.014571 | 0.030403 | 0.5879 |
| {'Timepoint_46'} | -0.0010491 | 0.011462 | -0.091529 | 1254 | 0.92709 | -0.023536 | 0.021438 | 0.9271 |

**Figure S5c- Peak Amp. of spont. Activity of all cells in C1 barrel**
Linear mixed-effects model fit by ML

Model information:
Number of observations 1259
Fixed effects coefficients 6
Random effects coefficients 22
Covariance parameters 4

Formula:
PeaksAmp_mean ~ 1 + Timepoint + (1 + index | MouseID

ANOVA marginal tests: DFMethod = 'Residual'

| Term | FStat | DF1 | DF2 | pValue |
|---|---|---|---|---|
| {'(Intercept)' | 976.85 | 1 | 1253 | 5.05E-159 |
| {'Timepoint' | 5.6163 | 5 | 1253 | 3.97E-05 |

Model fit statistics:

| AIC | BIC | LogLikelihood | Deviance |
|---|---|---|---|
| 6524.5 | 6575.9 | -3252.3 | 6504.5 |

| Name | Estimate | SE | tStat | DF | pValue | Lower | Upper | adj p |
|---|---|---|---|---|---|---|---|---|
| {'(Intercept)' } | 8.6872 | 0.27795 | 31.255 | 1253 | 5.05E-159 | 8.1419 | 9.2325 | 0 |
| {'Timepoint_3' } | -1.2586 | 0.30912 | -4.0714 | 1253 | 4.97E-05 | -1.865 | -0.6521 | 0.0001 |
| {'Timepoint_6' } | -1.3005 | 0.30912 | -4.207 | 1253 | 2.77E-05 | -1.9069 | -0.69402 | 0.0001 |
| {'Timepoint_12'} | -1.1705 | 0.3095 | -3.782 | 1253 | 0.0001629 | -1.7777 | -0.56333 | 0.0002 |
| {'Timepoint_18'} | -0.65736 | 0.30912 | -2.1265 | 1253 | 0.033655 | -1.2638 | -0.050901 | 0.0337 |
| {'Timepoint_46'} | -1.2464 | 0.30912 | -4.0322 | 1253 | 5.86E-05 | -1.8529 | -0.63998 | 0.0001 |

**Figure S6a - MI to D1 whisker By Switch category - D1 barrel**

| Kruskal Wallis | 2.63E-22 | | | | |
|---|---|---|---|---|---|
| Group A | Group B | Lower Limit | A-B | Upper Limit | P-value |
| NR to NR | NR to R | -62.027391 | -18.657143 | 24.7131051 | 0.68635641 |
| NR to NR | R to NR | -112.76191 | -64.184416 | -15.606922 | 3.84E-03 |
| NR to NR | R to R | -124.61528 | -97.808705 | -71.002129 | 3.77E-09 |
| NR to R | R to NR | -101.31405 | -45.527273 | 10.2595024 | 0.15415894 |
| NR to R | R to R | -117.50497 | -79.151563 | -40.798155 | 6.88E-07 |
| R to NR | R to R | -77.780585 | -33.62429 | 10.5320054 | 0.20478454 |

**Figure S6b - MI to C1 whisker By Switch category - D1 barrel**

| Kruskal Wallis | 3.85E-11 | | | | |
|---|---|---|---|---|---|
| Group A | Group B | Lower Limit | A-B | Upper Limit | P-value |
| NR to NR | NR to R | -62.513105 | -19.142857 | 24.2273909 | 0.66849815 |
| NR to NR | R to NR | -74.683987 | -26.106494 | 22.4710004 | 0.51150973 |
| NR to NR | R to R | -95.768183 | -68.961607 | -42.155031 | 3.99E-09 |
| NR to R | R to NR | -62.750412 | -6.9636364 | 48.8231388 | 0.98860145 |
| NR to R | R to R | -88.172158 | -49.81875 | -11.465342 | 0.00469939 |
| R to NR | R to R | -87.011409 | -42.855114 | 1.30118153 | 0.0609308 |

**Figure S6c - SI By Switch category - D1 barrel**

| Kruskal Wallis | 0.0411 | | | | |
|---|---|---|---|---|---|
| Group A | Group B | Lower Limit | A-B | Upper Limit | P-value |
| NR to NR | NR to R | -22.812605 | 20.5333333 | 63.8792718 | 0.61602051 |
| NR to NR | R to NR | -2.0684475 | 46.4818182 | 95.0320838 | 6.64E-02 |
| NR to NR | R to R | -1.7415509 | 25.05 | 51.8415509 | 7.67E-02 |
| NR to R | R to NR | -29.807021 | 25.9484848 | 81.7039908 | 0.62961154 |
| NR to R | R to R | -33.815244 | 4.51666667 | 42.848577 | 9.90E-01 |
| R to NR | R to R | -65.563363 | -21.431818 | 22.6997269 | 0.59644432 |

**Figure S6d - MI to D1 whisker By Switch category - C1 barrel**

| Kruskal Wallis | 1.19E-29 | | | | |
|---|---|---|---|---|---|
| Group A | Group B | Lower Limit | A-B | Upper Limit | P-value |
| NR to NR | NR to R | -56.545219 | -26.35278 | 3.83965968 | 0.11197664 |
| NR to NR | R to NR | -138.70475 | -91.084367 | -43.463989 | 5.32E-06 |
| NR to NR | R to R | -135.95479 | -110.07038 | -84.185977 | 3.77E-09 |
| NR to R | R to NR | -113.6514 | -64.731588 | -15.811779 | 0.00377146 |
| NR to R | R to R | -111.92132 | -83.717602 | -55.513885 | 3.77E-09 |
| R to NR | R to R | -65.370997 | -18.986014 | 27.3989691 | 0.71899349 |

**Figure S6e - MI to C1 whisker By Switch category - C1 barrel**

| Kruskal Wallis | 2.27E-08 | | | | |
|---|---|---|---|---|---|
| Group A | Group B | Lower Limit | A-B | Upper Limit | P-value |
| NR to NR | NR to R | -61.699989 | -31.50755 | -1.3151104 | 0.03688871 |
| NR to NR | R to NR | -124.59929 | -76.978908 | -29.35853 | 0.00019234 |
| NR to NR | R to R | -82.9009 | -57.016496 | -31.132091 | 9.46E-08 |
| NR to R | R to NR | -94.391167 | -45.471358 | 3.44844999 | 0.07936883 |
| NR to R | R to R | -53.712663 | -25.508946 | 2.69477106 | 0.09269166 |
| R to NR | R to R | -26.42257 | 19.9624126 | 66.3473956 | 0.68606997 |

**Figure S6f - SI By Switch category - C1 barrel**

| Kruskal Wallis | 2.05E-05 | | | | |
|---|---|---|---|---|---|
| Group A | Group B | Lower Limit | A-B | Upper Limit | P-value |
| NR to NR | NR to R | -40.148634 | -9.9574468 | 20.2337406 | 0.83178006 |
| NR to NR | R to NR | -47.387634 | 0.23076923 | 47.8491726 | 1.00E+00 |
| NR to NR | R to R | 11.6166689 | 37.5 | 63.3833311 | 1.13E-03 |
| NR to R | R to NR | -38.729564 | 10.188216 | 59.105996 | 0.95052102 |
| NR to R | R to R | 19.2548994 | 47.4574468 | 75.6599942 | 9.07E-05 |
| R to NR | R to R | -9.1138289 | 37.2692308 | 83.6522904 | 0.164927 |

**Figure S6g - PCA of baseline NR switching behavior**

```
pca_data=[double(pca_dataset.Barrel_imaged),pca_dataset.MI_C1stim,pca_dataset.MI_D1stim,pca_dataset.SI_MI_BL,pca_dataset.Spont_mean_Z];
```

| PCA Coefficients | PC1 | PC2 | PC3 | PC4 | PC5 |
|---|---|---|---|---|---|
| Barrel imaged | -0.0283 | 0.0079 | 0.088 | -0.6205 | 0.7787 |
| MI C1stim | 0.9979 | -0.0119 | -0.0088 | -0.0625 | -0.0125 |
| MI D1 stim | 0.0296 | -0.0293 | 0.9361 | 0.3164 | 0.1477 |
| SI | 0.049 | -0.0293 | -0.34 | 0.714 | 0.6094 |
| Spont mean Z | 0.0144 | 0.999 | 0.0167 | 0.0344 | 0.0159 |

| Variance explained | |
|---|---|
| PC1 | 88.8005 |
| PC2 | 8.6403 |
| PC3 | 1.5407 |
| PC4 | 0.7713 |
| PC5 | 0.2472 |

**Figure S6h - GLME of baseline NR switching behavior**

Model information:

| | |
|---|---|
| Number of observations | 159 |
| Fixed effects coefficients | 6 |
| Random effects coefficients | 11 |
| Covariance parameters | 1 |
| Distribution | Binomial |
| Link | Logit |
| FitMethod | MPL |

Formula:

Switch_BL_to_D12 ~ 1 + Barrel_imaged + MI_C1stim + MI_D1stim + SI_MI_BL + Spont_mean_Z + (1 | MouseID)

Model fit statistics:

| AIC | BIC | LogLikelihood | Deviance | AUC | | |
|---|---|---|---|---|---|---|
| 733.15 | 754.63 | -359.58 | 719.15 | 0.7679 | 0.6895 | 0.8394 |

| Name | Estimate | SE | tStat | DF | pValue | Lower | Upper |
|---|---|---|---|---|---|---|---|
| {'(Intercept)' } | 1.3474 | 0.53486 | 2.5192 | 153 | 0.012791 | 0.29073 | 2.4041 |
| {'Barrel_imaged_D1'} | -0.13854 | 0.5386 | -0.25722 | 153 | 0.79735 | -1.2026 | 0.92552 |
| {'MI_C1stim' } | -0.041889 | 0.039036 | -1.0731 | 153 | 0.28492 | -0.11901 | 0.03523 |
| {'MI_D1stim' } | -1.2086 | 0.32623 | -3.7046 | 153 | 0.00029523 | -1.8531 | -0.56406 |
| {'SI_MI_BL' } | -0.55405 | 0.49595 | -1.1172 | 153 | 0.26568 | -1.5338 | 0.42574 |
| {'Spont_mean_Z' } | 0.15878 | 0.11866 | 1.3381 | 153 | 0.18285 | -0.075644 | 0.3932 |

**Figure S7c - Change in TCW BL vs D46 - C1 Barrel**

Linear mixed-effects model fit by ML

Model information:

| | |
|---|---|
| Number of observations | 200 |
| Fixed effects coefficients | 2 |
| Random effects coefficients | 200 |
| Covariance parameters | 4 |

Formula:

TCW ~ 1 + Timepoint + (1 + Timepoint | MouseID:Index)

Model fit statistics:

| AIC | BIC | LogLikelihood | Deviance |
|---|---|---|---|
| 647.08 | 666.87 | -317.54 | 635.08 |

| Name | Estimate | SE | tStat | DF | pValue | Lower | Upper |
|---|---|---|---|---|---|---|---|
| {'(Intercept)' | 1.2454 | 0.098868 | 12.597 | 198 | 4.17E-27 | 1.0505 | 1.4404 |
| {'Timepoint_ | 0.38073 | 0.16887 | 2.2546 | 198 | 0.025256 | 0.047713 | 0.71375 |

**Figure S9a - hM3D(Gq) - PV Cells - Fbase_noise**
Linear mixed-effects model fit by REML

Model information:
Number of observations 184
Fixed effects coefficients 2
Random effects coefficients 92
Covariance parameters 2

Formula:
Fbase_noise ~ 1 + condition + (1 | MouseID:index)

Model fit statistics:

| AIC | BIC | LogLikelihood | Deviance |
|---|---|---|---|
| -912.1 | -899.28 | 460.05 | -920.1 |

| Name | Estimate | SE | tStat | DF | pValue | Lower | Upper |
|---|---|---|---|---|---|---|---|
| {'(Intercept)' } | 0.067026 | 0.0020002 | 33.509 | 182 | 8.99E-80 | 0.063079 | 0.070973 |
| {'condition_C21'} | -0.020783 | 0.0024285 | -8.5582 | 182 | 4.68E-15 | -0.025575 | -0.015992 |

**Figure S9b - hM3D(Gq) - PV Cells - MI (also reported as Fig. 4e)**
Linear mixed-effects model fit by REML

Model information:
Number of observations 184
Fixed effects coefficients 2
Random effects coefficients 92
Covariance parameters 2

Formula:
MI_dF ~ 1 + condition + (1 | MouseID:index)

Model fit statistics:

| AIC | BIC | LogLikelihood | Deviance |
|---|---|---|---|
| -399.87 | -387.05 | 203.93 | -407.87 |

| Name | Estimate | SE | tStat | DF | pValue | Lower | Upper |
|---|---|---|---|---|---|---|---|
| {'(Intercept)' } | 0.13664 | 0.0082346 | 16.593 | 182 | 2.92E-38 | 0.12039 | 0.15288 |
| {'condition_C21'} | -0.11042 | 0.0096522 | -11.44 | 182 | 3.52E-23 | -0.12946 | -0.091373 |

**Figure S9c - hM4D(Gi) - PV Cells - Fbase_noise**
Linear mixed-effects model fit by REML

Model information:
Number of observations 264
Fixed effects coefficients 2
Random effects coefficients 132
Covariance parameters 2

Formula:
Fbase_noise ~ 1 + condition + (1 | MouseID:index)

Model fit statistics:

| AIC | BIC | LogLikelihood | Deviance |
|---|---|---|---|
| -888.79 | -874.52 | 448.4 | -896.79 |

| Name | Estimate | SE | tStat | DF | pValue | Lower | Upper |
|---|---|---|---|---|---|---|---|
| {'(Intercept)' } | 0.067204 | 0.0037562 | 17.892 | 262 | 2.53E-47 | 0.059808 | 0.074601 |
| {'condition_C21'} | 0.039553 | 0.0048834 | 8.0995 | 262 | 2.10E-14 | 0.029937 | 0.049169 |

**Figure S9d - hM4D(Gi) - PV Cells - MI 1-3 s**
Linear mixed-effects model fit by REML

Model information:
Number of observations 264
Fixed effects coefficients 2
Random effects coefficients 132
Covariance parameters 2

Formula:
MI_dF ~ 1 + condition + (1 | MouseID:index)

Model fit statistics:

| AIC | BIC | LogLikelihood | Deviance |
|---|---|---|---|
| -335.25 | -320.98 | 171.62 | -343.25 |

| Name | Estimate | SE | tStat | DF | pValue | Lower | Upper |
|---|---|---|---|---|---|---|---|
| {'(Intercept)' } | 0.1631 | 0.012212 | 13.356 | 262 | 2.20E-31 | 0.13906 | 0.18715 |
| {'condition_C21'} | 0.0069389 | 0.010444 | 0.66442 | 262 | 5.07E-01 | -0.013625 | 0.027503 |

**Figure S9e - hM4D(Gi) - PV Cells - MI 1-2 s (also reported as Fig. 4h)**
Linear mixed-effects model fit by REML

Model information:
Number of observations 264
Fixed effects coefficients 2
Random effects coefficients 132
Covariance parameters 2

Formula:
MIearly_dF ~ 1 + condition + (1 | MouseID:index)

Model fit statistics:

| AIC | BIC | LogLikelihood | Deviance |
|---|---|---|---|
| -268.05 | -253.77 | 138.02 | -276.05 |

| Name | Estimate | SE | tStat | DF | pValue | Lower | Upper |
|---|---|---|---|---|---|---|---|
| {'(Intercept)' } | 0.1545 | 0.012677 | 12.187 | 262 | 2.29E-27 | 0.12954 | 0.17946 |
| {'condition_C21'} | 0.072483 | 0.014176 | 5.1131 | 262 | 6.12E-07 | 0.04457 | 0.1004 |

**Figure S9f - mCherry - PV Cells - Fbase_noise**
Linear mixed-effects model fit by REML

Model information:
Number of observations 142
Fixed effects coefficients 2
Random effects coefficients 71
Covariance parameters 2

Formula:
Fbase_noise ~ 1 + condition + (1 | MouseID:index)

Model fit statistics:

| AIC | BIC | LogLikelihood | Deviance |
|---|---|---|---|
| -334.06 | -322.3 | 171.03 | -342.06 |

| Name | Estimate | SE | tStat | DF | pValue | Lower | Upper |
|---|---|---|---|---|---|---|---|
| {'(Intercept)' } | 0.075894 | 0.0082486 | 9.2009 | 140 | 4.55E-16 | 0.059586 | 0.092202 |
| {'condition_C21'} | 0.021612 | 0.010843 | 1.9931 | 140 | 4.82E-02 | 0.00017369 | 0.04305 |

**Figure S9g - mCherry - PV Cells - MI (also reported as Fig. 4k)**
Linear mixed-effects model fit by REML

Model information:
Number of observations 142
Fixed effects coefficients 2
Random effects coefficients 71
Covariance parameters 2

Formula:
MI_dF ~ 1 + condition + (1 | MouseID:index)

Model fit statistics:

| AIC | BIC | LogLikelihood | Deviance |
|---|---|---|---|
| -134.77 | -123.01 | 71.387 | -142.77 |

| Name | Estimate | SE | tStat | DF | pValue | Lower | Upper |
|---|---|---|---|---|---|---|---|
| {'(Intercept)' } | 0.21105 | 0.020543 | 10.274 | 140 | 8.45E-19 | 0.17044 | 0.25166 |
| {'condition_C21'} | -0.018158 | 0.014569 | -1.2463 | 140 | 2.15E-01 | -0.046962 | 0.010646 |

**Figure S9a - hM3D(Gq) - Pyr Cells - Fbase_noise**
Linear mixed-effects model fit by REML

Model information:
Number of observations 1480
Fixed effects coefficients 2
Random effects coefficients 7
Covariance parameters 2

Formula:
Fbase_noise ~ 1 + condition + (1 | MouseID)

Model fit statistics:

| AIC | BIC | LogLikelihood | Deviance |
|---|---|---|---|
| 800.73 | 821.92 | -396.36 | 792.73 |

| Name | Estimate | SE | tStat | DF | pValue | Lower | Upper |
|---|---|---|---|---|---|---|---|
| {'(Intercept)' } | 0.11844 | 0.011047 | 10.722 | 1478 | 7.00E-26 | 0.096774 | 0.14011 |
| {'condition_C21'} | -0.062391 | 0.016456 | -3.7915 | 1478 | 1.56E-04 | -0.09467 | -0.030112 |

**Figure S9b- hM3D(Gq) - Pyr Cells - MI (also reported as Fig. 4f)**
Linear mixed-effects model fit by REML

Model information:
Number of observations 1480
Fixed effects coefficients 2
Random effects coefficients 7
Covariance parameters 2

Formula:
MI_dF ~ 1 + condition + (1 | MouseID)

Model fit statistics:

| AIC | BIC | LogLikelihood | Deviance |
|---|---|---|---|
| -429.85 | -408.66 | 218.93 | -437.85 |

| Name | Estimate | SE | tStat | DF | pValue | Lower | Upper |
|---|---|---|---|---|---|---|---|
| {'(Intercept)' } | 0.092244 | 0.0072853 | 12.662 | 1478 | 5.87E-35 | 0.077953 | 0.10653 |
| {'condition_C21'} | -0.081541 | 0.010852 | -7.5138 | 1478 | 9.90E-14 | -0.10283 | -0.060253 |

**Figure S9c - hM4D(Gi) - Pyr Cells - Fbase_noise**
Linear mixed-effects model fit by REML

Model information:
Number of observations 2092
Fixed effects coefficients 2
Random effects coefficients 8
Covariance parameters 2

Formula:
Fbase_noise ~ 1 + condition + (1 | MouseID)

Model fit statistics:

| AIC | BIC | LogLikelihood | Deviance |
|---|---|---|---|
| -2236.5 | -2213.9 | 1122.3 | -2244.5 |

| Name | Estimate | SE | tStat | DF | pValue | Lower | Upper |
|---|---|---|---|---|---|---|---|
| {'(Intercept)' } | 0.082372 | 0.0088119 | 9.3478 | 2090 | 2.22E-20 | 0.065091 | 0.099653 |
| {'condition_C21'} | 0.082427 | 0.0061637 | 13.373 | 2090 | 3.39E-39 | 0.070339 | 0.094514 |

**Figure S9d - hM4D(Gi) - Pyr Cells - MI (also reported as Fig. 4i)**
Linear mixed-effects model fit by REML

Model information:
Number of observations 2092
Fixed effects coefficients 2
Random effects coefficients 8
Covariance parameters 2

Formula:
MI_dF ~ 1 + condition + (1 | MouseID)

Model fit statistics:

| AIC | BIC | LogLikelihood | Deviance |
|---|---|---|---|
| -123.33 | -100.75 | 65.666 | -131.33 |

| Name | Estimate | SE | tStat | DF | pValue | Lower | Upper |
|---|---|---|---|---|---|---|---|
| {'(Intercept)' } | 0.089146 | 0.020289 | 4.3938 | 2090 | 1.17E-05 | 0.049357 | 0.12894 |
| {'condition_C21'} | 0.073513 | 0.010208 | 7.2016 | 2090 | 8.26E-13 | 0.053495 | 0.093532 |

**Figure S9e - hM4D(Gi) - PV Cells - MI 2-3 s (also reported as Fig.4h)**
Linear mixed-effects model fit by REML

Model information:
Number of observations 264
Fixed effects coefficients 2
Random effects coefficients 132
Covariance parameters 2

Formula:
MIlate_dF ~ 1 + condition + (1 | MouseID:index)

Model fit statistics:

| AIC | BIC | LogLikelihood | Deviance |
|---|---|---|---|
| -288.02 | -273.74 | 148.01 | -296.02 |

| Name | Estimate | SE | tStat | DF | pValue | Lower | Upper |
|---|---|---|---|---|---|---|---|
| {'(Intercept)' } | 0.17171 | 0.0139 | 12.353 | 262 | 6.23E-28 | 0.14434 | 0.19908 |
| {'condition_C21'} | -0.058605 | 0.010775 | -5.4391 | 262 | 1.23E-07 | -0.079822 | -0.037389 |

**Figure S9f - mCherry - Pyr Cells - Fbase_noise**
Linear mixed-effects model fit by REML

Model information:
Number of observations 1215
Fixed effects coefficients 2
Random effects coefficients 6
Covariance parameters 2

Formula:
Fbase_noise ~ 1 + condition + (1 | MouseID)

Model fit statistics:

| AIC | BIC | LogLikelihood | Deviance |
|---|---|---|---|
| -3390.4 | -3370 | 1699.2 | -3398.4 |

| Name | Estimate | SE | tStat | DF | pValue | Lower | Upper |
|---|---|---|---|---|---|---|---|
| {'(Intercept)' } | 0.0776 | 0.0063119 | 12.294 | 1213 | 7.99E-33 | 0.065217 | 0.089983 |
| {'condition_C21'} | 0.00094461 | 0.0033941 | 0.27831 | 1213 | 0.78082 | -0.0057143 | 0.0076035 |

**Figure S9g - Figure 4l - mCherry - Pyr Cells - MI (also reported as Fig. 4l)**
Linear mixed-effects model fit by REML

Model information:
Number of observations 1215
Fixed effects coefficients 2
Random effects coefficients 6
Covariance parameters 2

Formula:
MI_dF ~ 1 + condition + (1 | MouseID)

Model fit statistics:

| AIC | BIC | LogLikelihood | Deviance |
|---|---|---|---|
| 27.756 | 48.16 | -9.8781 | 19.756 |

| Name | Estimate | SE | tStat | DF | pValue | Lower | Upper |
|---|---|---|---|---|---|---|---|
| {'(Intercept)' } | 0.093275 | 0.0099233 | 9.3996 | 1213 | 2.63E-20 | 0.073806 | 0.11274 |
| {'condition_C21'} | 0.0043966 | 0.013925 | 0.31573 | 1213 | 0.75226 | -0.022923 | 0.031717 |

**Figure S11a - Gq Spontaneous AUC - PV cells**
Linear mixed-effects model fit by REML

Model information:
Number of observations 184
Fixed effects coefficients 2
Random effects coefficients 7
Covariance parameters 2

Formula:
AUC_dF ~ 1 + condition + (1 | MouseID)

Model fit statistics:

| AIC | BIC | LogLikelihood | Deviance |
|---|---|---|---|
| 2231.5 | 2244.3 | -1111.7 | 2223.5 |

| Name | Estimate | SE | tStat | DF | pValue | Lower | Upper |
|---|---|---|---|---|---|---|---|
| {'(Intercept)' } | 100.58 | 11.065 | 9.0901 | 182 | 1.68E-16 | 78.75 | 122.41 |
| {'condition_C21'} | -22.841 | 15.648 | -1.4596 | 182 | 0.14611 | -53.716 | 8.0343 |

**Figure S11b - Gq Spontaneous Peak Freq. - PV cells**
Linear mixed-effects model fit by REML

Model information:
Number of observations 184
Fixed effects coefficients 2
Random effects coefficients 7
Covariance parameters 2

Formula:
PeakFreq_dF ~ 1 + condition + (1 | MouseID)

Model fit statistics:

| AIC | BIC | LogLikelihood | Deviance |
|---|---|---|---|
| -213.34 | -200.52 | 110.67 | -221.34 |

| Name | Estimate | SE | tStat | DF | pValue | Lower | Upper |
|---|---|---|---|---|---|---|---|
| {'(Intercept)' } | 0.16778 | 0.013397 | 12.524 | 182 | 2.36E-26 | 0.14134 | 0.19421 |
| {'condition_C21'} | 0.035195 | 0.018946 | 1.8577 | 182 | 0.064829 | -0.0021863 | 0.072577 |

**Figure S11c - Gq Spontaneous Peak Amp. - PV cells**
Linear mixed-effects model fit by REML

Model information:
Number of observations 181
Fixed effects coefficients 2
Random effects coefficients 7
Covariance parameters 2

Formula:
PeaksAmp_mean_dF ~ 1 + condition + (1 | MouseID)

Model fit statistics:

| AIC | BIC | LogLikelihood | Deviance |
|---|---|---|---|
| -259.62 | -246.87 | 133.81 | -267.62 |

| Name | Estimate | SE | tStat | DF | pValue | Lower | Upper |
|---|---|---|---|---|---|---|---|
| {'(Intercept)' } | 0.35072 | 0.011649 | 30.108 | 179 | 5.71E-72 | 0.32773 | 0.37371 |
| {'condition_C21'} | -0.11056 | 0.016612 | -6.6553 | 179 | 3.33E-10 | -0.14334 | -0.077778 |

**Figure S11d - Gi Spontaneous AUC - PV cells**
Linear mixed-effects model fit by REML

Model information:
Number of observations 264
Fixed effects coefficients 2
Random effects coefficients 8
Covariance parameters 2

Formula:
AUC_dF ~ 1 + condition + (1 | MouseID)

Model fit statistics:

| AIC | BIC | LogLikelihood | Deviance |
|---|---|---|---|
| 3228.1 | 3242.4 | -1610.1 | 3220.1 |

| Name | Estimate | SE | tStat | DF | pValue | Lower | Upper |
|---|---|---|---|---|---|---|---|
| {'(Intercept)' } | 76.006 | 24.292 | 3.1288 | 262 | 1.95E-03 | 28.172 | 123.84 |
| {'condition_C21'} | 68.612 | 13.202 | 5.197 | 262 | 4.08E-07 | 42.616 | 94.608 |

**Figure S11e - Gi Spontaneous Peak Freq. - PV cells**
Linear mixed-effects model fit by REML

Model information:
Number of observations 264
Fixed effects coefficients 2
Random effects coefficients 8
Covariance parameters 2

Formula:
PeakFreq_dF ~ 1 + condition + (1 | MouseID)

Model fit statistics:

| AIC | BIC | LogLikelihood | Deviance |
|---|---|---|---|
| -459.86 | -445.59 | 233.93 | -467.86 |

| Name | Estimate | SE | tStat | DF | pValue | Lower | Upper |
|---|---|---|---|---|---|---|---|
| {'(Intercept)' } | 0.16364 | 0.014717 | 11.119 | 262 | 8.82E-24 | 0.13466 | 0.19262 |
| {'condition_C21'} | 0.053865 | 0.011722 | 4.5952 | 262 | 6.73E-06 | 0.030784 | 0.076946 |

**Figure S11f - Gi Spontaneous Peak Amp. - PV cells**
Linear mixed-effects model fit by REML

Model information:
Number of observations 263
Fixed effects coefficients 2
Random effects coefficients 8
Covariance parameters 2

Formula:
PeaksAmp_mean_dF ~ 1 + condition + (1 | MouseID)

Model fit statistics:

| AIC | BIC | LogLikelihood | Deviance |
|---|---|---|---|
| -13.158 | 1.0996 | 10.579 | -21.158 |

| Name | Estimate | SE | tStat | DF | pValue | Lower | Upper |
|---|---|---|---|---|---|---|---|
| {'(Intercept)' } | 0.34138 | 0.060096 | 5.6805 | 261 | 3.58E-08 | 0.22304 | 0.45971 |
| {'condition_C21'} | 0.25748 | 0.02708 | 9.5082 | 261 | 1.34E-18 | 0.20416 | 0.3108 |

**Figure S11g - mCherry Spontaneous AUC - PV cells**
Linear mixed-effects model fit by REML

Model information:
Number of observations 142
Fixed effects coefficients 2
Random effects coefficients 6
Covariance parameters 2

Formula:
AUC_dF ~ 1 + condition + (1 | MouseID)

Model fit statistics:

| AIC | BIC | LogLikelihood | Deviance |
|---|---|---|---|
| 989.22 | 1001 | -490.61 | 981.22 |

| Name | Estimate | SE | tStat | DF | pValue | Lower | Upper |
|---|---|---|---|---|---|---|---|
| {'(Intercept)' } | 9.2668 | 1.5878 | 5.8362 | 140 | 3.55E-08 | 6.1276 | 12.406 |
| {'condition_C21'} | 2.517 | 1.2729 | 1.9773 | 140 | 0.04997 | 0.00033079 | 5.0336 |

**Figure S11h - mCherry Spontaneous Peak Freq. - PV cells**
Linear mixed-effects model fit by REML

Model information:
Number of observations 142
Fixed effects coefficients 2
Random effects coefficients 6
Covariance parameters 2

Formula:
PeakFreq_dF ~ 1 + condition + (1 | MouseID)

Model fit statistics:

| AIC | BIC | LogLikelihood | Deviance |
|---|---|---|---|
| -301.13 | -289.36 | 154.56 | -309.13 |

| Name | Estimate | SE | tStat | DF | pValue | Lower | Upper |
|---|---|---|---|---|---|---|---|
| {'(Intercept)' } | 0.18135 | 0.017745 | 10.219 | 140 | 1.16E-18 | 0.14626 | 0.21643 |
| {'condition_C21'} | 0.0053598 | 0.012626 | 0.42452 | 140 | 0.67184 | -0.019602 | 0.030321 |

**Figure S11i - mCherry Spontaneous Peak Amp. - PV cells**
Linear mixed-effects model fit by REML

Model information:
Number of observations 142
Fixed effects coefficients 2
Random effects coefficients 6
Covariance parameters 2

Formula:
PeaksAmp_mean_dF ~ 1 + condition + (1 | MouseID)

Model fit statistics:

| AIC | BIC | LogLikelihood | Deviance |
|---|---|---|---|
| -220.46 | -208.69 | 114.23 | -228.46 |

| Name | Estimate | SE | tStat | DF | pValue | Lower | Upper |
|---|---|---|---|---|---|---|---|
| {'(Intercept)' } | 0.35224 | 0.026429 | 13.328 | 140 | 1.11E-26 | 0.29999 | 0.40449 |
| {'condition_C21'} | 0.076348 | 0.016764 | 4.5544 | 140 | 1.13E-05 | 0.043205 | 0.10949 |

**Figure S11a - Gq Spontaneous AUC - Pyr cells**
Linear mixed-effects model fit by REML

Model information:
Number of observations 1412
Fixed effects coefficients 2
Random effects coefficients 7
Covariance parameters 2

Formula:
AUC_dF ~ 1 + condition + (1 | MouseID)

Model fit statistics:

| AIC | BIC | LogLikelihood | Deviance |
|---|---|---|---|
| 23163 | 23184 | -11577 | 23155 |

| Name | Estimate | SE | tStat | DF | pValue | Lower | Upper |
|---|---|---|---|---|---|---|---|
| {'(Intercept)' } | 192.35 | 31.864 | 6.0367 | 1410 | 2.01E-09 | 129.85 | 254.86 |
| {'condition_C21'} | -125.1 | 47.403 | -2.639 | 1410 | 8.41E-03 | -218.09 | -32.109 |

**Figure S11b - Gq Spontaneous Peak Freq. - Pyr cells**
Linear mixed-effects model fit by REML

Model information:
Number of observations 1412
Fixed effects coefficients 2
Random effects coefficients 7
Covariance parameters 2

Formula:
PeakFreq_dF ~ 1 + condition + (1 | MouseID)

Model fit statistics:

| AIC | BIC | LogLikelihood | Deviance |
|---|---|---|---|
| -2123.9 | -2102.9 | 1066 | -2131.9 |

| Name | Estimate | SE | tStat | DF | pValue | Lower | Upper |
|---|---|---|---|---|---|---|---|
| {'(Intercept)' } | 0.26358 | 0.013458 | 19.585 | 1410 | 9.81E-76 | 0.23718 | 0.28998 |
| {'condition_C21'} | -0.15454 | 0.006032 | -25.621 | 1410 | 3.52E-119 | -0.16638 | -0.14271 |

**Figure S11c - Gq Spontaneous Peak Amp. - Pyr cells**
Linear mixed-effects model fit by REML

Model information:
Number of observations 1364
Fixed effects coefficients 2
Random effects coefficients 7
Covariance parameters 2

Formula:
PeaksAmp_mean_dF ~ 1 + condition + (1 | MouseID)

Model fit statistics:

| AIC | BIC | LogLikelihood | Deviance |
|---|---|---|---|
| 3711.6 | 3732.5 | -1851.8 | 3703.6 |

| Name | Estimate | SE | tStat | DF | pValue | Lower | Upper |
|---|---|---|---|---|---|---|---|
| {'(Intercept)' } | 0.83238 | 0.033756 | 24.658 | 1362 | 2.68E-111 | 0.76616 | 0.8986 |
| {'condition_C21'} | -0.51288 | 0.051239 | -10.01 | 1362 | 8.31E-23 | -0.6134 | -0.41236 |

**Figure S11d - Gi Spontaneous AUC - Pyr cells**
Linear mixed-effects model fit by REML

Model information:
Number of observations 1927
Fixed effects coefficients 2
Random effects coefficients 8
Covariance parameters 2

Formula:
AUC_dF ~ 1 + condition + (1 | MouseID)

Model fit statistics:

| AIC | BIC | LogLikelihood | Deviance |
|---|---|---|---|
| 25213 | 25235 | -12602 | 25205 |

| Name | Estimate | SE | tStat | DF | pValue | Lower | Upper |
|---|---|---|---|---|---|---|---|
| {'(Intercept)' } | 95.757 | 31.326 | 3.0568 | 1925 | 2.27E-03 | 34.32 | 157.19 |
| {'condition_C21'} | 97.612 | 7.6514 | 12.757 | 1925 | 7.67E-36 | 82.606 | 112.62 |

**Figure S11e - Gi Spontaneous Peak Freq. - Pyr cells**
Linear mixed-effects model fit by REML

Model information:
Number of observations 1927
Fixed effects coefficients 2
Random effects coefficients 8
Covariance parameters 2

Formula:
PeakFreq_dF ~ 1 + condition + (1 | MouseID)

Model fit statistics:

| AIC | BIC | LogLikelihood | Deviance |
|---|---|---|---|
| -2927.4 | -2905.1 | 1467.7 | -2935.4 |

| Name | Estimate | SE | tStat | DF | pValue | Lower | Upper |
|---|---|---|---|---|---|---|---|
| {'(Intercept)' } | 0.19779 | 0.0198 | 9.989 | 1925 | 6.09E-23 | 0.15896 | 0.23662 |
| {'condition_C21'} | 0.024724 | 0.0051232 | 4.8259 | 1925 | 1.50E-06 | 0.014677 | 0.034772 |

**Figure S11f - Gi Spontaneous Peak Amp. - Pyr cells**
Linear mixed-effects model fit by REML

Model information:
Number of observations 1917
Fixed effects coefficients 2
Random effects coefficients 8
Covariance parameters 2

Formula:
PeaksAmp_mean_dF ~ 1 + condition + (1 | MouseID)

Model fit statistics:

| AIC | BIC | LogLikelihood | Deviance |
|---|---|---|---|
| 2844.4 | 2866.6 | -1418.2 | 2836.4 |

| Name | Estimate | SE | tStat | DF | pValue | Lower | Upper |
|---|---|---|---|---|---|---|---|
| {'(Intercept)' } | 0.53743 | 0.082383 | 6.5235 | 1915 | 8.76E-11 | 0.37586 | 0.699 |
| {'condition_C21'} | 0.40168 | 0.0231 | 17.389 | 1915 | 5.35E-63 | 0.35638 | 0.44699 |

**Figure S11g - mCherry Spontaneous AUC - Pyr cells**
Linear mixed-effects model fit by REML

Model information:
Number of observations 1269
Fixed effects coefficients 2
Random effects coefficients 6
Covariance parameters 2

Formula:
AUC_dF ~ 1 + condition + (1 | MouseID)

Model fit statistics:

| AIC | BIC | LogLikelihood | Deviance |
|---|---|---|---|
| 11038 | 11059 | -5515.2 | 11030 |

| Name | Estimate | SE | tStat | DF | pValue | Lower | Upper |
|---|---|---|---|---|---|---|---|
| {'(Intercept)' } | 11.195 | 2.5982 | 4.3089 | 1267 | 1.77E-05 | 6.098 | 16.292 |
| {'condition_C21'} | 2.2615 | 1.0474 | 2.1592 | 1267 | 3.10E-02 | 0.20671 | 4.3162 |

**Figure S11h - mCherry Spontaneous Peak Freq. - Pyr cells**
Linear mixed-effects model fit by REML

Model information:
Number of observations 1269
Fixed effects coefficients 2
Random effects coefficients 6
Covariance parameters 2

Formula:
PeakFreq_dF ~ 1 + condition + (1 | MouseID)

Model fit statistics:

| AIC | BIC | LogLikelihood | Deviance |
|---|---|---|---|
| -1943.5 | -1922.9 | 975.74 | -1951.5 |

| Name | Estimate | SE | tStat | DF | pValue | Lower | Upper |
|---|---|---|---|---|---|---|---|
| {'(Intercept)' } | 0.19474 | 0.028033 | 6.947 | 1267 | 5.95E-12 | 0.13975 | 0.24974 |
| {'condition_C21'} | 0.019065 | 0.0062251 | 3.0627 | 1267 | 0.0022399 | 0.0068528 | 0.031278 |

**Figure S11i - mCherry Spontaneous Peak Amp. - Pyr cells**
Linear mixed-effects model fit by REML

Model information:
Number of observations 1267
Fixed effects coefficients 2
Random effects coefficients 6
Covariance parameters 2

Formula:
PeaksAmp_mean_dF ~ 1 + condition + (1 | MouseID)

Model fit statistics:

| AIC | BIC | LogLikelihood | Deviance |
|---|---|---|---|
| 1676.5 | 1697 | -834.23 | 1668.5 |

| Name | Estimate | SE | tStat | DF | pValue | Lower | Upper |
|---|---|---|---|---|---|---|---|
| {'(Intercept)' } | 0.6178 | 0.064043 | 9.6467 | 1265 | 2.69E-21 | 0.49216 | 0.74344 |
| {'condition_C21'} | 0.020439 | 0.026092 | 0.78336 | 1265 | 0.43356 | -0.030749 | 0.071627 |

**Figure S13a - Gq - Spont. AUC - SST cells**
Linear mixed-effects model fit by REML

Model information:

| | |
|---|---|
| Number of observations | 274 |
| Fixed effects coefficients | 2 |
| Random effects coefficients | 137 |
| Covariance parameters | 2 |

Formula:
AUC_dF ~ 1 + condition + (1 | MouseID:index)

Model fit statistics:

| AIC | BIC | LogLikelihood | Deviance |
|---|---|---|---|
| 1344.6 | 1359 | -668.31 | 1336.6 |

| Name | Estimate | SE | tStat | DF | pValue | Lower | Upper |
|---|---|---|---|---|---|---|---|
| {'(Intercept)' } | 1.6606 | 0.24076 | 6.8973 | 272 | 3.71E-11 | 1.1866 | 2.1346 |
| {'condition_C21'} | -1.0159 | 0.29502 | -3.4437 | 272 | 0.00066426 | -1.5968 | -0.43515 |

**Figure S13b - Gq - Spont. Peak Freq. - SST cells**
Linear mixed-effects model fit by REML

Model information:

| | |
|---|---|
| Number of observations | 274 |
| Fixed effects coefficients | 2 |
| Random effects coefficients | 137 |
| Covariance parameters | 2 |

Formula:
PeakFreq_dF ~ 1 + condition + (1 | MouseID:index)

Model fit statistics:

| AIC | BIC | LogLikelihood | Deviance |
|---|---|---|---|
| -491.79 | -477.36 | 249.89 | -499.79 |

| Name | Estimate | SE | tStat | DF | pValue | Lower | Upper |
|---|---|---|---|---|---|---|---|
| {'(Intercept)' } | 0.10628 | 0.0085105 | 12.488 | 272 | 1.35E-28 | 0.089529 | 0.12304 |
| {'condition_C21'} | -0.033137 | 0.0091423 | -3.6246 | 272 | 0.00034531 | -0.051136 | -0.015139 |

**Figure S13c - Gq - Spont. Peak Amp. - SST cells**
Linear mixed-effects model fit by REML

Model information:

| | |
|---|---|
| Number of observations | 247 |
| Fixed effects coefficients | 2 |
| Random effects coefficients | 137 |
| Covariance parameters | 2 |

Formula:
PeaksAmp_mean_dF ~ 1 + condition + (1 | MouseID:index)

Model fit statistics:

| AIC | BIC | LogLikelihood | Deviance |
|---|---|---|---|
| -720.4 | -706.4 | 364.2 | -728.4 |

| Name | Estimate | SE | tStat | DF | pValue | Lower | Upper |
|---|---|---|---|---|---|---|---|
| {'(Intercept)' } | 0.078686 | 0.0050212 | 15.671 | 245 | 8.28E-39 | 0.068795 | 0.088576 |
| {'condition_C21'} | -0.024562 | 0.0053602 | -4.5823 | 245 | 7.33E-06 | -0.03512 | -0.014004 |

**Figure S13d - Gi - Spont. AUC - SST cells**
Linear mixed-effects model fit by REML

Model information:

| | |
|---|---|
| Number of observations | 346 |
| Fixed effects coefficients | 2 |
| Random effects coefficients | 173 |
| Covariance parameters | 2 |

Formula:
AUC_dF ~ 1 + condition + (1 | MouseID:index)

Model fit statistics:

| AIC | BIC | LogLikelihood | Deviance |
|---|---|---|---|
| 1761.9 | 1777.3 | -876.96 | 1753.9 |

| Name | Estimate | SE | tStat | DF | pValue | Lower | Upper |
|---|---|---|---|---|---|---|---|
| {'(Intercept)' } | 0.733 | 0.24442 | 2.9989 | 344 | 2.91E-03 | 0.25224 | 1.2137 |
| {'condition_C21'} | 1.4844 | 0.25986 | 5.7121 | 344 | 2.42E-08 | 0.97325 | 1.9955 |

**Figure S13e - Gi - Spont. Peak Freq. - SST cells**
Linear mixed-effects model fit by REML

Model information:

| | |
|---|---|
| Number of observations | 346 |
| Fixed effects coefficients | 2 |
| Random effects coefficients | 173 |
| Covariance parameters | 2 |

Formula:
PeakFreq_dF ~ 1 + condition + (1 | MouseID:index)

Model fit statistics:

| AIC | BIC | LogLikelihood | Deviance |
|---|---|---|---|
| -591.58 | -576.21 | 299.79 | -599.58 |

| Name | Estimate | SE | tStat | DF | pValue | Lower | Upper |
|---|---|---|---|---|---|---|---|
| {'(Intercept)' } | 0.063328 | 0.0076377 | 8.2915 | 344 | 2.55E-15 | 0.048305 | 0.07835 |
| {'condition_C21'} | 0.10632 | 0.0098372 | 10.808 | 344 | 1.23E-23 | 0.086969 | 0.12567 |

**Figure S13f - Gi - Spont. Peak Amp. - SST cells**
Linear mixed-effects model fit by REML

Model information:

| | |
|---|---|
| Number of observations | 331 |
| Fixed effects coefficients | 2 |
| Random effects coefficients | 173 |
| Covariance parameters | 2 |

Formula:
PeaksAmp_mean_dF ~ 1 + condition + (1 | MouseID:index)

Model fit statistics:

| AIC | BIC | LogLikelihood | Deviance |
|---|---|---|---|
| -583.98 | -568.8 | 295.99 | -591.98 |

| Name | Estimate | SE | tStat | DF | pValue | Lower | Upper |
|---|---|---|---|---|---|---|---|
| {'(Intercept)' } | 0.07817 | 0.0098074 | 7.9705 | 329 | 2.62E-14 | 0.058876 | 0.097463 |
| {'condition_C21'} | 0.027249 | 0.005956 | 4.575 | 329 | 6.76E-06 | 0.015532 | 0.038966 |

**Figure S13g - mCherry - Spont. AUC - SST cells**
Linear mixed-effects model fit by REML

Model information:

| | |
|---|---|
| Number of observations | 160 |
| Fixed effects coefficients | 2 |
| Random effects coefficients | 80 |
| Covariance parameters | 2 |

Formula:
AUC_dF ~ 1 + condition + (1 | MouseID:index)

Model fit statistics:

| AIC | BIC | LogLikelihood | Deviance |
|---|---|---|---|
| 740.31 | 752.56 | -366.16 | 732.31 |

| Name | Estimate | SE | tStat | DF | pValue | Lower | Upper |
|---|---|---|---|---|---|---|---|
| {'(Intercept)' } | 0.87584 | 0.30052 | 2.9144 | 158 | 4.08E-03 | 0.28227 | 1.4694 |
| {'condition_C21'} | 0.26678 | 0.26425 | 1.0096 | 158 | 0.31425 | -0.25514 | 0.78871 |

**Figure S13h - mCherry - Spont. Peak Freq. - SST cells**
Linear mixed-effects model fit by REML

Model information:

| | |
|---|---|
| Number of observations | 160 |
| Fixed effects coefficients | 2 |
| Random effects coefficients | 80 |
| Covariance parameters | 2 |

Formula:
PeakFreq_dF ~ 1 + condition + (1 | MouseID:index)

Model fit statistics:

| AIC | BIC | LogLikelihood | Deviance |
|---|---|---|---|
| -239.14 | -226.89 | 123.57 | -247.14 |

| Name | Estimate | SE | tStat | DF | pValue | Lower | Upper |
|---|---|---|---|---|---|---|---|
| {'(Intercept)' } | 0.083286 | 0.013167 | 6.3252 | 158 | 2.48E-09 | 0.057279 | 0.10929 |
| {'condition_C21'} | 0.025286 | 0.012502 | 2.0225 | 158 | 0.044809 | 0.00059298 | 0.049979 |

**Figure S13i - mCherry - Spont. Peak Amp. - SST cells**
Linear mixed-effects model fit by REML

Model information:

| | |
|---|---|
| Number of observations | 149 |
| Fixed effects coefficients | 2 |
| Random effects coefficients | 80 |
| Covariance parameters | 2 |

Formula:
PeaksAmp_mean_dF ~ 1 + condition + (1 | MouseID:index)

Model fit statistics:

| AIC | BIC | LogLikelihood | Deviance |
|---|---|---|---|
| -397.24 | -385.28 | 202.62 | -405.24 |

| Name | Estimate | SE | tStat | DF | pValue | Lower | Upper |
|---|---|---|---|---|---|---|---|
| {'(Intercept)' } | 0.062264 | 0.007608 | 8.184 | 147 | 1.20E-13 | 0.047229 | 0.077299 |
| {'condition_C21'} | 0.0056466 | 0.0069401 | 0.81361 | 147 | 0.41718 | -0.0080687 | 0.019362 |
